# Genetic modifiers of *Cep290*-mediated retinal degeneration

**DOI:** 10.1101/2022.01.26.477596

**Authors:** K.J. Meyer, S.S. Whitmore, E.R. Burnight, M. Riker, H.E. Mercer, A. Ajose, G.R. Howell, E.M. Stone, R.F. Mullins, B.A. Tucker, T.E. Scheetz, M.G. Anderson

## Abstract

Mutations in *CEP290* cause up to 30% of cases of Leber congenital amaurosis (LCA), a severe childhood blindness resulting from abnormalities in the photoreceptor connecting cilia that lead to rapid retinal degeneration. Like many genetic diseases, CEP290-LCA has considerable variable expressivity, indicating the presence of other factors that influence phenotypic outcome. Here, we have undertaken a phenotype-driven approach in mice to identify genetic modifiers of CEP290-mediated retinal degeneration through backcrosses and intercrosses between BXD24-*Cep290^rd16^* mice and the genetically distinct inbred CAST strain to introduce genetic variation. Optical coherence tomography was used to quantitatively measure retinal thickness as a surrogate indication of photoreceptor degeneration in the resulting *rd16*-mutant mice. It was readily apparent that CEP290-mediated retinal degeneration in the resulting mice is sensitive to genetic background, with some mice exhibiting relatively thick laminated retinas and others having thin retinas with advanced disease. Quantitative trait locus (QTL) analysis identified multiple genomic loci capable of influencing the retinal degeneration phenotype in *Cep290*-mutant mice that together account for 71.7% of the phenotypic variation in retinal thickness observed in our population. Following the QTL analysis, two suppressor loci were studied in detail through a combination of physical and molecular approaches to narrow the critical region for each QTL and identify the probable causative genetic variations.

## INTRODUCTION

Leber congenital amaurosis (LCA) type 10 is an early onset form of retinal degeneration that falls into a broad spectrum of ciliopathies caused by mutations in the *CEP290* gene. This disease (hereafter referred to as “CEP290-LCA”) has two unfortunate distinctions: LCA is considered the most severe form of inherited retinal disease, typically causing severe vision impairment in infants; and mutations in *CEP290* are the most identified cause of LCA (15–30% of cases in Caucasians)^1–3^. To gain a better understanding of CEP290-LCA, many aspects of CEP290 cell biology have been extensively studied. Combined, these studies suggest that CEP290 influences both ciliogenesis^4,5^ and ciliary protein content by acting as a gatekeeper in protein trafficking through the connecting cilium that normally bridges the inner and outer segment of photoreceptors^6–8^. CEP290 mutations can preclude the ability for normal outer segments to form and cause an accumulation of proteins typically in the outer segment, such as rhodopsin, within the inner segment^6^. A pro-apoptotic unfolded protein response associated with ER-stress ensues^9^, leading to the degeneration of rod photoreceptors characteristic of LCA^10,11^. Despite this progress, many aspects of CEP290-LCA remain enigmatic and point to a framework for understanding CEP290 that is likely still missing several components.

One example of an area of CEP290 biology that is not well understood pertains to phenotypic variability. The list of disease-associated *CEP290* mutations is relatively large (>112) and recalcitrant to predictable genotype:phenotype correlations^12^. In addition to causing LCA, *CEP290* mutations can variously cause kidney disease (nephronophthisis, Senior-Loken syndrome), neurological disease (Joubert syndrome, Joubert syndrome related disorders), and multiorgan diseases (Bardet-Biedl syndrome, Meckel-Gruber syndrome, cerebello-oculo-renal syndrome). Understanding this pleiotropy has been a longstanding challenge^5,12,13^. Retinopathy is typically present to some degree with all *CEP290* mutations^14, 15,16^, suggesting that photoreceptors may be particularly sensitive to CEP290 function and exhibit phenotypes for even hypomorphic alleles. Interestingly, CEP290-LCA mutations don’t cluster according to a specific protein function, but instead seem to frequently involve a specific kind of mutation associated with atypical splicing. The most common mutation (c.2991+1655A>G) is an intronic variant that introduces a cryptic exon and premature stop codon^12,13,17,18^; many other mutations tend to cluster in exons prone to basal exon skipping^19–22^. Thus, CEP290-LCA mutations likely give rise to near full-length proteins whose expression levels may be predictive of disease severity. However, *CEP290* mutations that are severe in one family are sometimes mild in another^23^, and even intrafamilial variability has been observed for CEP290- and other forms of LCA^24–26^. Genetic modifiers are often evoked to explain these types of variability, as well as the overall pleiotropy of LCA^2,3,12^. To date, studies testing this hypothesis directly have largely relied on candidate-driven approaches that typically test interactions between genes with overt disease-causing influences^27–32^, while agnostic genome-wide searches for modifiers with potentially less overt autonomous phenotypes have been uncommon^33^. Studies of genetic modifiers are a classic approach for gaining new insight into the basic biology of gene function, and have been successful in a broad range of organisms, from flies^34,35^ to mice^36,37^ and humans^38,39^.

Here, we have undertaken a phenotype-driven approach in mice to identify genetic modifiers of CEP290-mediated retinal degeneration through backcrosses and intercrosses between BXD24-*Cep290^rd16^* mice *(“‘rd16’)* and the genetically distinct inbred CAST strain to introduce genetic variation. The recessive *rd16* mutation results from an in-frame deletion of exons 35-39, removing a functional domain that allows CEP290 to bind microtubules^5^. As in human CEP290-LCA patients, homozygosity for the mutation leads to early onset retinal degeneration characterized by progressive retinal thinning due to degeneration of photoreceptor outer segments. The CAST strain is a wild-derived inbred strain of mice lacking overt retinal disease. After performing the crosses to collect a large cohort of *rd16* homozygotes, we used optical coherence tomography (OCT) to quantitatively measure retinal thickness as a surrogate indication of photoreceptor degeneration. It was readily apparent that CEP290-mediated retinal degeneration in the resulting mice is sensitive to genetic background, with some mice exhibiting relatively thick laminated retinas and others having thin retinas with advanced disease. QTL analysis identified multiple genomic loci capable of influencing the retinal degeneration phenotype in *Cep290*-mutant mice that together account for 71.7% of the phenotypic variation in retinal thickness observed in our population. Following the QTL analysis, two suppressor loci were studied in detail through a combination of physical and molecular approaches to narrow the critical region for each QTL and identify the probable causative genetic variations.

## RESULTS

### Retinal thickness phenotypes of CAST and *BXD24-rd16* mouse strains

To determine the baseline retinal thickness for each inbred parent strain, we imaged mice at incremental time points using optical coherence tomography (OCT). Retinal thickness of CAST mice was 183.3 ± 2.1 μm, 178.6 ± 4.0 μm, 175.8 ± 3.6 μm, and 169.4 ± 1.4 μm at 4, 6, 8, and 12 weeks of age, respectively *(n* = 10 mice at each time point; Figure 1A). Retinal thickness of *BXD24-rd16* mice was 113.0 ± 3.5 μm, 96.8 ± 3.3 μm, 92.9 ± 3.6 μm, and 86.4 ± 2.3 μm at 4, 6, 8, and 12 weeks of age, respectively (*n* = 10 mice at each time point; Figure 1A). The retinal thickness of CAST mice is significantly thicker than *BXD24-rd16* mice at every age tested *(p* < 0.0001 for each comparison; two-way ANOVA with Tukey post-test; *n* = 10 mice at each time point for each strain; Figure 1A). Small standard deviations at each time point for both inbred parent strains (CAST and BXD24-*rd16*) indicate that retinal thickness is under strong genetic regulation, even in the context of retinal degeneration in *BXD24-rd16* mice (Figure 1A).

**Figure 1.**
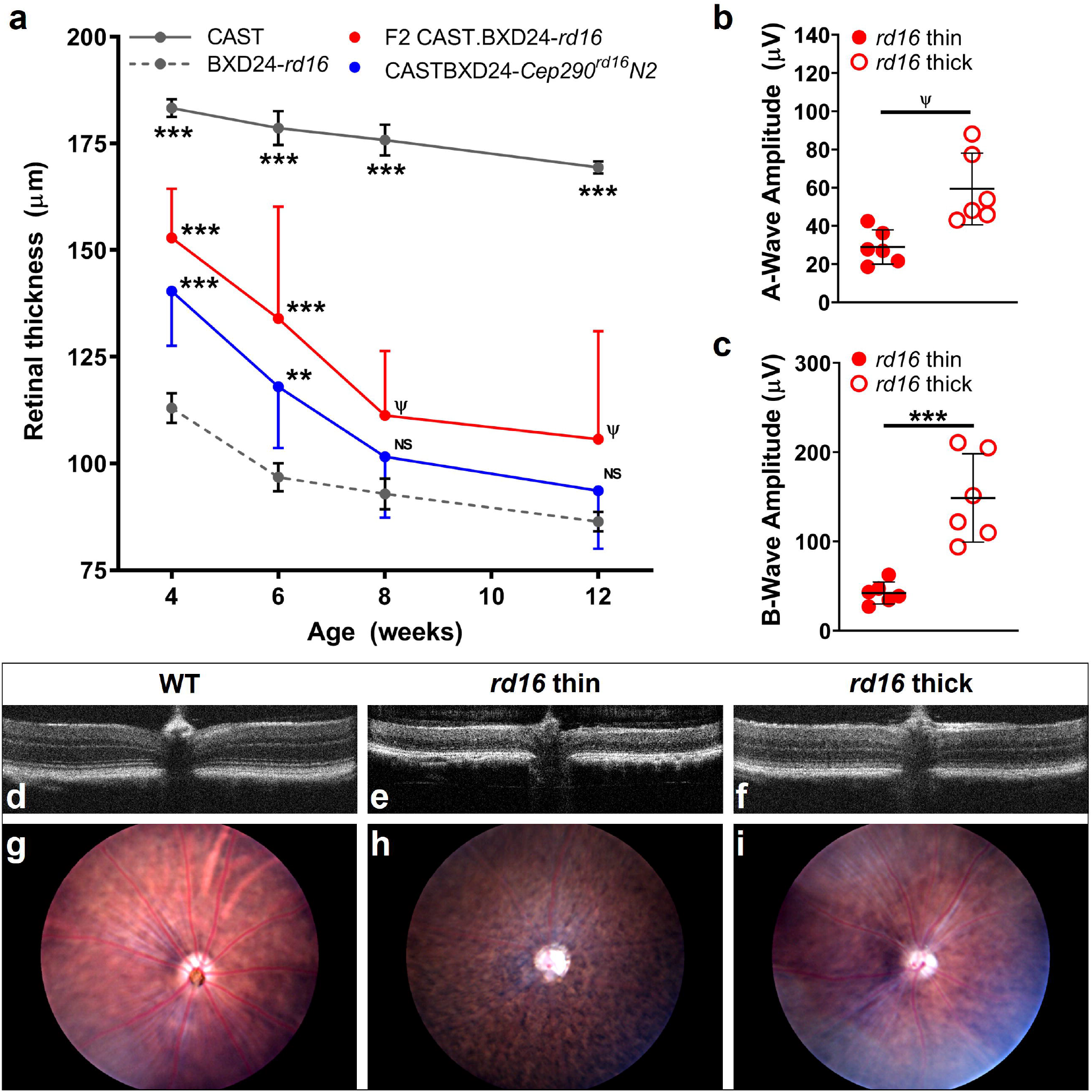
The sensitivity of *Cep290*-mediated retinal degeneration to genetic background. **a** The longitudinal retinal thickness measured using optical coherence tomography (OCT) in various genetic contexts (*error bars* = mean ± SD; N2 and F2 datasets display only half of an error bar for clarity; *n* = 120 mice, 10 per cohort per timepoint; *** = *p* < 0.001; ** = *p* < 0.05; Ψ = trending *p* < 0.10; NS = not significant). Note the differences in error bar length for inbred CAST and BXD24-*rd16* compared to the N2 and F2 *rd16*-mutant mice that have a segregating genetic background. Graphs of electroretinogram (ERG) **b** A-wave and **c** B-wave amplitude from F2 7-week-old *rd16-*mutant littermate mice with ‘thin’ (≤115 μm) retinas and ‘thick’ (≥151 μm) retinas (*error bars* = mean ± SD; *n* = 6 eyes per cohort). Representative OCT images of F2 6-week-old littermates from **d** WT retina, **e** *rd16*-mutant ‘thin’ retina, and **f** *rd16*-mutant ‘thick’ retina. Note the differences in retinal lamination in the ‘thick’ *rd16*-mutant retina (panel f) compared to the ‘thin’ *rd16*-mutant retina (panel e). Representative fundus images of F2 6-week-old littermates from **g** WT retina, **h** *rd16*-mutant ‘thin’ retina, and **i** *rd16*-mutant ‘thick’ retina. Note the differences in retinal vasculature and pigmentation in the ‘thick’ *rd16*-mutant retina (panel i) compared to the ‘thin’ *rd16*-mutant retina (panel h).

### Retinal thickness phenotypes of N2 and F2 *rd16* mice

To assess the sensitivity of the retinal phenotype conferred by the *rd16* mutation to genetic background, we performed a backcross (N2) and intercross (F2) between BXD24-*rd16* and CAST mice and compared the retinal thickness of the *rd16* homozygotes over time. Retinal thickness of N2 *rd16* homozygotes was 140.4 ± 12.8 μm, 118.0 ± 14.4 μm, 101.6 ± 14.3 μm, and 93.6 ± 13.6 μm, at 4, 6, 8, and 12 weeks of age, respectively (*n* = 10 mice at each time point; Figure 1A). Retinal thickness of F2 *rd16* homozygotes was 152.9 ± 11.5 μm, 134.0 ± 26.2 μm, 111.3 ± 15.1 μm, and 105.7 ± 25.3 μm, at 4, 6, 8, and 12 weeks of age, respectively (*n* = 10 mice at each time point; Figure 1A). As the percentage of CAST-derived genome increases (i.e., 0% in BXD24-*rd16*, ~25% in N2 mice and ~50% in F2 mice), the retinal thinning phenotype of homozygotes becomes increasingly suppressed on average (Figure 1A).

N2 and F2 *rd16* homozygotes, which have segregating genetic backgrounds, have an increased range of retinal thickness indicated by large standard deviations (Figure 1A). Average retinal thickness of N2 *rd16* homozygotes is significantly thicker than retinal thickness of the BXD24-*rd16* parent strain at 4 and 6 weeks old (*p* = 0.0001 and *p* = 0.0204, respectively; two-way ANOVA with Tukey post-test; *n* = 10 mice for each time point for each strain; Figure 1A). Average retinal thickness of F2 *rd16* homozygotes is significantly thicker than retinal thickness of the BXD24-*rd16* parent strain at 4 and 6 weeks old (*p* < 0.0001 for each comparison; two-way ANOVA with Tukey post-test; *n* = 10 mice at each time point for each strain; Figure 1A) and trending thicker at 8 and 12 weeks old (*p* = 0.0913, *p* = 0.0582, respectively; two-way ANOVA with Tukey post-test; *n* = 10 mice at each time point for each strain; Figure 1A). These results directly demonstrate that the *rd16* mutation is sensitive to genetic background and that the CAST parent strain contains genetic modifiers that influence CEP290-mediated retinal degeneration.

Electroretinography shows that 7-week-old F2 *rd16* homozygotes with relatively thick retinas trend toward increased a-wave amplitude (*p* < 0.1, Student’s two-tailed *t*-test; n = 6 eyes in each cohort; Figure 1B) and have a significantly increased b-wave amplitude (*p* < 0.001, Student’s two-tailed *t*-test, *n* = 6 eyes in each cohort; Figure 1C) compared to F2 *rd16* homozygotes with relatively thin retinas, indicating a correlation between retinal thickness and retinal function. Retinal examination by OCT and fundus imaging demonstrates that F2 *rd16* homozygotes with relatively thick retinas at six weeks of age have qualitatively improved retinal lamination, retinal vasculature, and pigmentation (Figure 1D-I). Therefore, the retinal thickness quantitative trait is physiologically relevant in the N2 and F2 populations of *rd16*-mutant mice.

### Population phenotyping

The six-week time point was chosen for further study because of the wide range of retinal thicknesses observed in N2 and F2 *rd16* homozygotes at this age. At six weeks of age, the range of retinal thickness was 82.3–157.0 μm in the N2 (*n* = 209 mice) *rd16* population and 75.3–180.8 μm in the F2 (*n* = 186 mice) *rd16* population (Figure 2a,b). In both the N2 and F2 populations there is a trend for increased retinal thickness in males compared to females, but it is not significant (Figure 2a,b). A histogram of retinal thickness in each separate population of N2 and F2 *rd16*-mutant mice shows that the phenotype is continuously and normally distributed (*p* = 0.256 and *p* = 0.144, respectively; Shapiro-Wilk goodness-of-fit test), indicating that retinal thickness is influenced by multiple genes (Supplementary Data 1). A combined N2 and F2 dataset did not follow a normal distribution (*p* < 0.001, Shapiro-Wilk goodness-of-fit test), so a log transformation of the retinal thickness phenotype data was used to correct the distribution prior to additional data analysis (*p* = 0.128, Shapiro-Wilk goodness-of-fit test).

**Figure 2.**
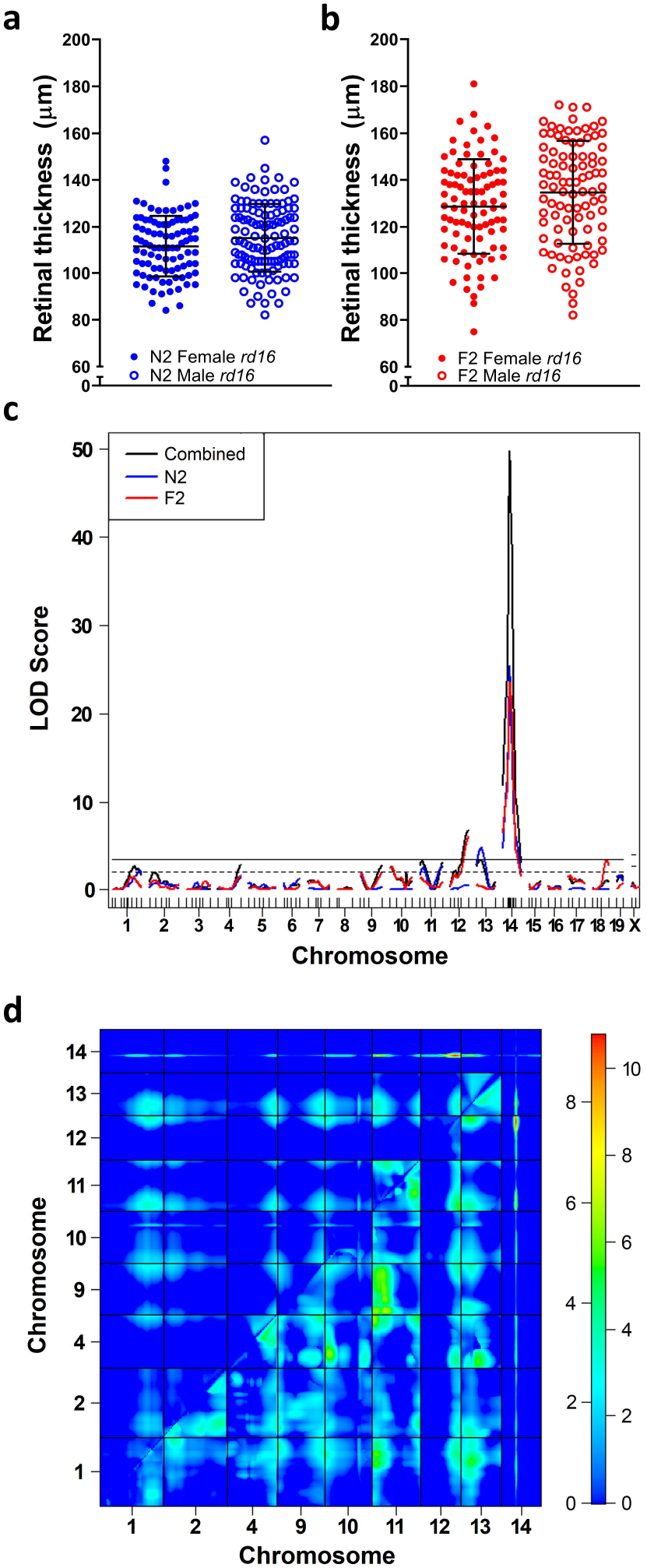
Quantitative trait locus (QTL) analysis of retinal thickness in mice with *Cep290*-mediated retinal degeneration. Retinal thickness measured by optical coherence tomography is plotted for 6-week-old (BXD24-*rd16* x CAST) **a** N2 (*n* = 209) and **b** F2 (*n* = 186) *rd16*-mutant mice. The combined group (*n* = 395) was subsequently genotyped for 90 polymorphic markers distributed across the genome and analyzed for the presence of QTL. **c** Plot shows the results of a one-dimensional genome scan to identify single QTL that modify retinal thickness in *rd16*-mutant mice. Significance thresholds determined by permutation testing are indicated by the solid (*p* < 0.05; genome-wide significant) and dashed (*p* < 0.63; genome-wide suggestive) lines. Tick marks along the bottom of the plot indicate the location of genetic markers. Five additional markers within the Chr 14 peak were genotyped in all mice. LOD = logarithm of odds. **d** Heat map shows the results of a two-dimensional, two-QTL genome scan to identify pairs of additive and/or interactive QTL capable of modifying retinal thickness in *rd16*-mutant mice. The upper left triangle displays the LOD score in which the interactive model is compared to a single QTL model (LOD_*fv1*_) and the lower right triangle displays the LOD score in which the additive model is compared to a single QTL model (LOD_*av1*_). The numbers to the left and right of the color scale bar correspond to LOD_*fv1*_ and LOD_*av1*_, respectively.

### QTL analysis: One-dimensional scan

To assess the genetic complexity of the phenotypic modification conferred by the CAST background, we performed a one-dimensional single-QTL genome scan. After genotyping 90 polymorphic genetic markers evenly dispersed throughout the genome (Supplementary Data 1) in all 395 N2 and F2 *rd16*-mutant mice having retinal thickness measurements from OCT at 6 weeks of age, we used R/qtl and permutation testing to establish significant loci from the N2, F2, and combined datasets. A one-dimensional genome-wide scan of retinal thickness using the combined dataset (209 N2 *rd16* homozygotes and 186 F2 *rd16* homozygotes) identified three loci that surpassed the genome-wide *p* = 0.05 significance threshold on Chr 12, 13, and 14 (Figure 2c, Supplementary Data 1). These loci were named *Mrdq1-3* (<u>M</u>odifier of <u>r</u>etinal <u>d</u>egeneration <u>Q</u>TL 1-3), respectively. Following initial analysis, 10 additional markers within the *Mrdq3* interval on Chr 14 were genotyped and included in all further analyses, refining the locations of *Mrdq1-3* to Chr 12 at 60.71 cM (LOD 6.72), Chr 13 at 20.3 cM (LOD 3.33), and Chr 14 at 26.83 cM (LOD 49.89), respectively. Five additional loci of interest were identified that surpassed the genome-wide *p* = 0.63 suggestive threshold: Chr 1 at 70.27 cM (LOD 2.65), Chr 4 at 82.69 cM (LOD 2.81), Chr9 at 71.33 cM (LOD 2.76), Chr 10 at 4.01 cM (LOD 2.51), and Chr 11 at 13.13 cM (LOD 3.24; Figure 2c). Independent one-dimensional genome scans of separate N2 and F2 datasets identified three unique suggestive loci in the N2 dataset on Chr 9 at 4.46 cM (LOD 1.58), Chr 11 at 75.93 cM (LOD 2.57), and Chr 19 at 20.17 cM (LOD 1.55), and one unique significant locus in the F2 dataset on Chr 18 at 49.88 cM (LOD 3.35; *Mrdq4;* Figure 2c).

### QTL analysis: Two-dimensional scan

A two-dimensional genome scan of the combined dataset under a two-QTL additive model identified loci that significantly improved the fit over the best single-QTL model on Chr 1 at 72.3 cM (Additive with *Mrdq1* and *Mrdq3*; LOD_*av1*_ = 3.38 and 4.27, respectively; *Mrdq5*), Chr 4 at 82.4 cM (Additive with *Mrdq3;* LOD_*av1*_ = 4.27; *Mrdq6*), Chr 9 at 70.5 cM (Additive with *Mrdq3;* LOD*a*w = 3.36; *Mrdq7*), Chr 10 at 16.0 cM (Additive with *Mrdq3;* LOD_*av1*_ = 3.58; *Mrdq8*), Chr 11 at 15.1 cM (Additive with *Mrdq3;* LOD_*av1*_ = 6.48; *Mrdq9),* and Chr 11 at 73.1 cM (Additive with *Mrdq9;* LOD_*av1*_ = 6.48; *Mrdq10),* that all overlap with suggestive loci identified by the one-dimensional scan (Figure 2d; Supplementary Data 1). This analysis also identified one significant novel locus on Chr 2 at 19.5 cM (Additive with *Mrdq3;* LOD_*av1*_ = 3.33; *Mrdq11;* Figure 2d; Supplementary Data 1). Significant two-QTL interactive effects that improve the fit over a single-QTL model were identified between *Mrdq1* and *Mrdq3* (LOD_*fv1*_ = 10.77) and between *Mrdq3* and *Mrdq8* (LOD_*fv1*_ = 7.61) in the combined dataset (Figure 2d; Supplementary Data 1). Independent two-dimensional genome scans of separate N2 and F2 datasets did not identify any additional unique loci (Supplementary Data 1).

### Phenotypic effects conferred by significant QTL

The chromosomal locations of *Mrdq1-11,* which account for 71.7% of the phenotypic variation within the combined population, were refined by comparing a model with and without each QTL while the positions of all other QTL are fixed (Table 1). Of the significant QTL, a CAST allele at *Mrdq1, Mrdq3, Mrdq5-8,* and *Mrdq10* suppresses retinal degeneration in a dose-dependent manner (Supplementary Fig 1a,c,e-h,j). A heterozygous genotype at *Mrdq2, Mrdq9,* and *Mrdq11* enhances retinal degeneration while homozygosity for the CAST allele at the same loci suppresses retinal degeneration (Supplementary Fig 1b,i,k). A CAST allele at *Mrdq4* suppresses retinal degeneration, with the HET genotype having the greatest suppressive effect (Supplementary Fig 1d). *Mrdq3* has the largest effect on retinal thickness, ranging from 105.3 ± 1.2 μm (mean ± SE) in mice homozygous for the BXD24 allele to 149.2 ± 1.9 μm (mean ± SE) in mice homozygous for the CAST allele, and accounting for 30.2% of the phenotypic variation within the combined population. The interactive effects between *Mrdq3* and *Mrdq8*, and between *Mrdq1* and *Mrdq3* are shown in Supplementary Fig 1l,m.

**Table 1.**
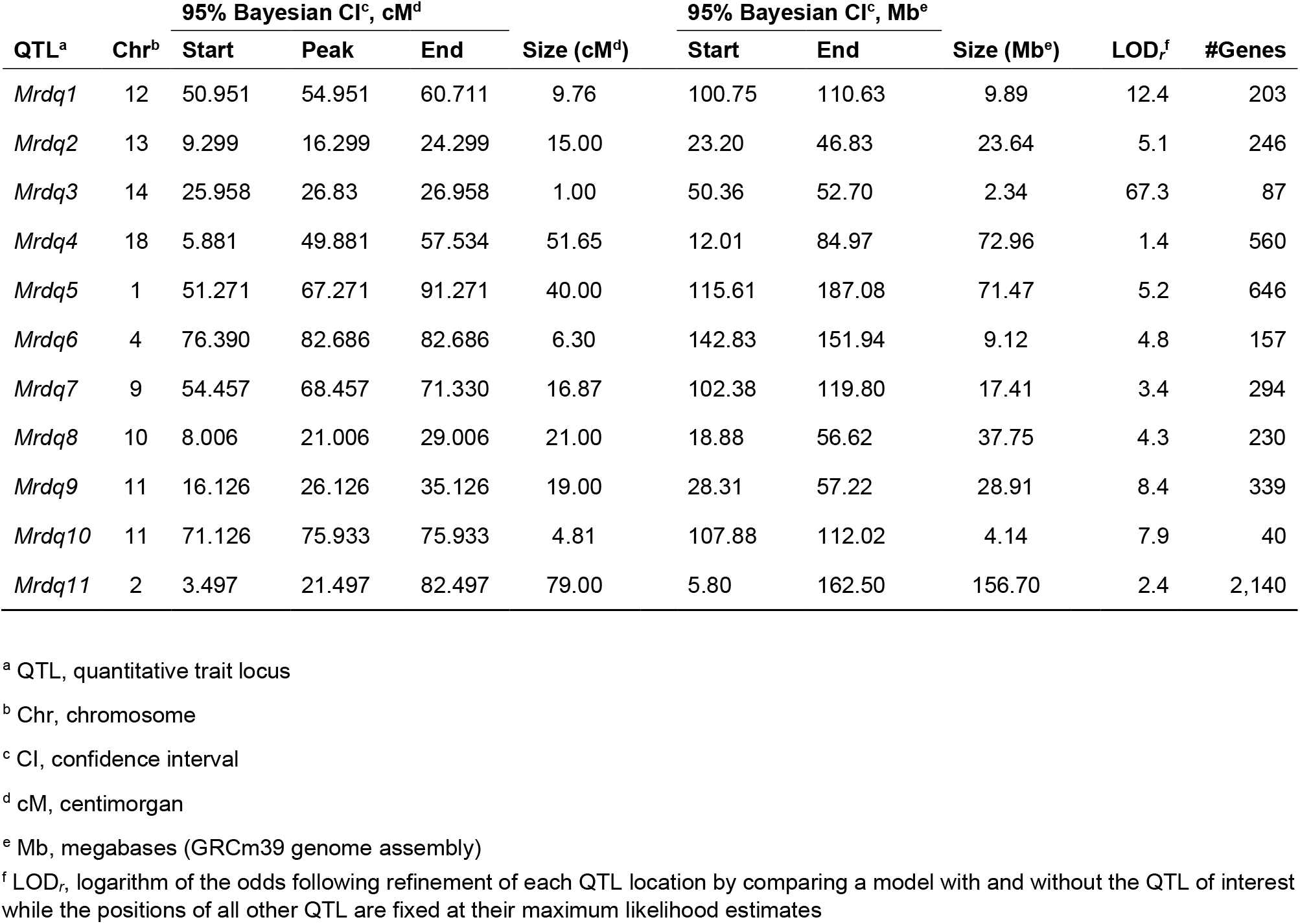
Quantitative trait loci that modify retinal thickness in *rd16*-mutant mice.

| QTL<sup>a</sup> | Chr<sup>b</sup> | 95% Bayesian CI<sup>c</sup>, cM<sup>d</sup> |  |  |  | 95% Bayesian CI<sup>c</sup>, Mb<sup>e</sup> |  |  | LOD<sub>r</sub><sup>f</sup> | #Genes |
| --- | --- | --- | --- | --- | --- | --- | --- | --- | --- | --- |
|  |  | Start | Peak | End | Size (cM<sup>d</sup>) | Start | End | Size (Mb<sup>e</sup>) |  |  |
| *Mrdq1* | 12 | 50.951 | 54.951 | 60.711 | 9.76 | 100.75 | 110.63 | 9.89 | 12.4 | 203 |
| *Mrdq2* | 13 | 9.299 | 16.299 | 24.299 | 15.00 | 23.20 | 46.83 | 23.64 | 5.1 | 246 |
| *Mrdq3* | 14 | 25.958 | 26.83 | 26.958 | 1.00 | 50.36 | 52.70 | 2.34 | 67.3 | 87 |
| *Mrdq4* | 18 | 5.881 | 49.881 | 57.534 | 51.65 | 12.01 | 84.97 | 72.96 | 1.4 | 560 |
| *Mrdq5* | 1 | 51.271 | 67.271 | 91.271 | 40.00 | 115.61 | 187.08 | 71.47 | 5.2 | 646 |
| *Mrdq6* | 4 | 76.390 | 82.686 | 82.686 | 6.30 | 142.83 | 151.94 | 9.12 | 4.8 | 157 |
| *Mrdq7* | 9 | 54.457 | 68.457 | 71.330 | 16.87 | 102.38 | 119.80 | 17.41 | 3.4 | 294 |
| *Mrdq8* | 10 | 8.006 | 21.006 | 29.006 | 21.00 | 18.88 | 56.62 | 37.75 | 4.3 | 230 |
| *Mrdq9* | 11 | 16.126 | 26.126 | 35.126 | 19.00 | 28.31 | 57.22 | 28.91 | 8.4 | 339 |
| *Mrdq10* | 11 | 71.126 | 75.933 | 75.933 | 4.81 | 107.88 | 112.02 | 4.14 | 7.9 | 40 |
| *Mrdq11* | 2 | 3.497 | 21.497 | 82.497 | 79.00 | 5.80 | 162.50 | 156.70 | 2.4 | 2,140 |

<sup>a</sup> QTL, quantitative trait locus
<sup>b</sup> Chr, chromosome
<sup>c</sup> CI, confidence interval
<sup>d</sup> cM, centimorgan
<sup>e</sup> Mb, megabases (GRCm39 genome assembly)
<sup>f</sup> $LOD_r$ , logarithm of the odds following refinement of each QTL location by comparing a model with and without the QTL of interest while the positions of all other QTL are fixed at their maximum likelihood estimates

### RNA-Seq identifies differentially expressed genes within Mrdq intervals

Due to the identification of 11 QTL capable of influencing retinal thickness in *rd16*-mutant mice as well as the large number of genetic variants that exist between the CAST and BXD24 mouse strains, RNA-Seq was used for retinal transcriptome profiling. Comparison of P25 retinas from CAST and BXD24 mice identified 3,049 differentially expressed genes (counts per million ≥ 5 and false discovery rate < 0.05) and 405 of these localize to Mrdq intervals (Supplementary Table 1). For the three most significant Mrdq intervals, 10 differentially expressed genes were identified within *Mrdq1,* 15 within *Mrdq2,* and 3 within *Mrdq3* (Table 2).

**Table 2.**
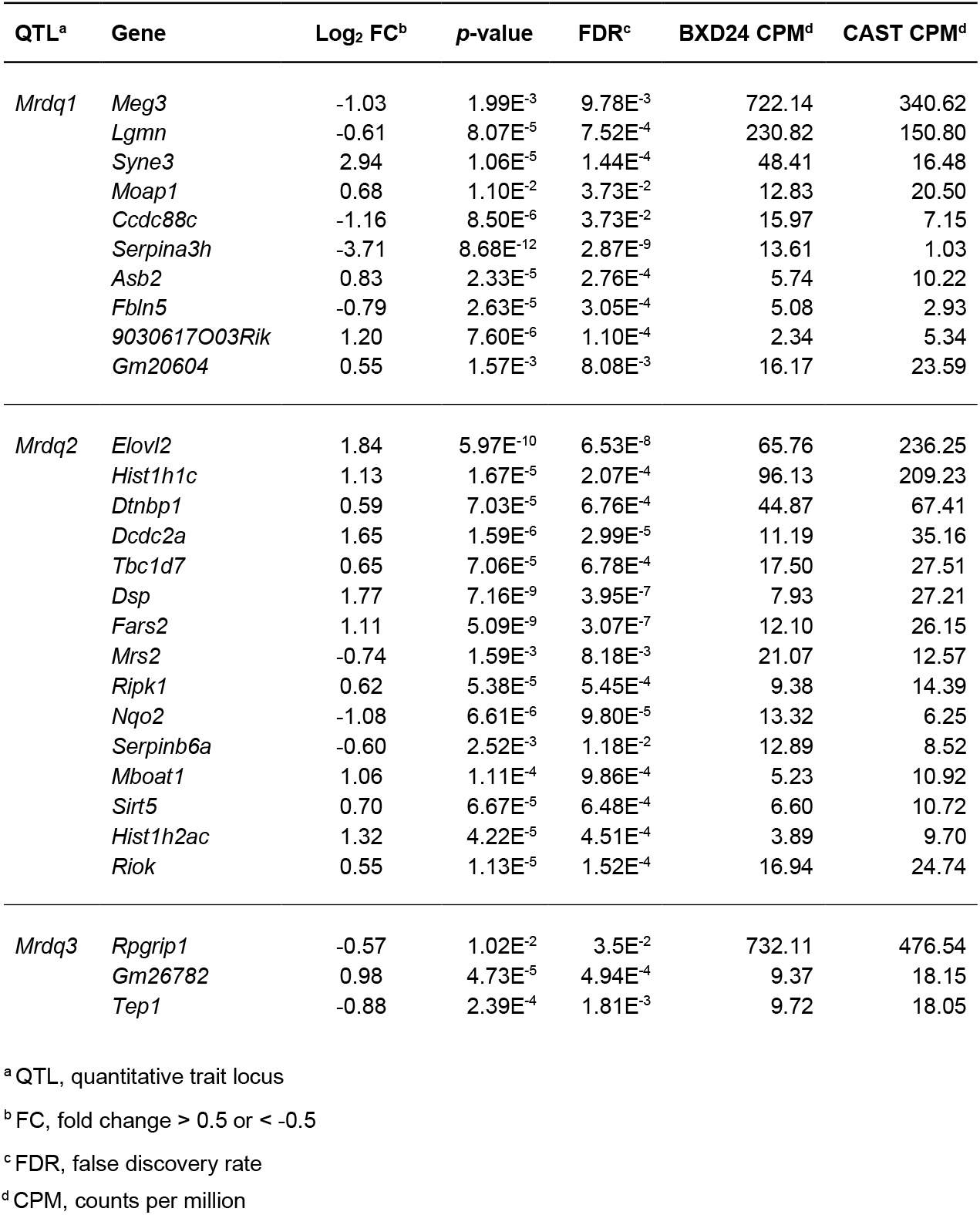
*Mrdq1, Mrdq2,* and *Mrdq3* positional candidate genes with significant expression differences in the P25 mouse retina between BXD24 and CAST.

### Recombination mapping of *Mrdq3*

The 95% Bayesian credible interval of *Mrdq3* spans a 1.0 cM (2.34 Mb) region on Chr 14 (25.958-26.958) and contains 87 genes. To reduce the genetic heterogeneity and physically narrow the *Mrdq3* interval, N2 *rd16*-mutant mice were backcrossed onto the BXD24-*rd16* background to the N10 generation. Retinal thickness was evaluated by OCT in a subset of 6-week-old *rd16*-mutant congenic mice at generations N5-N10. At each generation, the retinal thickness of mice carrying a CAST allele at *Mrdq3* was consistently and significantly increased compared to mice carrying only BXD24 alleles at *Mrdq3* (*p* < 3.1 x 10^-4^ for each comparison; two-way ANOVA with Sidak post-test; Figure 3a). The retinal thickness of both groups decreased at each subsequent generation for N5-N7, with overlapping retinal thickness phenotypes, and stabilized at generations N8-N10, with distinct non-overlapping retinal thickness phenotypes dependent on *Mrdq3* genotypes (Figure 3a). At generation N10, congenic mice were intercrossed, and the resulting *rd16*-mutant mice were screened for recombination events within the *Mrdq3* interval. Congenic *rd16*-mutant mice having intact genotypes across *Mrdq3* were phenotyped at 6-weeks-old by OCT and show a significant genotype-dependent retinal thickness with BXD24 genotypes at *Mrdq3* having the thinnest retinas, heterozygous genotypes having an intermediate retinal thickness, and CAST genotypes having the thickest retinas (95.3 ± 2.3 μm vs. 108.7 ± 3.3 μm vs. 125.7 ± 2.2 μm, respectively; *p* < 1.0 x 10^-15^ for all comparisons, one-way ANOVA with Tukey post-test; Figure 3b). *Mrdq3* recombination events were identified in 14 of the 397 N10F2 *rd16*-mutant mice screened. OCT retinal thickness phenotyping of 6-week-old *Mrdq3* recombinants was used to deduce the expected genotype at *Mrdq3* (Figure 3b). Based on recombination breakpoints, the critical region was physically narrowed to a 2.4 Mb region flanked by markers *Mrdq3-STR15* and *Mdrq3-STR12* and containing 20 genes (Figure 4). Some breakpoints were additionally supported by phenotyping progeny resulting from testcrosses.

**Figure 3.**
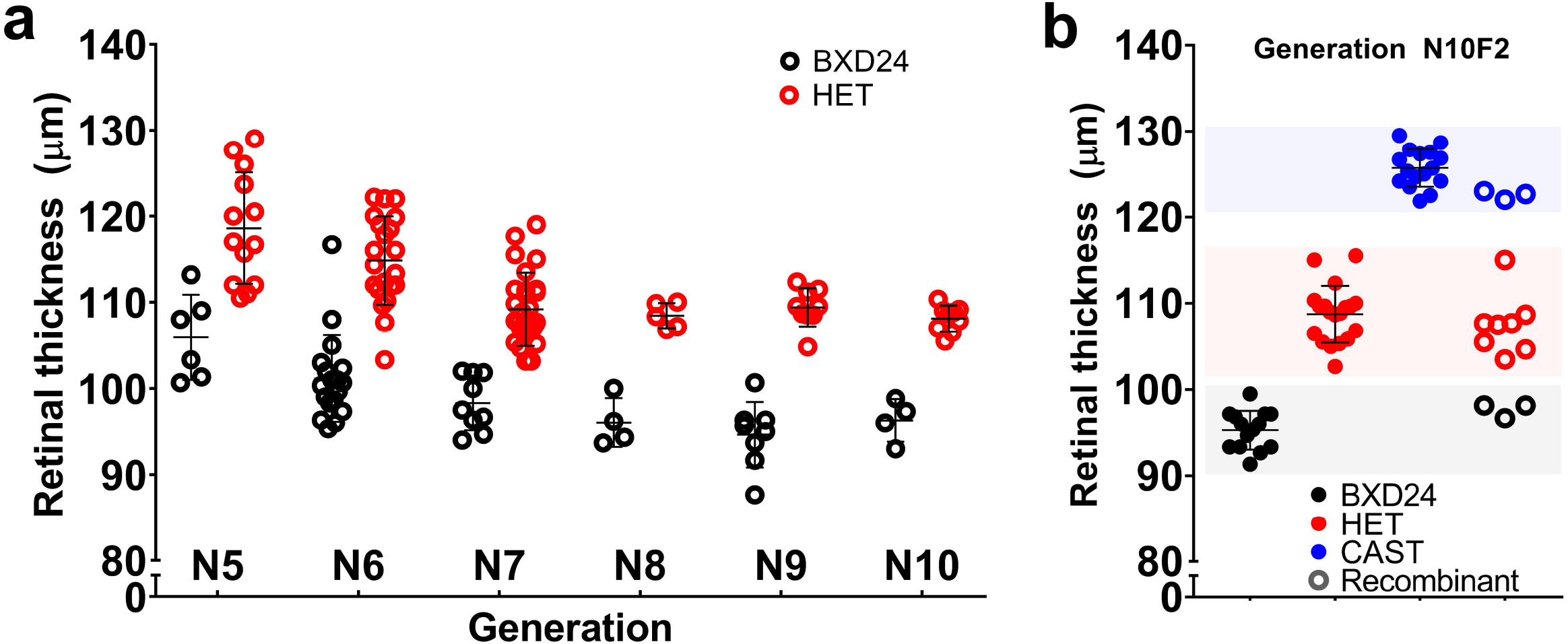
Retinal phenotypes of BXD24-*rd16* mice congenic for CAST-derived Chr14. Retinal thickness measured by optical coherence tomography OCT is plotted for 6-week-old BXD24;cg(CAST-Chr14) *rd16*-mutant mice from **a** iterative generations of backcrosses (note the separation of phenotypic overlap in generations N8-N10 between mice with BXD24 vs. heterozygous (HET) genotypes across the Chr 14 congenic interval); and a **b** N10F2 intercross. A total of 397 *rd16*-mutant mice were screened for chromosomal recombination events within the Chr 14 congenic interval. Phenotypes are shown for a subset of mice having an intact congenic interval with BXD24, HET, or CAST alleles; and littermate mice having a recombination event (*n* = 14; *error bars* = mean ± SD). Shading indicates the discreet phenotypic range for each congenic interval genotype.

**Figure 4.**
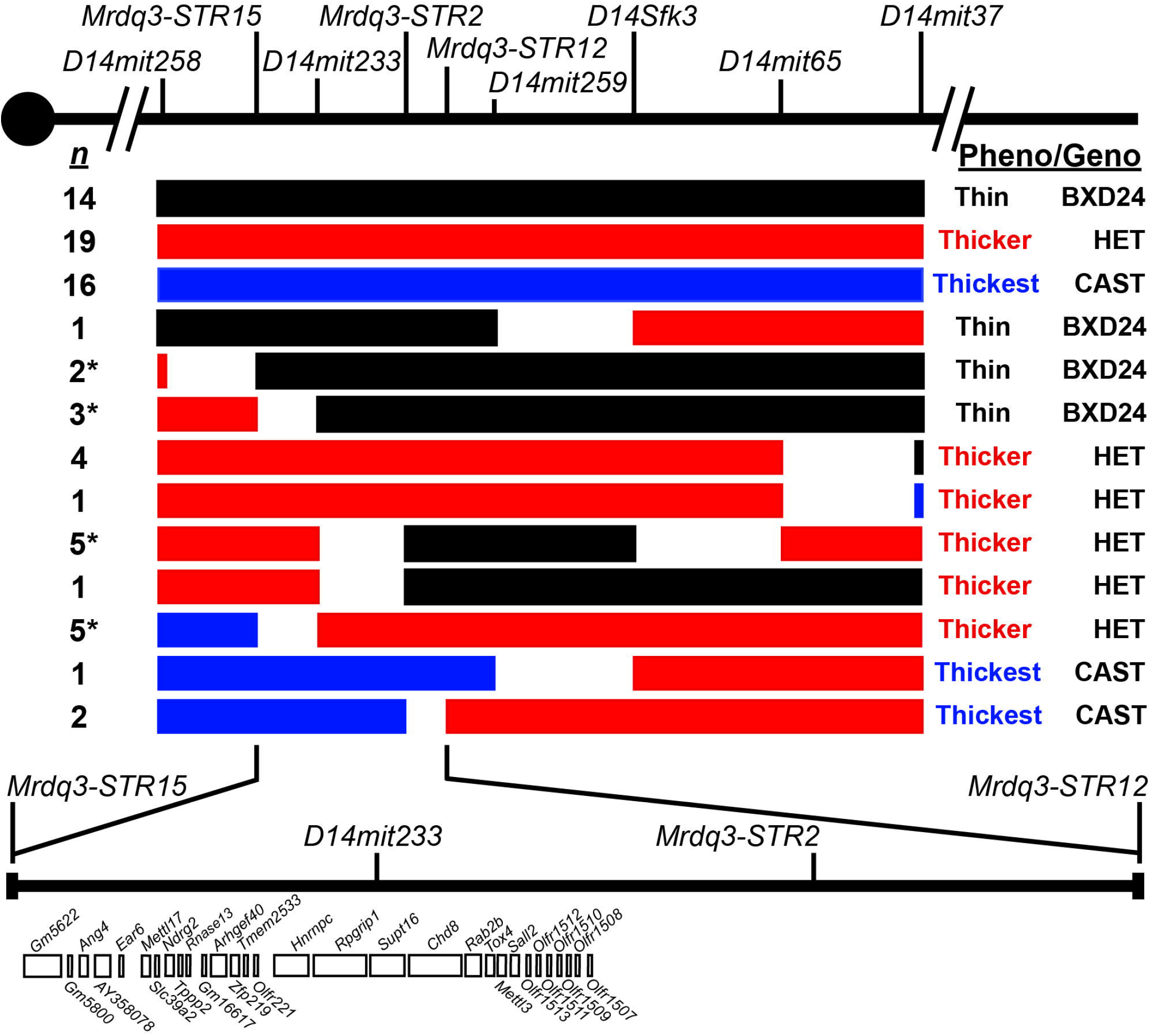
Genetic mapping of *Mrdq3* on mouse chromosome 14 using intercrosses of BXD24-*rd16*.CAST-*Mrdq3* congenic mice. *Black boxes* represent the BXD24 genotype associated with a thin retina, *red boxes* represent the heterozygous (HET) genotype associated with a thicker retina, and *blue boxes* represent the CAST genotype associated with the thickest retina. The number of mice (*n*) with each haplotype is listed to the left of each row, with progeny-tested mice denoted with an asterisk (*). The 6-week-old retina phenotype *(pheno;* measured by optical coherence tomography) and the deduced genotype (*geno*) for each haplotype is listed to the right of each row as “Thin BXD24”, “Thicker HET”, or “Thickest CAST”. The *vertical lines* across the chromosome represent markers that are polymorphic between BXD24 and CAST mice.

### Haplotype analysis to narrow the *Mrdq3* interval

Comparison of genome sequence across inbred mouse strains within the *Mrdq3* interval identified two major haplotypes: 1) those that match the C57BL/6J reference haplotype such as BXD24/TyJ, and 2) those that carry the more common alternate haplotype such as DBA/2J, 129-derived strains, BALB/cJ, and other commonly utilized laboratory strains. The CAST/EiJ strain carries a large number of private genetic variants on a mixed haplotype. To determine if *Mrdq3* was caused by a private or ancestral allele, DBA/2J was backcrossed to BXD24-*rd16* to generate 6-week-old N2 *rd16*-mutant mice for OCT analysis. Mice were genotyped for SNP marker *rs3681109,* which is located near the peak of *Mrdq3* within *Rpgrip1,* and retinal thickness plotted by genotype shows that mice with a D2 allele have a significantly increased retinal thickness compared to mice homozygous for BXD24 alleles (104.0 ± 2.8 μm vs. 97.2 ± 3.2, respectively; *p* = 8.9 x 10^-5^, Student’s two-tailed *t*-test; Figure 5a) indicating that *Mrdq3* is caused by an ancestral allele. Two additional inbred mouse strains, BXD63 and BTBR, have a break in ancestral haplotype within the *Mrdq3* interval and were additionally backcrossed to BXD24-*rd16* to generate 6-week-old N2 *rd16*-mutant mice for OCT analysis. Resulting mice were also genotyped for *rs3681109.* Mice with a BXD63 allele have the same retinal thickness as mice homozygous for BXD24 alleles (*p* = 0.758; Figure 5b) and mice with a BTBR allele have a significantly increased retinal thickness compared to mice homozygous for BXD24 alleles (112.9 ± 7.9 μm vs. 96.1 ± 6.3 μm; p = 5.4 x 10^-5^, Student’s two-tailed *t*-test; Figure 5c). Intersection of haploytype information and OCT results from each cross further narrowed *Mrdq3* to a 59.2 kb region containing *Hnrnpc* and the first seven exons of *Rpgrip1* (Figure 5d). Publicly available genome sequence (Mouse Genomes Project website) of the *Mrdq3* critical interval for the B6, D2, BTBR, and CAST mouse strains was filtered to identify genetic variants that don’t match the B6 reference sequence and where the D2, BTBR, and CAST strains all harbor the same allele. This resulted in a list of 144 candidate alleles for *Mrdq3* with only six alleles localized to coding sequence, all of these within *Rpgrip1* (Figure 5e; Supplementary Table 2). Three of the alleles *(rs3681636, rs3681648,* and *rs47971646)* are synonymous and three are missense changes *(rs3681109, rs3681193,* and *rs3681773).* The missense changes all occur near or within a low complexity protein region encoded by exon 4 of *Rpgrip1* with *rs3681109* resulting in Pro96Leu, *rs3681193* resulting in Pro110Ser, and *rs3681773* resulting in Ala137Val.

**Figure 5.**
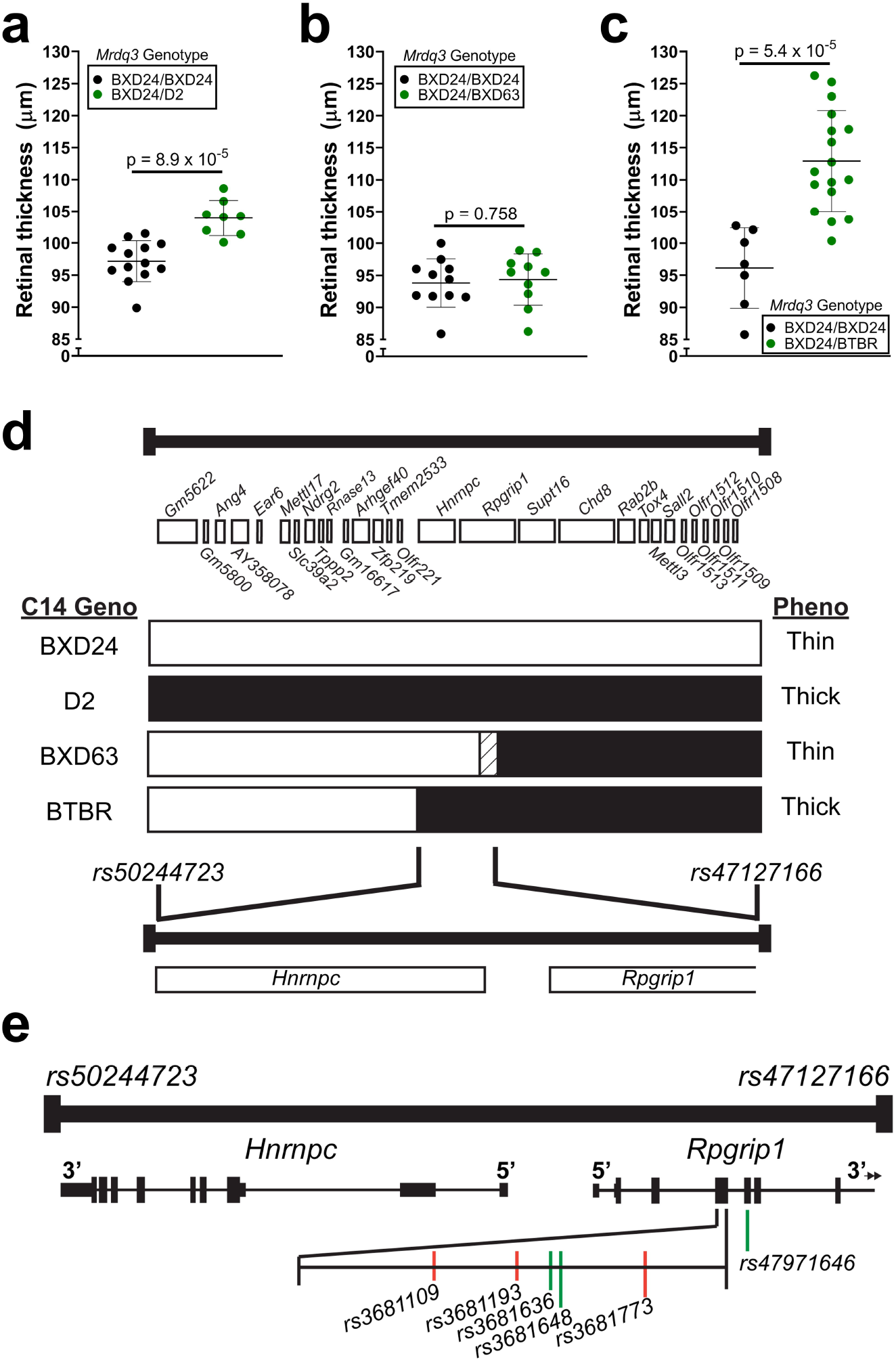
Haplotype analysis using strain-specific genome sequence to narrow the mouse chromosome 14 *Mrdq3* interval. Three inbred mouse strains with different *Mrdq3* ancestral haplotypes were backcrossed to BXD24-*rd16* mice and retinal thickness of 6-week-old N2 *rd16*-mutants was measured by optical coherence tomography for each cross. Results were plotted by genotype of *rs3681109* for **a** BXD24-*rd16*.D2 (*n* = 21 *rd16*-mutant mice), **b** BXD24-*rd16*.BXD63 (*n* = 21 *rd16*-mutant mice), and **c** BXD24-*rd16*.BTBR (*n* = 24 *rd16*-mutant mice). *Error bars* = mean ± SD. The *Mrdq3* haplotype map for each strain is depicted in panel **d** with the *white boxes* representing a haplotype matching the C57BL/6J reference, the *black boxes* representing the more common alternate haplotype, and the *white with diagonal-striped box* representing a region of unknown haplotype. The derivative mouse strain for each haplotype is listed to the left of each row. Note that the BXD63 and BTBR strains each have a recombination event within the *Mrdq3* interval. The retina thickness phenotype of the N2 *rd16*-mutant mice (shown in panels **a-c**) associated with an *Mrdq3* heterozygous genotype; i.e. BXD24/D2, BXD24/BXD63, or BXD24/BTBR; is listed to the right of each row as “thin” or “thick”. **e** The deduced critical region is flanked by markers *rs50244723* and *rs47127166* and contains the entirety of *Hnrnpc* as well as the 5’ portion of *Rpgrip1.* Publicly available genome sequence from the B6, D2, BTBR, and CAST strains of mice was used to filter for genetic variants in this region that differ from the B6 reference genome and where the D2, BTBR, and CAST strains all harbor the same allele. 144 genetic variants met these criteria and are listed in Supplementary Table 2. Six variants localized to coding sequence, all of them within *Rpgrip1;* three are synonymous *(green lines)* and three are missense *(orange lines).*

### *Mrdq1* exhibits grandparental-origin-dependent phenotypic effects

Our crosses to scan for modifiers were done in such a way that we could also determine the importance of directionality in our backcross. In the initial cross, a male CAST was mated to a female BXD24-*rd16*, while in the reciprocal cross, a female CAST was mated to a male BXD24-*rd16*. A female F1 from each direction of the cross (which are genetically identical, differing only in the sex of the parent transmitting each allele) was backcrossed to a male BXD24-*rd16* to produce the N2 generation. Separate one-dimensional scans of each backcross direction in the N2 dataset uncovered a surprising effect; *Mrdq1,* which was not identified in the analysis of the full N2 dataset (LOD 0.44; Figure 2c), was also not identified in the separate analysis of the initial backcross (LOD 0.04) but was significant (LOD 3.39) in the reciprocal cross (Figure 6a-b). Therefore, *Mrdq1* exhibits a grandparent-of-origin influence whereby the sex of the grandparent from whom the CAST allele is inherited may determine whether the allele confers a phenotypic influence.

**Figure 6.**
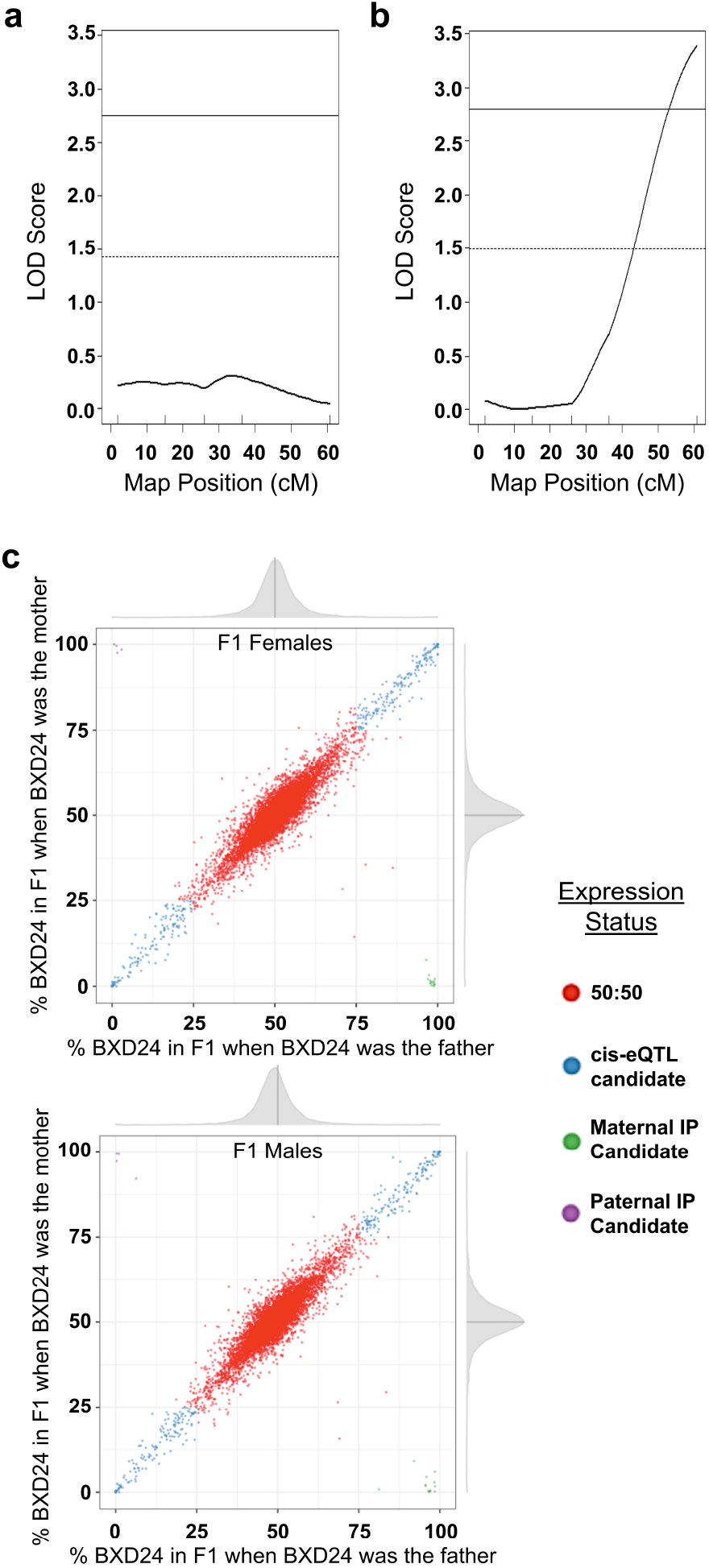
Grandparental-origin-dependent effect of *Mrdq1* on retinal thickness in *rd16*-mutant mice. To determine the importance of directionality in the quantitative trait locus (QTL) analysis, the backcross was set up in reciprocal between the inbred BXD24-*rd16* and CAST mouse strains. In the initial cross, a male CAST was mated to a female BXD24-*rd16*, while in the reciprocal cross, a female CAST was mated to a male BXD24-*rd16*. A female F1 mouse from each direction of the cross (genetically identical regardless of directionality) was backcrossed to a male BXD24-*rd16* mouse to produce the N2 generation. Separate QTL analyses of the **a** initial backcross and **b** the reciprocal backcross indicate the presence of the significant quantitative trait locus on Chr 12, *Mrdq1,* in the reciprocal backcross (LOD 3.39, *n* = 67 *rd16*-mutant mice with CAST grandmother) but not the initial backcross (LOD 0.04, *n* = 67 *rd16*-mutant mice with BXD24-*rd16* grandmother). Significance thresholds determined by permutation testing are indicated by the solid (*p* < 0.05; genome-wide significant) and dashed (*p* < 0.63; genome-wide suggestive) lines. Tick marks along the bottom of the plots indicate the location of genetic markers. LOD = logarithm of odds; cM = centimorgan) Quantification of differential allelic expression using RNA-Seq data from reciprocal (BXD24-*rd16* x CAST) F1 mice at p25 was used to identify imprinted genes in the retina. **c** Plots of joint distribution for F1 mice from the initial (BXD24-*rd16* mother) and reciprocal (BXD24-*rd16* father) crosses shows gene expression ratios: 1) genes with a 50:50 biallelic expression ratio, 2) cis-eQTL (expression QTL) candidate genes deviating from a 50:50 expression ratio independent of parental inheritance, 3) paternally imprinted candidate genes deviating from a 50:50 expression ratio with a bias for expression of maternally inherited alleles, and 4) maternally imprinted candidate genes deviating from a 50:50 expression ratio with a bias for expression of paternally inherited alleles. IP = imprinted.

### Identification of imprinted genes expressed in the mouse retina

The hereditary pattern exhibited by *Mrdq1,* along with the presence of a cluster of imprinted genes underlying the *Mrdq1* peak, led to the hypothesis that the causative gene is imprinted. To identify potential candidates for *Mrdq1,* RNA-Seq was used to establish a list of imprinted genes expressed in the retina of P25 mice. Reciprocal F1 hybrids of CAST and BXD24-*rd16* were generated in order to identify transcripts in the retina that exhibit differential allelic expression that is dependent on the parent-of-origin. The parent-of-origin for allelic expression can be determined for all expressed transcripts that include a SNP that is polymorphic between the parental mouse strains^40,41^, so the high degree of genetic polymorphism between CAST and BXD24-*rd16* make these strains well-suited for this experiment. Our analysis identified 40 genes that are both expressed and imprinted in the retinas of mice (Figure 6c; Table 3). Four of these genes are located within the *Mdrq1* interval: *Meg3, Rtl1, Rian,* and *Mirg,* and only *Meg3* expression differed significantly between the parent strains, with a 2.06-fold increased expression level (*p* = 1.99E_-3_; FDR = 9.78E^-3^) in BXD24 compared to CAST (Table 2). *Meg3* is a long non-coding RNA and genomic sequence data from the mouse Genomes Project shows 114 exonic genetic variants between CAST and D2 (the derivative strain for the *Mrdq1* region in BXD24 mice).

**Table 3.**
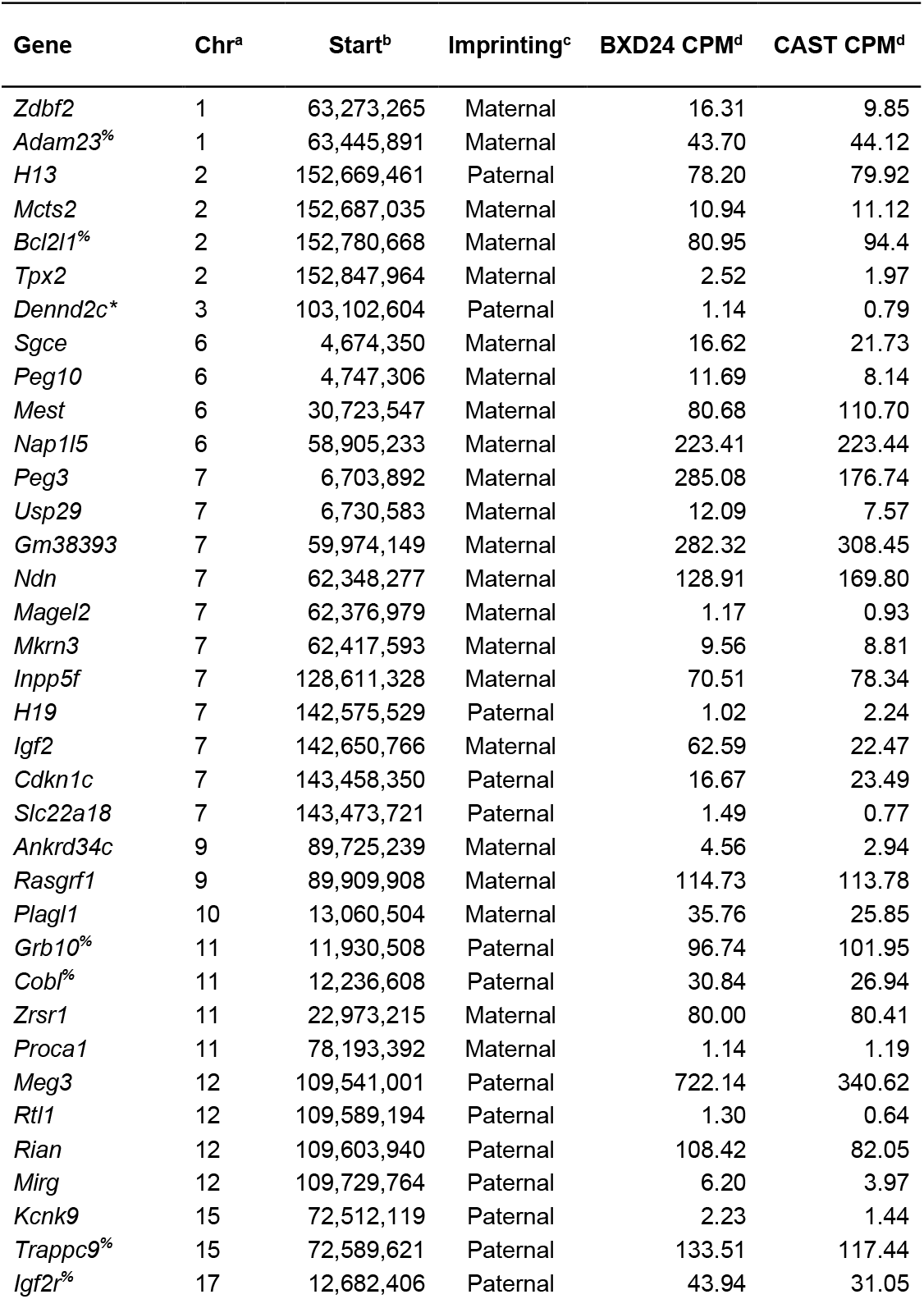
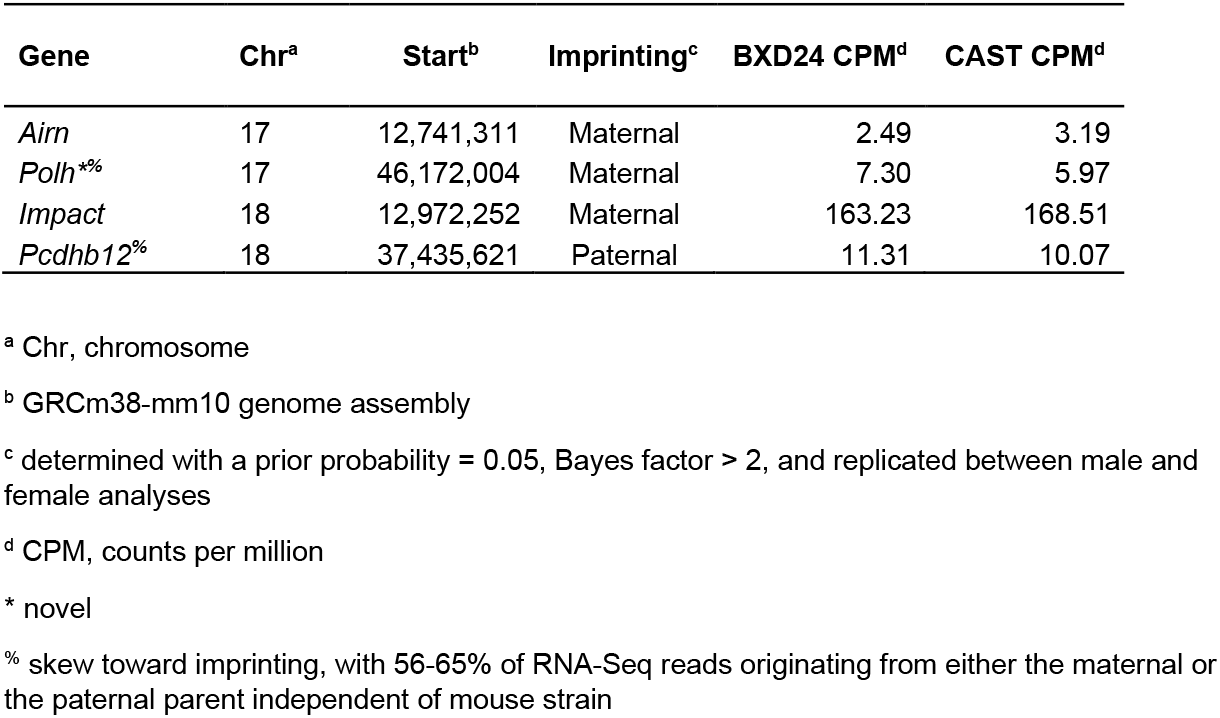
Genes that are expressed and imprinted in the P25 mouse retina.

### Identification of microRNA genes expressed in the mouse retina

Also underlying *Mrdq1* is one of the largest clusters of microRNA (miR) genes in the mouse genome with evidence that many, if not all, are imprinted (PMID: 12796779, 15310658). Therefore, miR-Seq was used to identify miRs expressed in the P25 mouse retina with differential expression between the BXD24 and CAST parental strains of mice. 593 small RNAs were expressed in the P25 retina with ≥ 5 reads per million in at least one of the parental strains; 87 of these were within the *Mrdq1* interval. The expression of 15 small RNAs within the interval was significantly different between CAST and BXD24 (padj < 0.05) and 8 have a log2 fold change > 0.5 or < −0.05 (padj < 0.02; Table 4). Of these, miR-341-3p stands out because it has the largest fold change (73.4-fold lower in CAST compared to BXD24); is highly expressed in BXD24 retina (584.58 reads per million); is the most significant change (padj = 1.63E-119); the sequence has a relatively high GC content, which some studies suggest promotes stability42; and its seed sequence is unique among miRs. Genomic sequence data from the Mouse Genomes Project shows that CAST has a 4-bp deletion in *Mir341* that is predicted to localize to the terminal loop of the pri-miRNA, a key site for miRNA processing, explaining the drastic reduction of mature miR-341-3p in the CAST retina.

**Table 4.**
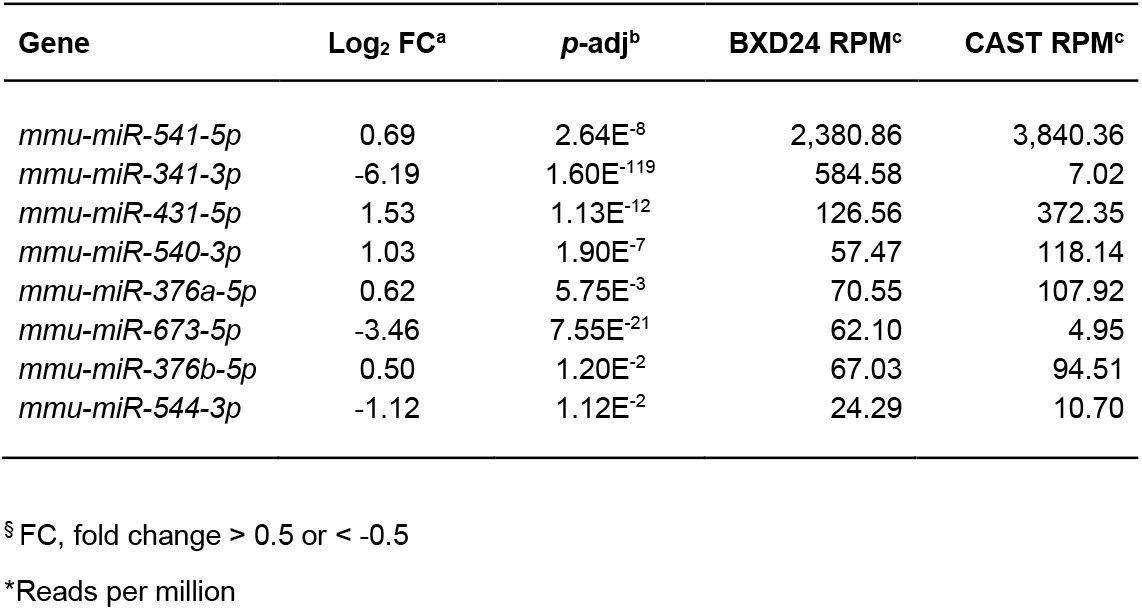
*Mrdq1* positional candidate microRNA genes with significant expression differences in P25 mouse retina between CAST and BXD24.

### Evaluation of *Mir341* as a candidate for *Mrdq1*

In situ hybridization with a probe for miR-341-3p in P25 BXD24 retina shows miR-341-3p expression in the ganglion cell layer, inner nuclear layer, and the inner segments of the photoreceptors (Figure 7a-b). To test the influence of *Mir341* on retinal thickness in *rd16*-mutant mice, the University of Iowa Genome Editing Core used CRISPR-Cas9 to generate a strain with a *Mir341* deletion by concurrently injecting two guide sequences, one targeting a region upstream of *Mir341* and the other targeting a region downstream. *Mir341* mutations were created on a pure D2 background, the parental strain-of-origin for the *Mrdq1* locus in BXD24 mice, because inbred BXD24 mice are relatively poor breeders and not available from The Jackson Laboratory in sufficiently large numbers. As expected, *Mir341* deletion in D2 mice did not change retinal thickness (Supplementary Data 1). The *D2-Mir341^Δ^* strain was then used in genetic crosses with BXD24-*rd16* to generate multiple cohorts of *rd16*-mutant mice. Cohorts were additionally either homozygous for the wild-type *Mir341* allele, homozygous for the *Mir341^Δ^* mutation, heterozygous for the *Mir341^Δ^* mutation due to paternal inheritance, or heterozygous for the *Mir341^Δ^* mutation due to maternal inheritance. Retinal thickness of mice within these cohorts was measured by OCT at 6 weeks of age and shows that deletion of *Mir341* does not modify retinal thickness in *rd16*-mutant mice in any of the contexts tested (Figure 7c-d).

**Figure 7.**
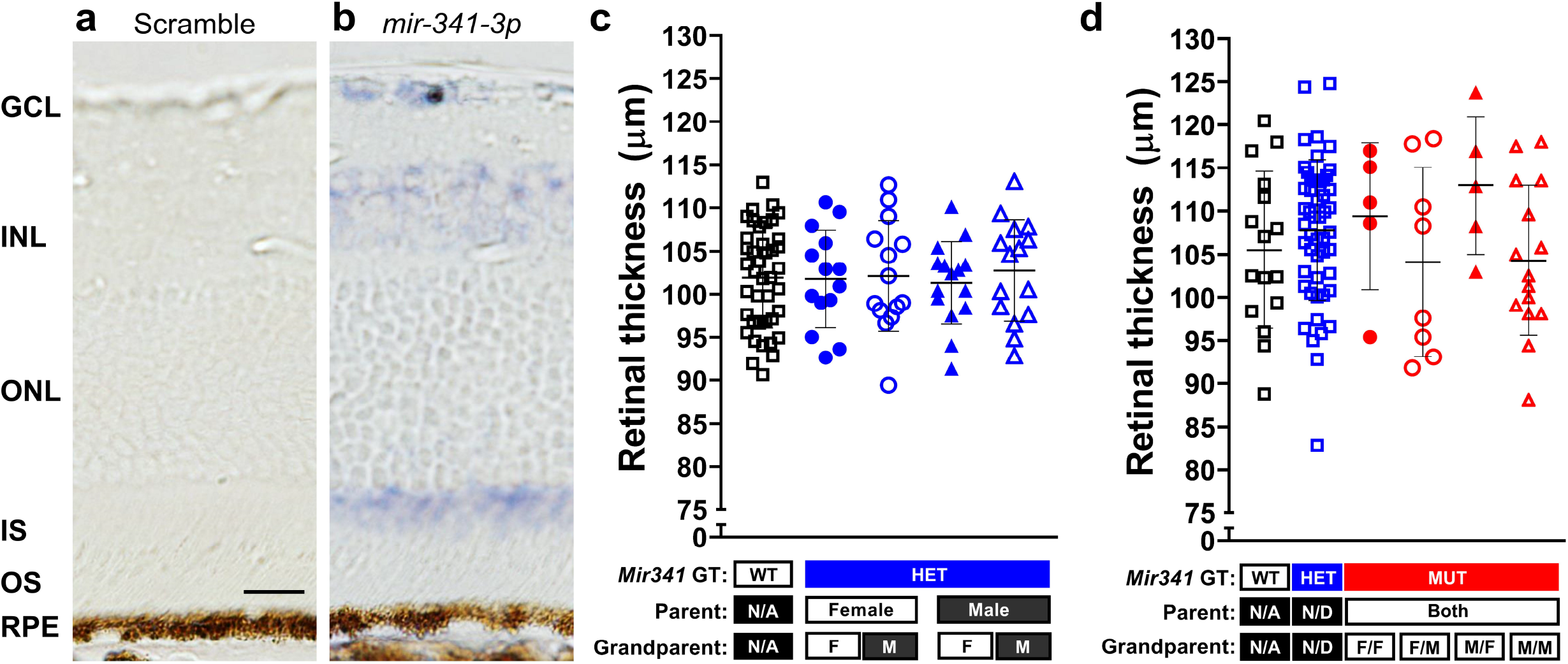
Retinal expression of *Mir341* and testing its influence on retinal thickness in *rd16*-mutant mice. Retinal sections from p25 BXD24 mice analyzed by in situ hybridization for **a** the Scramble-miR control and **b** miR-341-3p with phase contrast to identify the retinal layers. GCL, ganglion cell layer; INL, inner nuclear layer; ONL, outer nuclear layer; IS, inner segments of the photoreceptors; OS, outer segments of the photoreceptors; RPE, retinal pigment epithelium. Scale bar = 20 μm. Backcrosses and intercrosses between *D2-Mir341^Δ^* and BXD24-*rd16* were set up in a manner to test the influence of *Mir341* on retinal thickness in rd16-mutant mice and assess the importance of cross directionality. Retinal thickness was measured by optical coherence tomography in 6-wk-old *rd16*-mutant mice resulting from the **c** backcross and **d** intercross. *Error bars* = mean ± SD. *Mir341* genotype, sex of the parent transmitting the mutant allele, and sex of the grandparent transmitting the mutant allele is indicated. Sex of the parent or grandparent is listed as not applicable (N/A) for mice wild-type for *Mir341,* and retinal thickness measurements from each direction are combined. Sex of the parent or grandparent is listed as not determined (N/D) for F2 heterozygous mice because the allele could derive from either parent or grandparent, and retinal thickness measurements from each direction are combined. Grandparent sex for F2 mutant mice indicates the sex of the maternal grandparent transmitting the mutant allele *(left)* and the sex of the paternal grandparent transmitting the mutant allele *(right).* GT, genotype; WT, wild type; HET, heterozygous; MUT, mutant; F, female; M, male.

## Discussion

The initial discovery of a role for CEP290 in the retina came from phenotype-driven mouse genetics^43^, followed by the identification of *CEP290* mutations as the most common cause of LCA in humans^18^. CEP290 is a centrosomal protein and, in the retina, localizes to the photoreceptor connecting cilium between the inner and outer segments where it’s involved in the regulation of protein trafficking^5,44–47^. The BXD24-*rd16* strain of mice contains a spontaneous in-frame deletion of *Cep290* exons 35-39 and homozygosity for the *rd16* mutation leads to an early onset retinal degeneration characterized by progressive retinal thinning due to photoreceptor outer segment degeneration, analogous to human LCA patients. Among patients with CEP290-mediated retinal degeneration, the substantial variable expressivity in retinal phenotype and vision^23–26^ indicates that other factors, such as genetic modifiers^2,3,12^, are capable of influencing patient outcome.

To identify naturally occurring genetic modifiers of CEP290-mediated retinal degeneration in mice, we utilized a phenotype-driven genome-wide QTL mapping approach. BXD24-*rd16* mice were backcrossed and intercrossed with the genetically divergent wild-derived CAST strain, allowing maximal introduction of genetic variation into the background. Akin to the variable expressivity observed in human LCA patients, a segregating genetic background in the N2 and F2 populations of *rd16*-mutant mice resulted in a wide range of retinal thickness phenotypes, indicating the successful introduction of genetic modifiers from the CAST background. QTL analysis identified 11 chromosomal loci that significantly modify retinal thickness in *rd16*-mutant mice, named *Mrdq1-11,* together accounting for 71.7% of the observed phenotypic variation in retinal thickness.

Following the QTL analysis, we completed detailed studies of two of these loci, *Mrdq1* and *Mrdq3*. Alleles at each locus are capable of significantly suppressing *Cep290*-mediated retinal degeneration. Each locus also has other remarkable features that make them particularly tractable: *Mrdq3* accounts for 30.2% of the phenotypic variability in *rd16*-mutant mice (LOD 49.89) and *Mrdq1* shows evidence of imprinting. Through a combination of physical and molecular approaches we narrowed the critical region for each QTL and identified the probable causative genetic variations.

*Mrdq3* is caused by genetic variation in *Rpgrip1,* a gene in which loss-of-function mutations can cause LCA^48–49^. RPGRIP1 is a known binding partner of RPGR, the most common cause of X-linked retinitis pigmentosa, and localizes to the photoreceptor connecting cilium in mice and the outer segment in humans^50–54^. In the absence of RPGR, RPGRIP1 localization is unchanged^52^, remaining in the connecting cilia, which suggests that RPGRIP1 localizes to connecting cilia independently of RPGR. In contrast, RPGR is absent in the connecting cilia of photoreceptors lacking RPGRIP1^55^. Accordingly, mice lacking RPGRIP1 have a more severe retinal degeneration than mice lacking RPGR and the retinal degeneration phenotype of a double mutant is unchanged compared to a RPGRIP1 mutant^55^. CEP290 localizes to the transition zone of the photoreceptor connecting cilium and can directly bind cellular membranes and microtubules, potentially connecting the ciliary membrane to the microtubule-based axoneme^5^. The *rd16*-mutant CEP290 protein binds more RPGR than wild-type CEP290 and is also unable to bind microtubules, potentially causing abnormal morphology of the connecting cilia at P14 followed by rapid photoreceptor degeneration^5,46,56^. The finding that genetic variation in *Rpgrip1,* correlated with reduced expression, can suppress (rather than enhance, which is perhaps the simplest prediction based on previous studies) retinal degeneration in BXD24-*rd16* mice suggests a complex interplay between CEP290, RPGRIP1, and RPGR that can be tweaked to alter phenotypic outcome.

*Mrdq1* is caused by genetic variation in the *Meg3-Mirg* locus (a single polycistronic RNA containing long non-coding RNAs and miRNAs) previously unknown to influence LCA, which implicates genomic imprinting as a mechanism of phenotypic variation in *Cep290*-mediated retinal degeneration. Hereditary parent-of-origin effects as observed for *Mrdq1* are typically associated with imprinting^57,58^. The alleles of imprinted genes exhibit differential gene expression—and therefore differential phenotypic manifestation—dependent on whether the allele was inherited from the mother or from the father, and is an important source of phenotypic variable expressivity for complex traits^59–62^ Transgenerational imprinting is an unusual phenomenon, but has been firmly established to occur in a handful of cases^63–68^ and might be more common than currently appreciated. Among loci known to be sensitive to transgenerational imprinting, the callipyge locus of sheep merits particular attention^69–71^. Sheep with the callipyge mutation exhibit an unusual pattern of muscle hypertrophy that is inherited in a parent- and grandparent-of-origin manner^72^. The basis for this effect is thought to be an imprint that is typically, but not absolutely, reset after passage through the germline, such that an imprint can exhibit transgenerational influences. Strikingly, the region associated with callipyge is in a region of conserved synteny with, and/or overlaps, *Mrdq1.*

Future work will focus on testing the effect of *Rpgrip1* genetic variants on its ability to bind RPGR and CEP290 as well as continued elucidation of the role of genomic imprinting on CEP290-mediated retinal degeneration.

## METHODS

### Experimental animals

All animals were treated in accordance with the ARVO Statement for the Use of Animals in Ophthalmic and Vision Research. All experimental protocols were approved by the Institutional Animal Care and Use Committee at the University of Iowa. The inbred mouse strains BXD24/TyJ-*Cep290^rd16^*/J (Stock No: 000031; abbreviated as BXD24-*rd16* throughout), BXD24/TyJ (Stock No: 005243; abbreviated as BXD24 throughout), CAST/EiJ (Stock No: 000928; abbreviated as CAST throughout), DBA/2J (Stock No: 000671; abbreviated as D2 throughout), BXD63/RwwJ (Stock No: 007108; abbreviated as BXD63 throughout), and BTBR *T^+^ Itpr3^tf^*/J (Stock No: 002282; abbreviated as BTBR throughout) were purchased from The Jackson Laboratory and subsequently housed and bred at the University of Iowa. The *Cep290^rd16^* mutation arose spontaneously on the recombinant inbred BXD24/TyJ genetic background and results in early onset retinal degeneration. BXD24-*rd16*.CAST (F1) and CAST.BXD24-*rd16* (F1) mice heterozygous for the *Cep290^rd16^* mutation were generated. F1 mice were either intercrossed to generate BXD24-*rd16*.CAST (F2) and CAST.BXD24-*rd16* (F2) populations (abbreviated as F2 throughout), or female F1 mice were backcrossed with male BXD24-*rd16* mice to generate (BXD24-*rd16*.CAST).BXD24-*rd16* (N2) and (CAST. BXD24-*rd16*). BXD24-*rd16* (N2) populations (abbreviated as N2 throughout). F2 and N2 mice were PCR-genotyped for the *Cep290^rd16^* mutation using a three primer pool: a forward primer specific to the wild-type allele (5’-CCACCTGACCTCTAAACTCCCT-3’), a forward primer specific to the mutant allele (5’-ATAGTGGGACATTTTTATAAGGCACAGTGG-3’), and a reverse primer that recognizes both alleles (5’-GCAGCATGAGATGGAATCCTTCTTAGG-3’).

### Retinal thickness phenotyping

All retinal thickness measurements were recorded from mice at the age listed in the experiment ± 3 days. Mice were injected with a standard mixture of ketamine/xylazine (intraperitoneal injection of (100 mg ketamine + 10 mg xylazine)/kg body weight (Ketaset^®^, Fort Dodge Animal Health, Fort Dodge, IA; AnaSed^®^, Lloyd Laboratories, Shenandoah, IA). Mice were provided supplemental indirect warmth by a heating pad during anesthesia. Upon anesthesia, eyes were hydrated with balanced salt solution (BSS; Alcon Laboratories, Fort Worth, TX). To obtain retinal images, the tear film was wicked away and eyes were imaged with a Bioptigen spectral domain optical coherence tomographer (SD-OCT; Bioptigen, Inc., USA) using the mouse retinal bore. The volume intensity projection was centered on the optic nerve. Scan parameters were as follows: rectangular volume scans 1.4 mm in diameter, 1000 A-scans/B-scan, 100 B-scans/volume, 1 frame/B-scan, and 1 volume. Following imaging, eyes were hydrated with artificial tears and mice were provided supplemental indirect warmth for anesthesia recovery. Retinal thickness was measured from retinal images within the Bioptigen InVivoVueClinic Software three times per eye using vertical angle-locked B-scan calipers. In wild-type retinas, measurements were made from the internal limiting membrane to the contact cylinders of the photoreceptor cell layer because it was not always possible to resolve the band corresponding to the contact cylinders of the photoreceptors from the band corresponding to the retinal pigment epithelium (RPE). In *rd16* mice, measurements were made from the internal limiting membrane to the RPE. The six retinal thickness measurements obtained from two eyes of each animal were averaged together to produce a single retinal thickness measurement per animal. All retinal thickness measurement values are reported as an average ± standard deviation (SD).

### Mouse genotyping

Genome-wide genotyping was performed using Fluidigm technology as previously described73. In the subsequent QTL analyses of Chr 14, F2 and N2 *Cep290^rd16/rd16^* mice were genotyped with five simple sequence length polymorphism markers within the region of interest *(D14mit212, D14mit18, D14mit259, D14mit102,* and *D14mit224*). Select F2 and N2 *Cep290^rd16/rd16^* mice with chromosomal recombinations occurring within the region of interest on Chr14 were genotyped with an additional five markers *(D14mit258, D14mit62, D14mit233, D14Sfk3,* and *D14Sfk5).* All primer sequences are publicly available on the Mouse Genome Informatics website74.

### Quantitative trait locus (QTL) analysis

Using 90 polymorphic genetic markers evenly dispersed through the genome, QTL analysis of retinal thickness was performed with R/qtl using the N2 and F2 *Cep290^rd16/rd16^* datasets separately and combined. The genome-wide scan (scanone) was conducted using the logarithm of odds (LOD) score at 1.0-cM steps with standard interval mapping by the EM algorithm. To determine the significance thresholds for the scanone, permutation testing was performed with 1,000 permutations. Loci with LOD scores above the *p* = 0.05 threshold were considered significant QTL while loci with LOD scores above the *p* = 0.63 threshold were considered suggestive75. Gene-gene interactions were determined by performing a two-dimensional genome scan (scantwo) as previously described. To further examine evidence for individual QTL, a multiple regression analysis was performed. In this analysis, each QTL is dropped from the multiple-QTL model one at a time and the full model is compared to the model with the QTL omitted. If the full model is more probable, then the LOD score of the omitted QTL will surpass significance thresholds calculated by permutation testing in the scanone. Only the QTL that remain suggestive/significant after multiple regression analysis are reported. The refineqtl function was used to improve the QTL location estimates.

### Retinal Fundus Imaging

The pupils of mice were dilated using a combination of 2% cyclopentolate hydrochloride ophthalmic solution (Cyclogyl^®^, Alcon Laboratories, Fort Worth, TX) and 2.5% phenylephrine hydrochloride ophthalmic solution (Paragon BioTeck, Inc., Portland, OR). When pupils were fully dilated, mice were injected with a standard mixture of ketamine/xylazine (intraperitoneal injection of (100 mg ketamine + 10 mg xylazine)/kg body weight (Ketaset^®^, Fort Dodge Animal Health, Fort Dodge, IA; AnaSed^®^, Lloyd Laboratories, Shenandoah, IA). Mice were provided supplemental indirect warmth by a heating pad during anesthesia. Immediately upon anesthesia, hypromellose 2.5% ophthalmic demulcent solution (Goniovisc^®^, HUB Pharmaceuticals, LLC, Rancho Cucamnoga, CA) was applied to each eye. Eyes were imaged with a Micron III retinal imaging microscope (Phoenix Research Labs, Pleasanton, CA). Following imaging, eyes were hydrated with artificial tears and mice were provided supplemental indirect warmth for anesthesia recovery.

### Electroretinogram

Electroretinograms were performed on 7-week-old F2 *rd16*-mutant mice to compare mice with a relatively thin retina (< 116 μm) to mice with a relatively thick retina (> 150 μm). To minimize potential confounders, each pair of mice was precisely age-matched and analyzed by ERG in the same session. Mice were placed in a light tight, darkened cabinet overnight for dark adapting their vision. The following morning, the mice were anesthetized using a ketamine/xylazine anesthetic mix. Once anesthetized, mice were moved to the testing equipment and electrodes were placed on the eyes with a drop of hypromellose ointment to keep the eyes from drying out and ensuring good electrical conduction. A ground needle was also placed subcutaneously at the base of the tail. A program of gradually increasing light flashes was run while recording the electrical signal from the electrodes. After testing the mice were placed on a heating pad until recovered from anesthesia.

### RNA-Seq

All RNA-seq experiments used RNA extracted from pools of 4-6 circadian-matched retinas (collected between 4-5 hours from lights-on) from 25-day-old mice. Three biological replicates were collected for each of eight samples (male and female of each of four mouse strains: CAST/EiJ, BXD24/TyJ, BXD24-*rd16*.CAST (F1) and CAST.BXD24-*rd16* (F1)). BXD24/TyJ is the wild-type coisogenic control for BXD24-*rd16* and was used because an intact normal retina with all cell types present was necessary for this set of experiments. To elaborate, retinas from BXD24-*rd16* mice would show aberrant expression of all genes in the retina, especially those localized to the degenerated outer layers, which could mask and confound the expression changes pertinent to these experiments. Mice were euthanized by cervical dislocation and eyes were immediately harvested and placed in RNAlater^®^ Solution (Life Technologies, Carlsbad, CA). Retinas were dissected from eyes in RNAlater^®^ Solution, placed in Lysis/Binding Buffer from the mirVana™ miRNA Isolation Kit (Life Technologies, Carlsbad, CA), and homogenized using a rotor/stator homogenizer. RNA was extracted using the mirVana™ miRNA Isolation Kit.

The DNA*free* Kit (Life Technologies, Carlsbad, CA) was used to remove contaminating genomic DNA. The integrity of RNA was assessed using a 2100 Bioanalyzer (Agilent Technologies, Santa Clara, CA); samples with an RNA integrity number ? 8.0 were used for RNA-Seq. Retinal cDNA libraries were created by the Iowa Institute of Human Genetics Genomics Division using the TruSeq Stranded mRNA Library Prep Kit and the TruSeq Small RNA Library Prep Kit (Illumina, Inc., San Diego, CA). Indexed libraries prepared from mRNA were normalized and pooled into three groups (corresponding to three biological replicates) of eight samples. Each pool of eight samples was split between two lanes of a HiSeq 2500 (Illumina, Inc., San Diego, CA) for 100 x 100 paired end sequencing. Indexed libraries prepared from small RNA were normalized and all 24 libraries were pooled and split between two lanes of a MiSeq (Illumina, Inc., San Diego, CA) for 50 base-pair single read sequencing.

### RNA-Seq Bioinformatic Analyses

A hybrid BXD24/TyJ transcriptome was constructed by first mapping RNA-Seq reads from BXD24 parental animals to Ensembl transcript sequences specific to either C57BL/6J and DBA/2J (available at: https://www.sanger.ac.uk/data/mouse-genomes-project/^76^. The mmseq software (ver. 1.0.8)^77^ was used to quantify the number of reads mapping to strain-specific transcripts. Regions of consecutive strain-specific transcripts were identified. These regions were used to create a single BXD24 transcriptome file for use in downstream analyses.

RNA-Seq reads were mapped to the publicly available CAST transcriptome, the BXD24 transcriptome, or a hybrid CAST-BXD24 transcriptome using Bowtie (ver. 1.1.1)^78^, using the parameters suggested in the mmseq documentation (https://github.com/eturro/mmseq/blob/master/doc/countsDE.md). The number of reads mapping to each transcript was calculated using the bam2hits program bundled with mmseq, and the resulting data files were used as input for mmseq.

Differential expression analysis was performed using edgeR (ver. v.3.16.5)^79^ to compare BXD24 and CAST inbred strains. Prior to analysis, mmseq output was converted to read count equivalents using the script provided with the mmseq software (https://github.com/eturro/mmseq/tree/master/src/R). Biological sex was included as a covariate in the model.

RNA-Seq samples from F1 animals were analyzed for evidence of cis-eQTL regulation, maternal imprinting, and paternal imprinting using the mmdiff software (ver. 1.0.8)^80^. Briefly, a set of design matrices were constructed corresponding to different models of expression as described previously^81^. The expression pattern of each gene was fit to these models, then the model with the best fit the expression pattern was selected. For this analysis, male and female animals were analyzed separately, and only polymorphic, autosomal reads were included (i.e., X, Y, and MT genes were excluded)^81^.

### miRNA-Seq Bioinformatic Analysis

To detect differences in expression of small RNA populations, miRNA-Seq reads were analyzed using the OASIS software pipeline (ver. 2.0)^82^. One miRNA-Seq sample from a male BXD24 mouse was identified as an outlier based on its position in the PCA plot of all small RNA species and also had similar numbers of trimmed reads that were flagged by Oasis as too short; this sample was removed prior to statistical analysis. Differential expression of miRNAs between BXD24 and CAST samples was determined using the DESeq2 algorithm^83^ as implemented in the OASIS pipeline. Small RNA transcripts were retained for analysis if their mean expression was at least 5 reads.

### Generation and analysis of BXD24-*rd16*.Cg-CAST^*Mrdq3*^

*BXD24-rd16.Cg-CAST^Mrdq3^* mice were generated by reiterative backcrossing to BXD24-*rd16* mice for 10 generations, breeding mice carrying the CAST-derived Chr 14 segment corresponding to *Mrdq3,* selected at each generation by the genetic markers *D14mit258* and *D14mit102.* N10 mice heterozygous for *Mrdq3* were intercrossed to produce N10F2 progeny. N10F2 mice were genotyped for *Mrdq3* and their retinal thickness was measured by OCT at 6 weeks ± 3 days of age. Only the mice with a recombination in the *Mrdq3* interval and their nonrecombinant littermates were phenotyped. Chromosomal breakpoints were further validated through testcrossing in which mice with a critical recombination were backcrossed to BXD24-*rd16* mice and segregation analysis was performed on the resulting progeny. Three primer sets capturing short tandem repeat polymorphisms between BXD24 and CAST were designed inhouse and to provide additional positional genotype information for recombination mapping. Primer sequences are as follows: 1) *Mrdq3-STR15* F: TGGGGTTTAATTGTTTTAGG and R: AAAACCCTCCTCTAAAAACC, 2) *Mrdq3-STR2* F: TCCTTGGTTCTTGTGTTCC and R: AAAGAATAGATAATGAGAACTTAGTGG, and 3) *Mrdq3-STR12* F: AGACTCTTTCCCACCTTCTC and R: TGGTGAAACTAACCAAACAAG

### Construction and analysis of *D2-Mir341^Δ^* mice

Mice with a deletion of *Mir341* on a pure DBA/2J genetic background were generated by the Genome Editing Facility at the University of Iowa using two CRISPR-Cas9 guide sequences simultaneously: 1) TCTTACCTCCTAGATCACTA and 2) ATCACCCACCCTGATTGCAT. D2 was used as the background strain because the *Mrdq1* region in BXD24-*rd16* mice is derived from D2 and this ameliorates the introduction of confounding alleles in genetics crosses. Founders were crossed with DBA/2J mice and offspring analyzed by Sanger sequencing for germline transmission of *Mir341* deletion.

### miRNA in situ hybridization

Anesthetized 25-day-old mice were perfused with ice cold 4% paraformaldehyde. Enucleated eyes were dissected to remove the lens and fixed in 10% neutral buffered formalin at room temperature overnight. Fixed eyes were embedded in paraffin and 6 μm sections were collected. In situ hybridization was performed using the following Double-DIG-labeled miRCURY LNA™ Detection Probes (Exiqon, Woburn, MA): mmu-miR-341-3p (Product no. 615525-360), Scramble-miR (negative control; Product no. 699004-360) and U6, hsa/mmu/rno (positive control; Product no. 699002-360), and following the manufacturer’s protocol for the microRNA ISH Buffer Set – FFPE (Product no. 90000). After deparaffinization, sections were incubated with 10 μg/mL Proteinase-K for 25 minutes at 37°C. Probe hybridization was performed at 52°C for one hour.

## Supporting information

Supplementary Figure 1: QTL Effect Plots

Supplementary Table 2: Mrdq3 Haplotype SNPs

Supplementary Table 1: Differentially expressed Mrdq positional candidate genes

## Notes

### Competing Interest Statement

The authors have declared no competing interest.

