## Supplementary Figure 1: QTL Effect Plots for "Genetic modifiers of *Cep290*-mediated retinal degeneration"

**Figure. Effect plots for *Mrdq1-11***. The retinal thickness phenotype (mean ± SE) as a function of genotype is plotted for each quantitative trait locus (QTL) peak in the **a-j** combined dataset and **k** F2 dataset. **l,m** The retinal thickness phenotype (mean ± SE) as a function of genotype for two interacting QTL is plotted for the combined dataset.

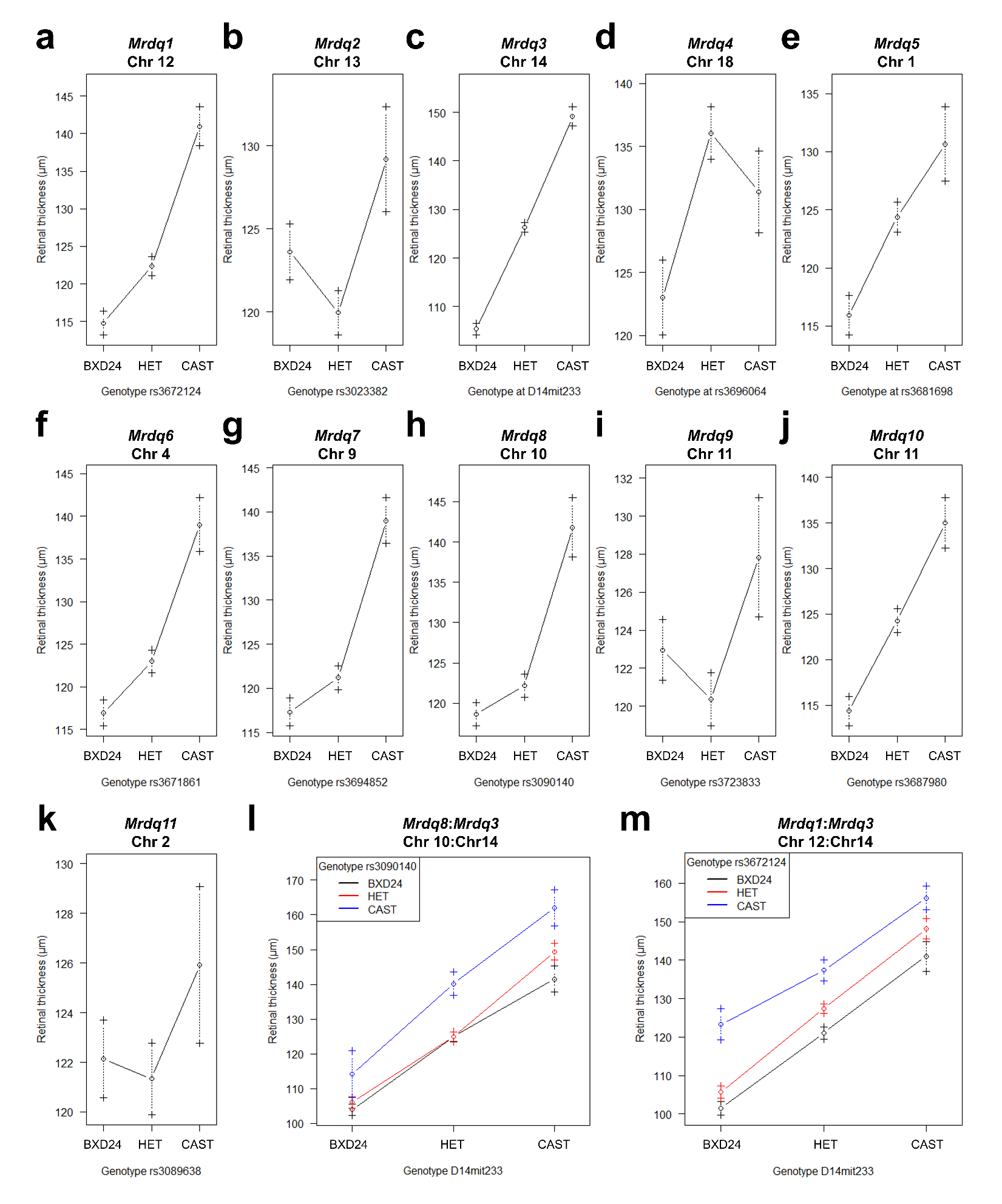
