## Supplementary Data 1 for "Genetic modifiers of *Cep290*-mediated retinal degeneration"

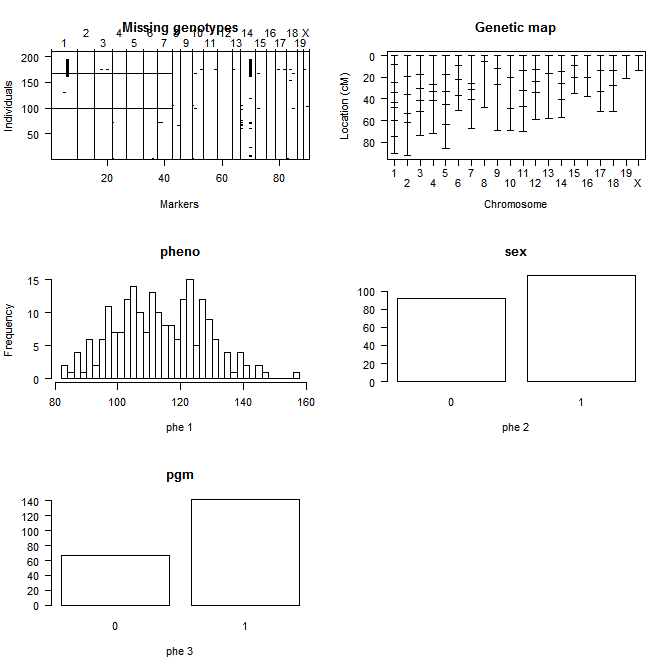

N2 RT histogram

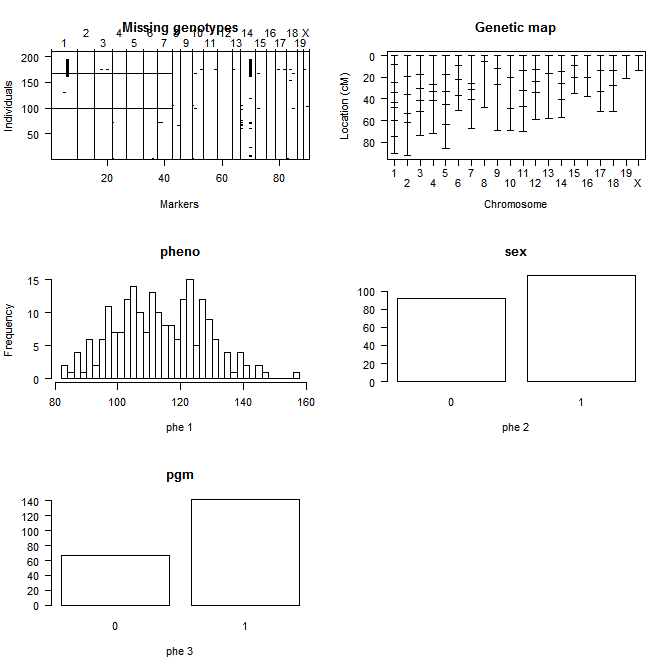

N2 RT looking at cross directionality

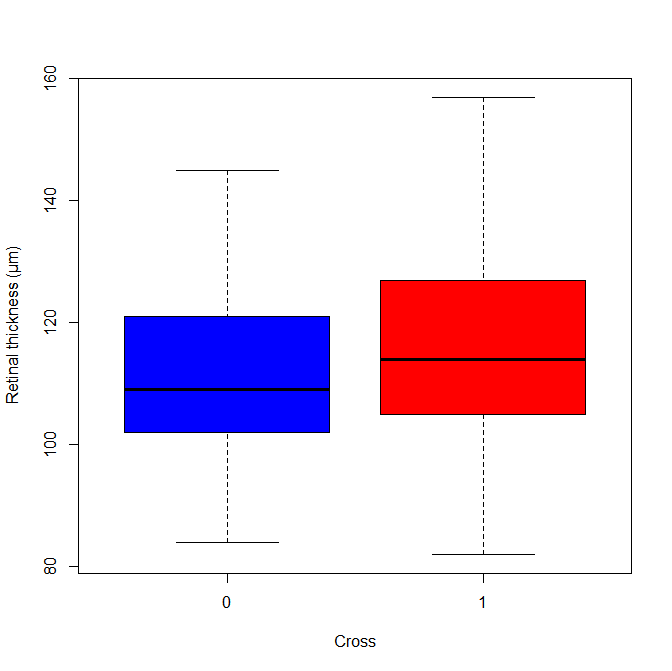

F2 RT histogram

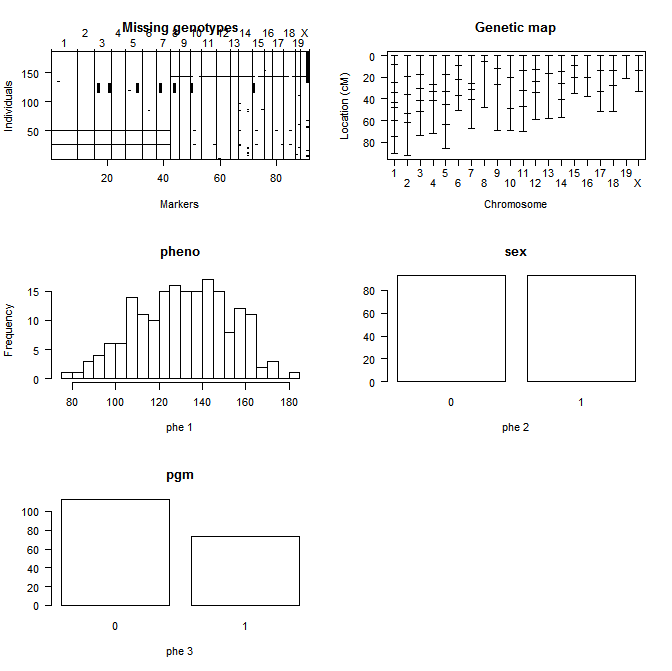

F2 RT looking at cross directionality

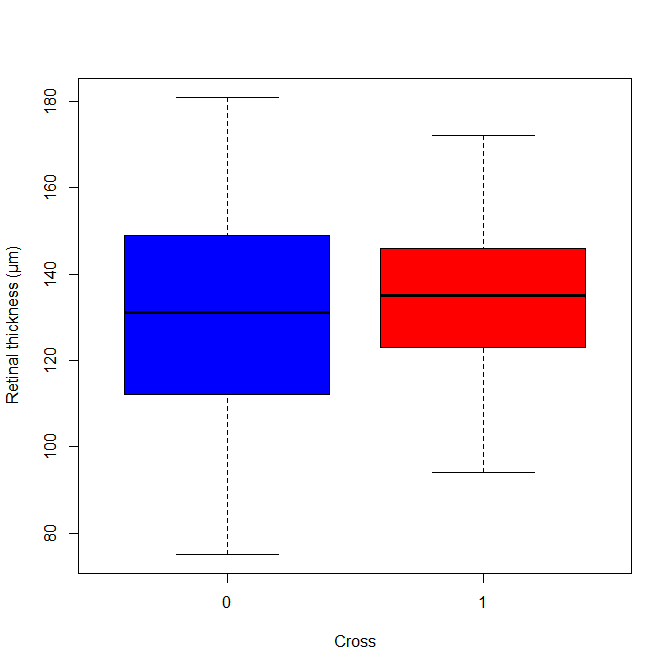

Combined – initial scanone without chr14 mit markers

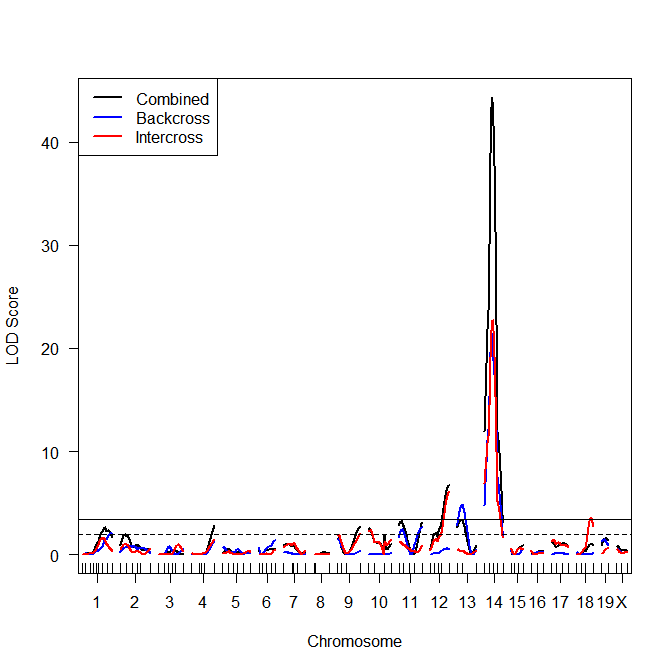

Combined – whole genome scantwo lodfv1 left lodav1 right

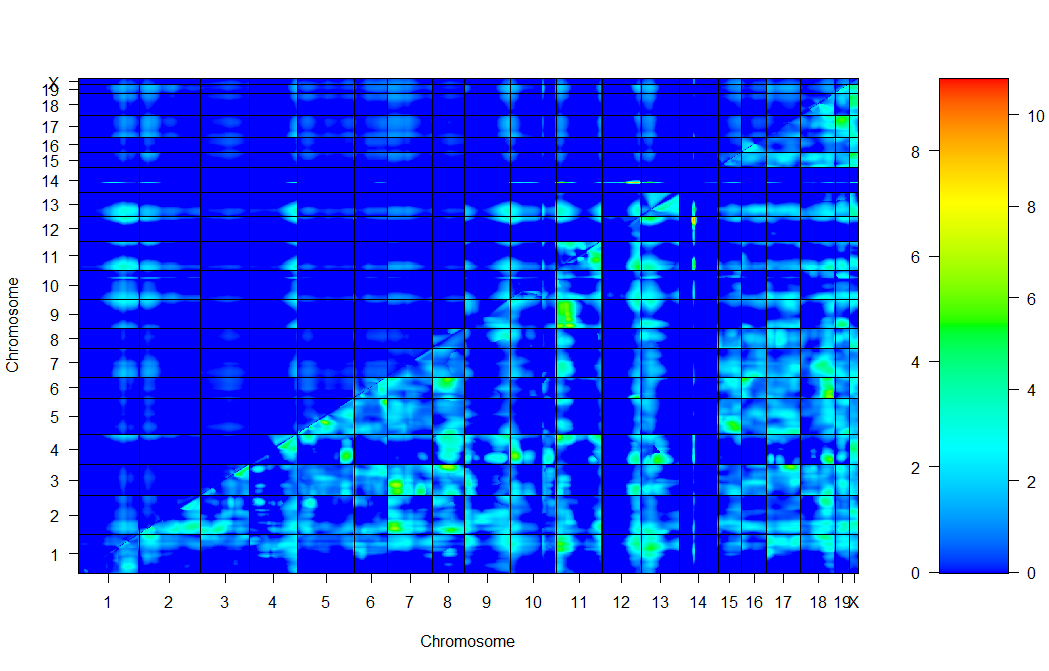

Combined scantwo chr1,12 lodfv1 left lodav1 right

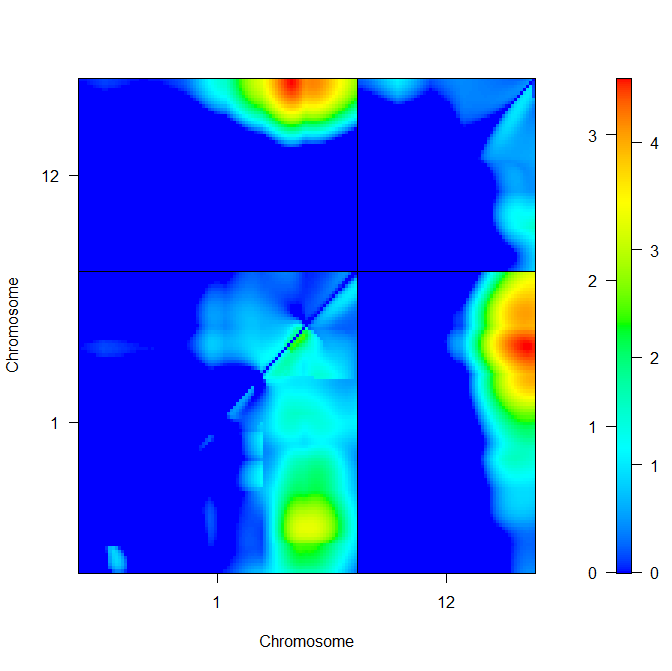

Combined scantwo chr1,14 lodfv1 left lodav1 right

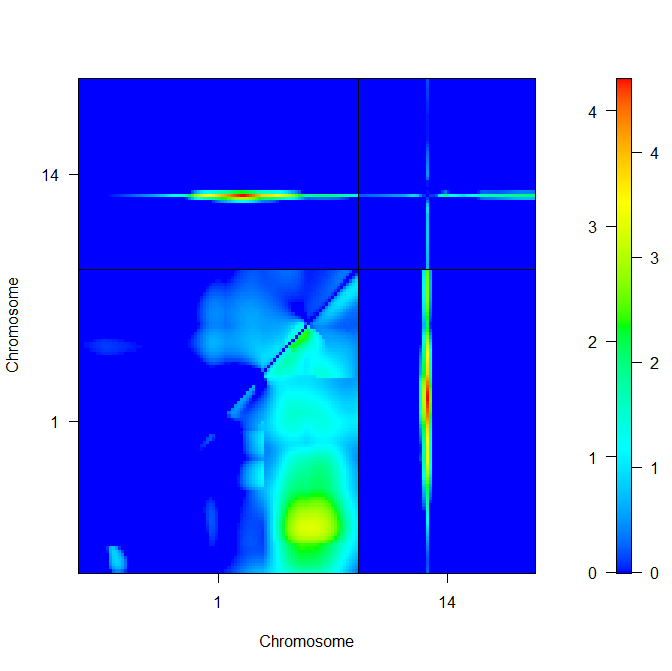

Combined scantwo chr2,14 lodfv1 left lodav1 right

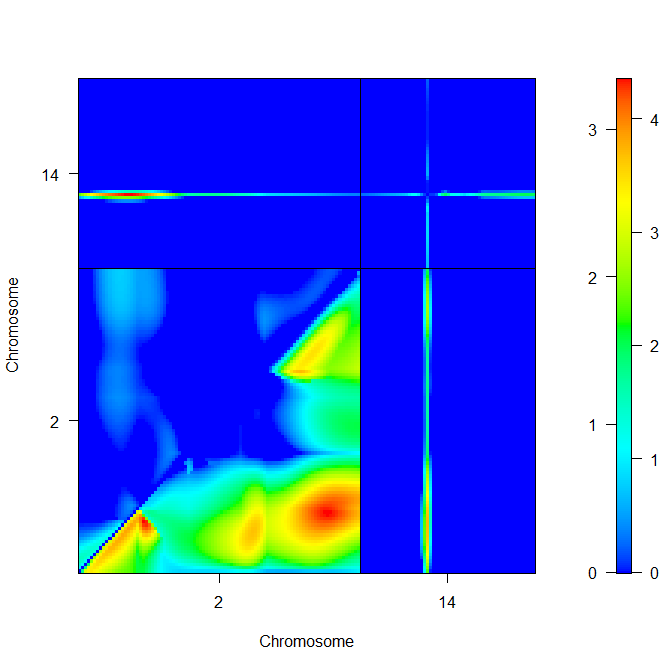

Combined scantwo chr4,14 lodfv1 left lodav1 right

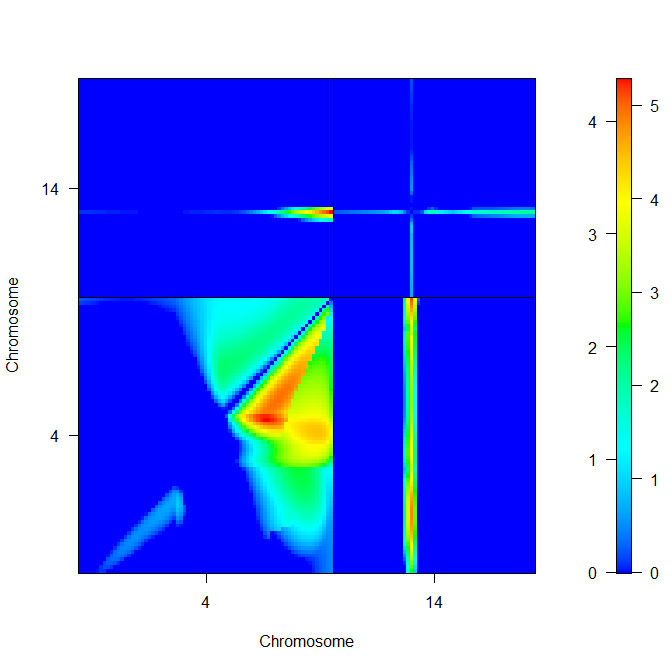

Combined scantwo chr9,14 lodfv1 left lodav1 right

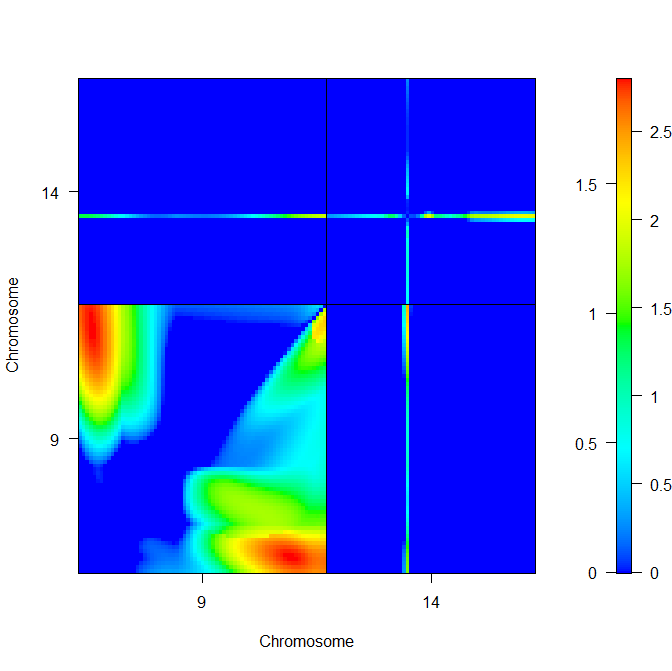

Combined scantwo chr10,14 lodfv1 left lodav1 right

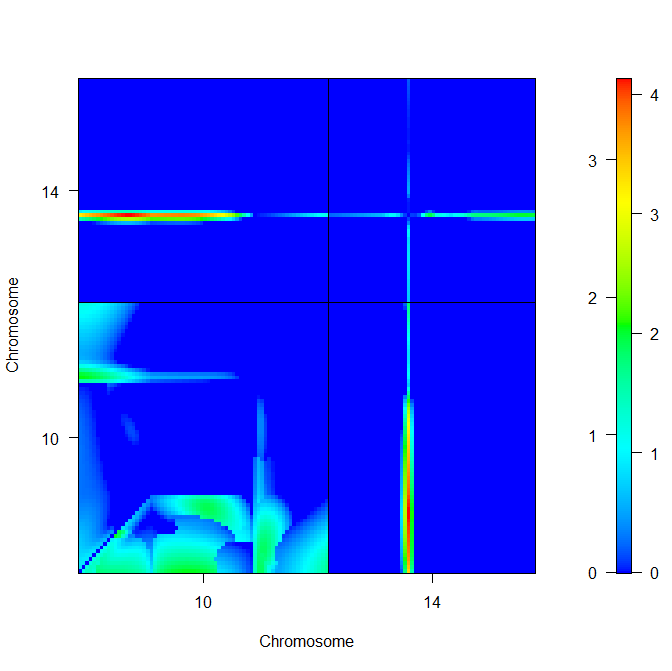

Combined scantwo chr11,14 lodfv1 left lodav1 right

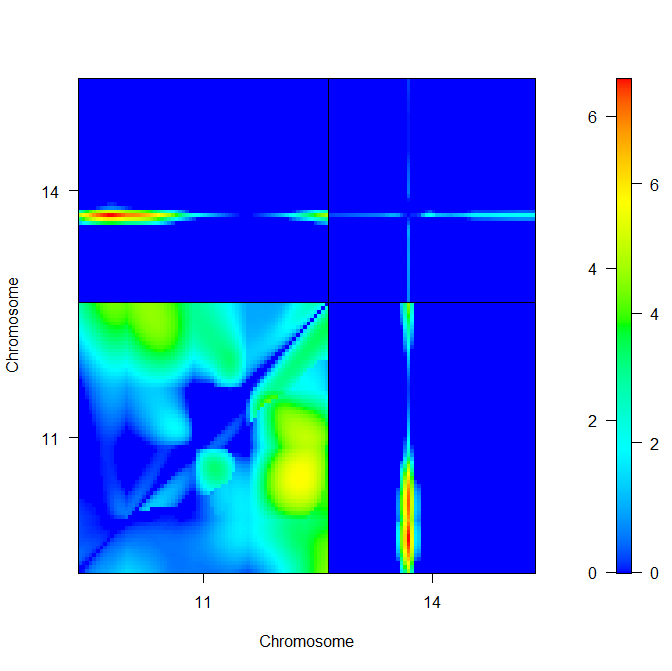

Combined scantwo chr11 lodfv1 left lodav1 right

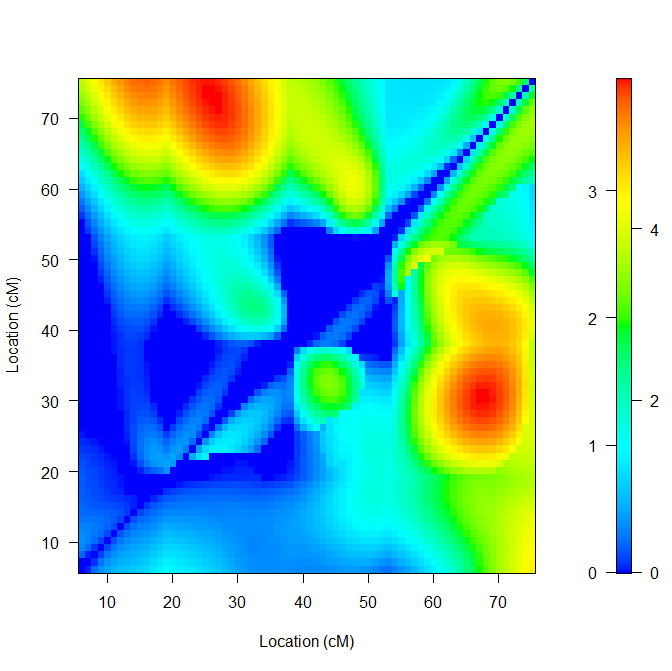

Combined scantwo chr12,14 lodfv1 left lodav1 right

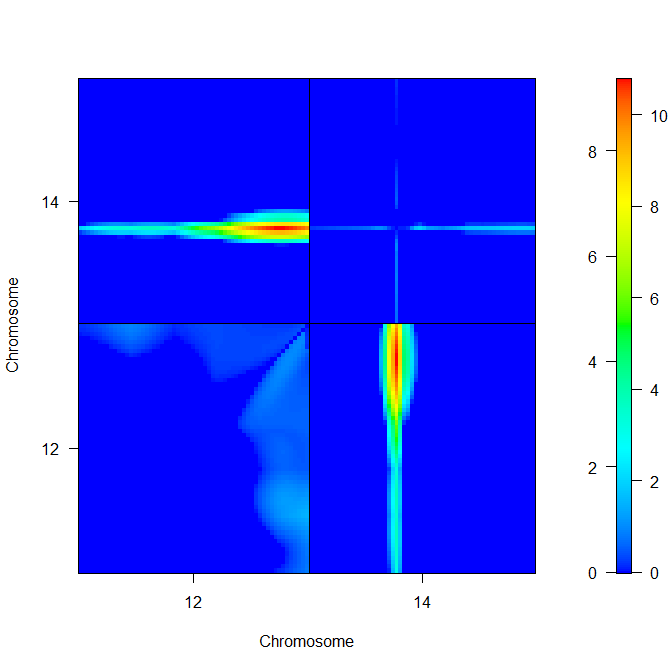

Combined scantwo chr13,14 lodfv1 left lodav1 right

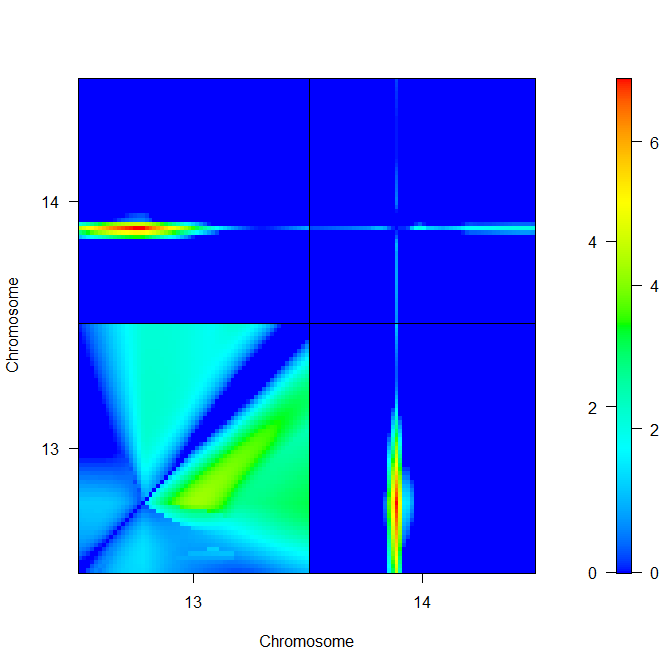

Backcross scantwo genome upper lodfv1 lower lodav1

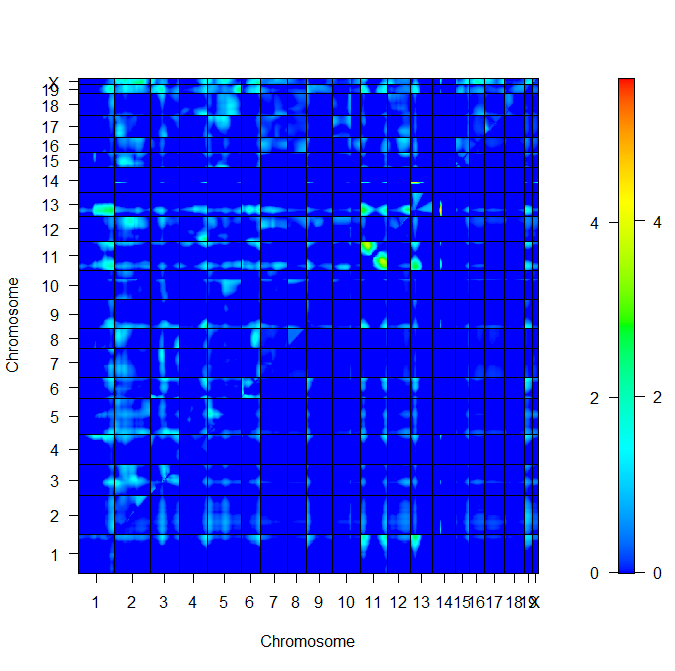

Intercross scantwo genome upper lodfv1 lower lodav1

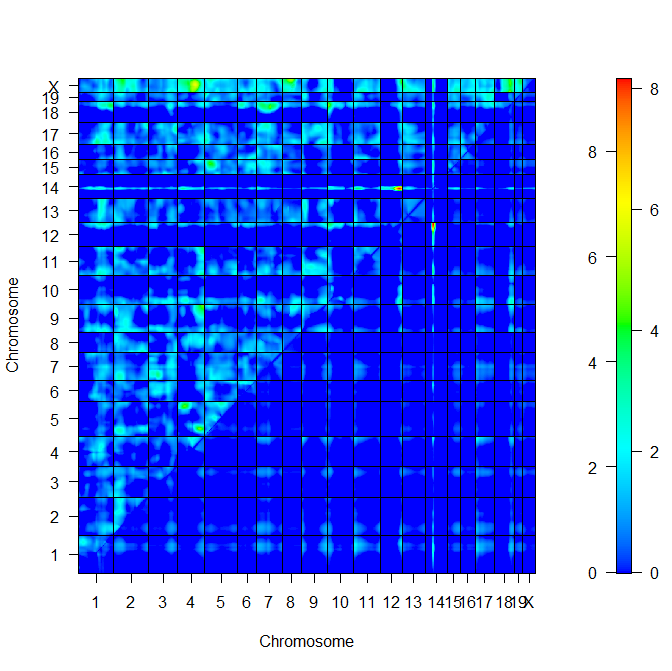

Intercross scantwo chr12,14 lodfv1 left lodav1 right

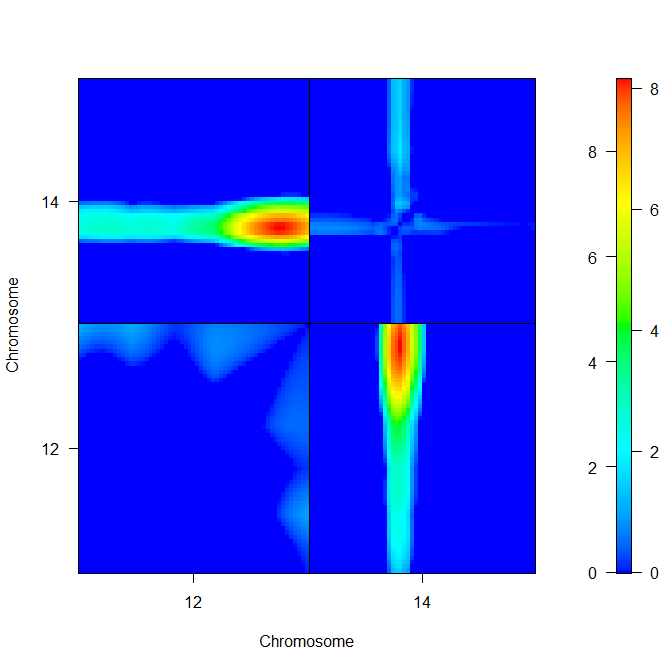

CRISPR-Cas9 targeting strategy of deletion of *Mir341*

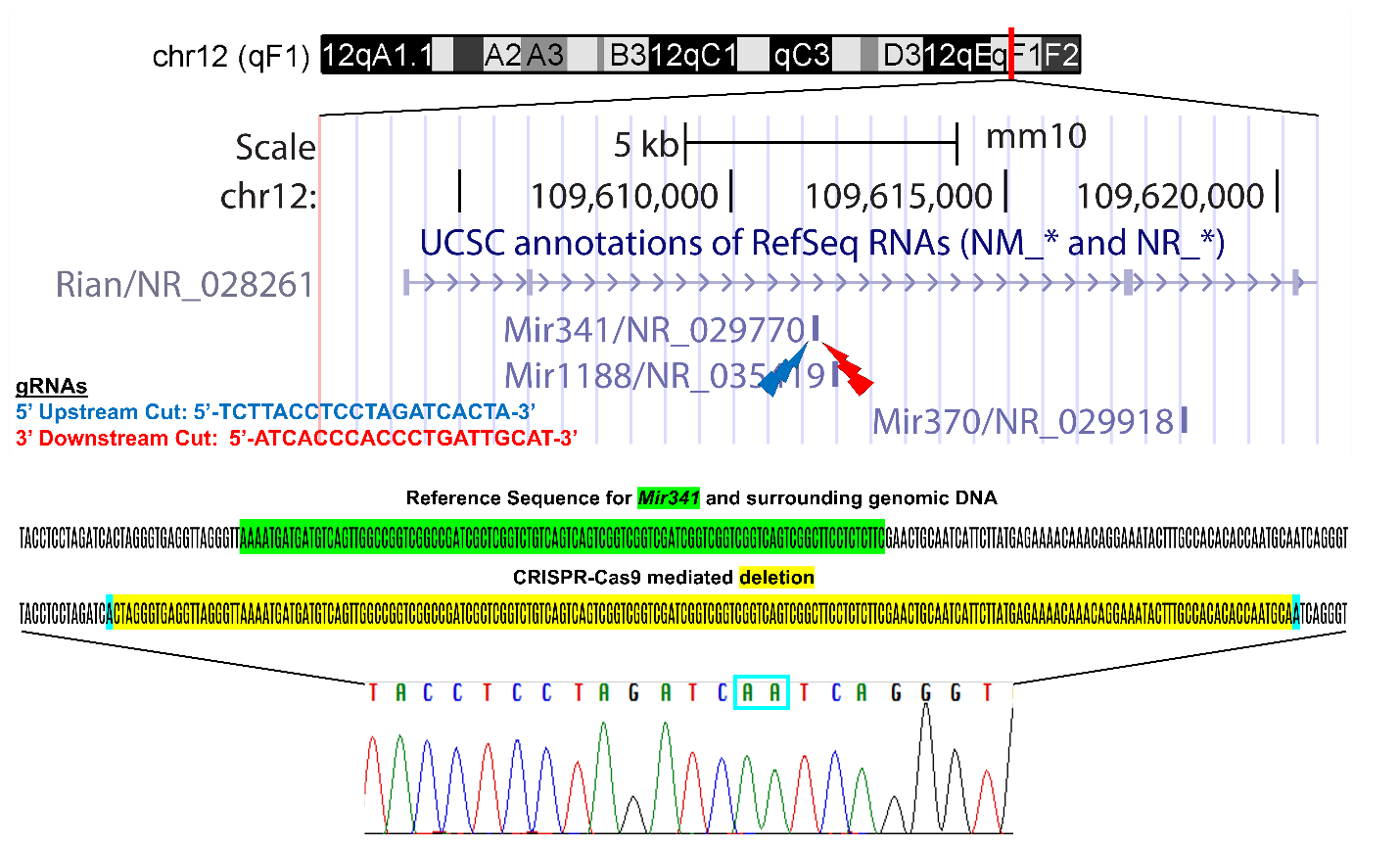

Retinal thickness of D2-*Mir341* mice at 18-22 wks old

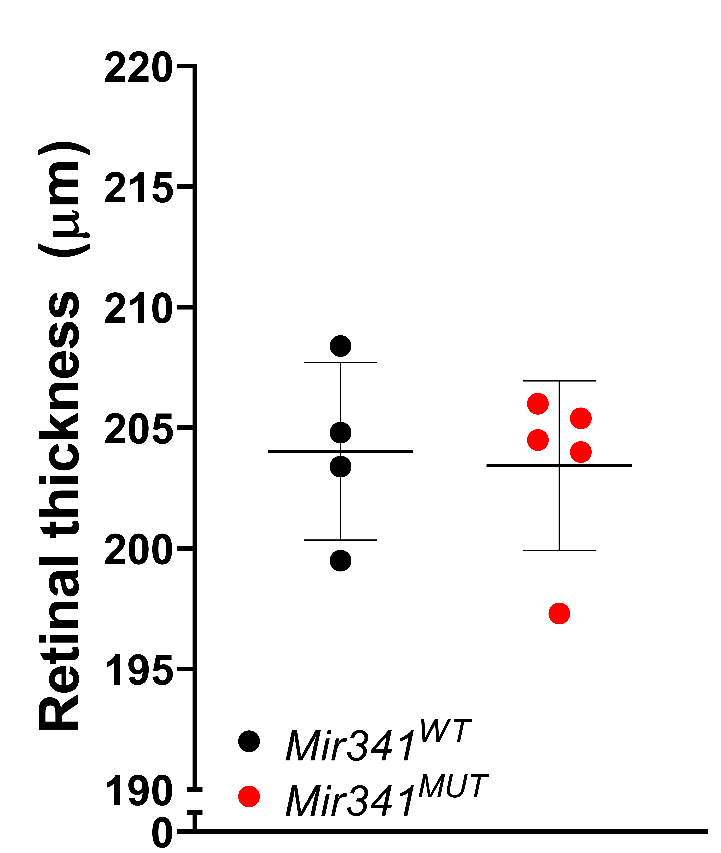
