## Supplementary Table 2: Mrdq3 Haplotype SNPs for "Genetic modifiers of *Cep290*-mediated retinal degeneration"

| Chr | Mb (GRCm38) | Gene | Consequence | dbSNP | Ref Allele | Alt Allele |
| --- | --- | --- | --- | --- | --- | --- |
| 14 | 52,104,761 | *Hnrnpc* | Intergenic; upstream | *rs31439328* | T | C |
| 14 | 52,105,217 | *Hnrnpc* | Intergenic; upstream | *rs234944185* | C | CTTTT |
| 14 | 52,105,571 | *Hnrnpc* | Intergenic; upstream | *rs226084803* | A | ACATATGCCGGG |
| 14 | 52,105,587 | *Hnrnpc* | Intergenic; upstream | *rs246764284* | C | T |
| 14 | 52,105,852 | *Hnrnpc* | Intergenic; upstream | *rs261443596* | T | C |
| 14 | 52,105,914 | *Hnrnpc* | Intergenic; upstream | *rs47155038* | A | G |
| 14 | 52,105,924 | *Hnrnpc* | Intergenic; upstream | *rs51130051* | A | G |
| 14 | 52,106,083 | *Hnrnpc* | Intergenic; upstream | *rs49455819* | A | T |
| 14 | 52,106,111 | *Hnrnpc* | Intergenic; upstream | *rs48676746* | T | C |
| 14 | 52,106,451 | *Hnrnpc* | Intergenic; upstream | *rs30310812* | G | T |
| 14 | 52,106,793 | *Hnrnpc* | Intergenic; upstream | *rs47739804* | G | C |
| 14 | 52,107,440 | *Hnrnpc* | Intergenic; upstream | *rs50542311* | C | G |
| 14 | 52,107,801 | *Hnrnpc* | Intergenic; upstream | *rs30309149* | C | T |
| 14 | 52,108,007 | *Hnrnpc* | Intergenic; upstream | *rs31539715* | C | T |
| 14 | 52,108,701 | *Hnrnpc* | Intergenic; upstream | *rs30452335* | T | A |
| 14 | 52,109,299 | *Rpgrip1* | Intergenic; upstream | *rs256484453* | G | A |
| 14 | 52,109,464 | *Rpgrip1* | Intergenic; upstream | *rs52359775* | A | G |
| 14 | 52,109,623 | *Rpgrip1* | Intergenic; upstream | *rs46670826* | T | A |
| 14 | 52,109,690 | *Rpgrip1* | Intergenic; upstream | *rs51301406* | G | C |
| 14 | 52,109,711 | *Rpgrip1* | Intergenic; upstream | *rs46268293* | A | G |
| 14 | 52,109,780 | *Rpgrip1* | Intergenic; upstream | *rs50564360* | G | C |
| 14 | 52,109,790 | *Rpgrip1* | Intergenic; upstream | *rs51475589* | G | C |
| 14 | 52,109,835 | *Rpgrip1* | Intergenic; upstream | *rs46188881* | G | A |
| 14 | 52,109,836 | *Rpgrip1* | Intergenic; upstream | *rs48898064* | A | G |
| 14 | 52,109,875 | *Rpgrip1* | Intergenic; upstream | *rs253898103* | T | c |
| 14 | 52,109,892 | *Rpgrip1* | Intergenic; upstream | *rs48018571* | C | t |
| 14 | 52,109,905 | *Rpgrip1* | Intergenic; upstream | *rs52119589* | T | A |
| 14 | 52,109,930 | *Rpgrip1* | Intergenic; upstream | *rs52295716* | C | T |
| 14 | 52,110,091 | *Rpgrip1* | Intergenic; upstream | *rs45997402* | A | C |
| Chr | Mb (GRCm38) | Gene | Consequence | dbSNP | Ref Allele | Alt Allele |
| 14 | 52,110,092 | *Rpgrip1* | Intergenic; upstream | *rs45994056* | G | A |
| 14 | 52,110,121 | *Rpgrip1* | Intergenic; upstream | *rs51160499* | C | T |
| 14 | 52,110,122 | *Rpgrip1* | Intergenic; upstream | *rs47556803* | T | C |
| 14 | 52,110,214 | *Rpgrip1* | Intergenic; upstream | *rs48882260* | A | C |
| 14 | 52,110,248 | *Rpgrip1* | Intergenic; upstream | *rs46061974* | A | G |
| 14 | 52,110,255 | *Rpgrip1* | Intergenic; upstream | *rs52564542* | C | T |
| 14 | 52,111,475 | *Rpgrip1* | Intron 1 | *rs52229469* | G | T |
| 14 | 52,111,765 | *Rpgrip1* | Intron 1 | *rs48960129* | A | G |
| 14 | 52,112,703 | *Rpgrip1* | Intron 2 | *rs4138897* | T | C |
| 14 | 52,112,786 | *Rpgrip1* | Intron 2 | *rs46479920* | T | G |
| 14 | 52,112,818 | *Rpgrip1* | Intron 2 | *rs264615719* | A | G |
| 14 | 52,113,675 | *Rpgrip1* | Intron 2 | *rs52315117* | C | A |
| 14 | 52,113,944 | *Rpgrip1* | Intron 2 | - | CCTGCCT | C |
| 14 | 52,114,138 | *Rpgrip1* | Intron 2 | *rs261665370* | A | AAAAC |
| 14 | 52,114,183 | *Rpgrip1* | Intron 2 | *rs264404745* | C | A |
| 14 | 52,114,320 | *Rpgrip1* | Intron 2 | *rs51627083* | C | T |
| 14 | 52,114,351 | *Rpgrip1* | Intron 2 | *rs48298743* | A | C |
| 14 | 52,114,601 | *Rpgrip1* | Intron 2 | *rs231443064* | A | AAAAT |
| 14 | 52,114,973 | *Rpgrip1* | Intron 3 | *rs31122581* | A | G |
| 14 | 52,115,045 | *Rpgrip1* | Intron 3 | *rs31230234* | G | A |
| 14 | 52,115,206 | *Rpgrip1* | Intron 3 | *rs244181953* | CGATTAGCATATCAGT | C |
| 14 | 52,115,261 | *Rpgrip1* | Intron 3 | *rs47511459* | A | T |
| 14 | 52,115,331 | *Rpgrip1* | Intron 3 | *rs213930478* | C | CCGAA |
| 14 | 52,115,340 | *Rpgrip1* | Intron 3 | *rs215741379* | G | A |
| 14 | 52,115,518 | *Rpgrip1* | Intron 3 | *rs49814691* | T | A |
| 14 | 52,115,519 | *Rpgrip1* | Intron 3 | *rs49302190* | C | G |
| 14 | 52,115,545 | *Rpgrip1* | Intron 3 | *rs46901350* | A | G |
| 14 | 52,115,784 | *Rpgrip1* | Intron 3 | *rs52054625* | T | G |
| 14 | 52,115,902 | *Rpgrip1* | Intron 3 | *rs51647175* | G | A |
| Chr | Mb (GRCm38) | Gene | Consequence | dbSNP | Ref Allele | Alt Allele |
| 14 | 52,116,153 | *Rpgrip1* | Intron 3 | *rs51222646* | A | T |
| 14 | 52,116,191 | *Rpgrip1* | Intron 3 | *rs51294582* | T | C |
| 14 | 52,116,442 | *Rpgrip1* | Intron 3 | *rs52533007* | T | C |
| 14 | 52,116,582 | *Rpgrip1* | Intron 3 | *rs31339756* | A | G |
| 14 | 52,116,621 | *Rpgrip1* | Intron 3 | *rs212144105* | T | TA |
| 14 | 52,116,851 | *Rpgrip1* | Intron 3 | *rs30314602* | A | G |
| 14 | 52,117,130 | *Rpgrip1* | Intron 3 | *rs247026625* | CT | C |
| 14 | 52,117,250 | *Rpgrip1* | Intron 3 | *rs226283181* | T | G |
| 14 | 52,117,254 | *Rpgrip1* | Intron 3 | *rs246455133* | T | C |
| 14 | 52,117,342 | *Rpgrip1* | Intron 3 | *rs47137849* | T | A |
| 14 | 52,117,455 | *Rpgrip1* | Intron 3 | *rs50738727* | T | A |
| 14 | 52,117,705 | *Rpgrip1* | Intron 3 | *rs52305207* | C | T |
| 14 | 52,117,823 | *Rpgrip1* | Intron 3 | *rs256879795* | T | G |
| 14 | 52,117,877 | *Rpgrip1* | Intron 3 | *rs48065287* | A | G |
| 14 | 52,117,918 | *Rpgrip1* | Intron 3 | *rs47247378* | C | G |
| 14 | 52,118,038 | *Rpgrip1* | Intron 3 | *rs260572179* | G | GCGA |
| 14 | 52,118,298 | *Rpgrip1* | Intron 3 | *rs50992653* | A | G |
| 14 | 52,118,517 | *Rpgrip1* | Intron 3 | *rs46547230* | G | A |
| 14 | 52,118,533 | *Rpgrip1* | Intron 3 | *rs47773474* | C | T |
| 14 | 52,118,551 | *Rpgrip1* | Intron 3 | *rs51939466* | C | A |
| 14 | 52,118,588 | *Rpgrip1* | Intron 3 | *rs48843053* | G | A |
| 14 | 52,118,707 | *Rpgrip1* | Intron 3 | *rs265013403* | AC | A |
| 14 | 52,118,747 | *Rpgrip1* | Intron 3 | *rs47362674* | G | A |
| 14 | 52,118,753 | *Rpgrip1* | Intron 3 | *rs52099118* | G | A |
| 14 | 52,118,776 | *Rpgrip1* | Intron 3 | *rs52512820* | A | T |
| 14 | 52,118,780 | *Rpgrip1* | Intron 3 | *rs52257045* | A | G |
| 14 | 52,118,848 | *Rpgrip1* | Intron 3 | *rs52582328* | A | T |
| 14 | 52,118,898 | *Rpgrip1* | Intron 3 | *rs47898394* | G | A |
| Chr | Mb (GRCm38) | Gene | Consequence | dbSNP | Ref Allele | Alt Allele |
| 14 | 52,119,037 | *Rpgrip1* | Intron 3 | *rs3679692* | T | A |
| 14 | 52,119,267 | *Rpgrip1* | Exon 4; Missense (P96L) | *rs3681109* | C | T |
| 14 | 52,119,308 | *Rpgrip1* | Exon 4; Missense (P110S) | *rs3681193* | C | T |
| 14 | 52,119,325 | *Rpgrip1* | Exon 4; Synonymous | *rs3681636* | C | T |
| 14 | 52,119,328 | *Rpgrip1* | Exon 4; Synonymous | *rs3681648* | A | C |
| 14 | 52,119,390 | *Rpgrip1* | Exon 4; Missense (A137V) | *rs3681773* | C | T |
| 14 | 52,119,527 | *Rpgrip1* | Intron 4 | *rs47589373* | A | G |
| 14 | 52,119,700 | *Rpgrip1* | Intron 4 | *rs52301489* | G | C |
| 14 | 52,119,834 | *Rpgrip1* | Intron 4 | *rs52041997* | A | G |
| 14 | 52,119,844 | *Rpgrip1* | Intron 4 | *rs47194128* | G | A |
| 14 | 52,119,998 | *Rpgrip1* | Intron 4 | *rs46511628* | T | C |
| 14 | 52,120,122 | *Rpgrip1* | Intron 4 | *rs51097614* | A | G |
| 14 | 52,120,522 | *Rpgrip1* | Exon 5; Synonymous | *rs47971646* | T | C |
| 14 | 52,120,786 | *Rpgrip1* | Intron 5 | *rs222105764* | T | TGTGA |
| 14 | 52,120,906 | *Rpgrip1* | Intron 5 | *rs30786861* | C | T |
| 14 | 52,120,924 | *Rpgrip1* | Intron 5 | *rs31244416* | G | T |
| 14 | 52,121,175 | *Rpgrip1* | Intron 6 | *rs31107077* | T | C |
| 14 | 52,121,186 | *Rpgrip1* | Intron 6 | *rs31481343* | C | G |
| 14 | 52,121,238 | *Rpgrip1* | Intron 6 | *rs31276246* | T | C |
| 14 | 52,121,793 | *Rpgrip1* | Intron 6 | *rs45698261* | G | A |
| 14 | 52,121,879 | *Rpgrip1* | Intron 6 | *rs47360380* | T | G |
| 14 | 52,122,017 | *Rpgrip1* | Intron 6 | *rs52523874* | G | A |
| 14 | 52,122,097 | *Rpgrip1* | Intron 6 | *rs52159070* | C | T |
| 14 | 52,122,373 | *Rpgrip1* | Intron 6 | *rs262329114* | AT | A |
| 14 | 52,122,760 | *Rpgrip1* | Intron 6 | *rs46525582* | C | G |
| 14 | 52,122,777 | *Rpgrip1* | Intron 6 | *rs48536021* | T | C |
| 14 | 52,123,107 | *Rpgrip1* | Intron 6 | *rs52608008* | T | G |
| 14 | 52,123,160 | *Rpgrip1* | Intron 6 | *rs251693236* | A | G |
| 14 | 52,123,332 | *Rpgrip1* | Intron 6 | *rs213809743* | G | A |
| Chr | Mb (GRCm38) | Gene | Consequence | dbSNP | Ref Allele | Alt Allele |
| 14 | 52,123,677 | *Rpgrip1* | Intron 6 | *rs30922501* | T | C |
| 14 | 52,124,159 | *Rpgrip1* | Intron 6 | *rs251731361* | A | ACTTTGT |
| 14 | 52,124,281 | *Rpgrip1* | Intron 6 | *rs50910977* | A | T |
| 14 | 52,124,290 | *Rpgrip1* | Intron 6 | *rs47897067* | T | C |
| 14 | 52,124,298 | *Rpgrip1* | Intron 6 | *rs215252352* | TGAGTGCAAGGCCAGCCTAGAG | T |
| 14 | 52,124,327 | *Rpgrip1* | Intron 6 | *rs219179190* | C | T |
| 14 | 52,124,369 | *Rpgrip1* | Intron 6 | *rs47987041* | C | A |
| 14 | 52,124,391 | *Rpgrip1* | Intron 6 | *rs108351655* | G | A |
| 14 | 52,124,396 | *Rpgrip1* | Intron 6 | *rs51296173* | C | T |
| 14 | 52,124,412 | *Rpgrip1* | Intron 6 | *rs46235848* | C | G |
| 14 | 52,124,699 | *Rpgrip1* | Intron 6 | *rs48373492* | G | A |
| 14 | 52,125,493 | *Rpgrip1* | Intron 6 | *rs387477272* | T | C |
| 14 | 52,125,995 | *Rpgrip1* | Intron 6 | *rs48163771* | T | G |
| 14 | 52,126,016 | *Rpgrip1* | Intron 6 | *rs51342684* | A | G |
| 14 | 52,127,115 | *Rpgrip1* | Intron 7 | *rs31019226* | T | C |
| 14 | 52,127,128 | *Rpgrip1* | Intron 7 | *rs31506515* | A | G |
| 14 | 52,127,706 | *Rpgrip1* | Intron 7 | *rs46567233* | T | C |
| 14 | 52,128,754 | *Rpgrip1* | Intron 7 | *rs222386295* | G | C |
| 14 | 52,128,987 | *Rpgrip1* | Intron 7 | *rs51264018* | A | G |
| 14 | 52,129,064 | *Rpgrip1* | Intron 7 | *rs49669988* | G | A |
| 14 | 52,129,110 | *Rpgrip1* | Intron 7 | *rs51094753* | G | A |
| 14 | 52,129,186 | *Rpgrip1* | Intron 7 | *rs50012539* | A | C |
| 14 | 52,129,188 | *Rpgrip1* | Intron 7 | *rs48126763* | G | A |
| 14 | 52,129,248 | *Rpgrip1* | Intron 7 | *rs47848574* | A | G |
| 14 | 52,129,447 | *Rpgrip1* | Intron 7 | *rs248697233* | GAGAT | G |
| 14 | 52,129,676 | *Rpgrip1* | Intron 7 | *rs51012611* | C | A |
| 14 | 52,129,826 | *Rpgrip1* | Intron 7 | *rs50209346* | G | A |
| 14 | 52,129,930 | *Rpgrip1* | Intron 7 | *rs51177150* | T | C |
| 14 | 52,129,933 | *Rpgrip1* | Intron 7 | *rs50654880* | C | G |
