## Supplementary Table 1: Differentially expressed Mrdq positional candidate genes for "Genetic modifiers of *Cep290*-mediated retinal degeneration"

**Supplementary Table 1.** Mrdq positional candidate genes with significant expression differences in P25 mouse retina between BXD24 and CAST

| **QTL^a^** | **Gene** | **Log_2_ FC^b^** | ***p*-value** | **FDR^c^** | **BXD24 CPM^d^** | **CAST CPM^d^** |
| --- | --- | --- | --- | --- | --- | --- |
| *Mrdq1* | *Meg3* | -1.03 | 1.99E^-3^ | 9.78E^-3^ | 722.14 | 340.62 |
|  | *Lgmn* | -0.61 | 8.07E^-5^ | 7.52E^-4^ | 230.82 | 150.80 |
|  | *Syne3* | 2.94 | 1.06E^-5^ | 1.44E^-4^ | 48.41 | 16.48 |
|  | *Moap1* | 0.68 | 1.10E^-2^ | 3.73E^-2^ | 12.83 | 20.50 |
|  | *Ccdc88c* | -1.16 | 8.50E^-6^ | 3.73E^-2^ | 15.97 | 7.15 |
|  | *Serpina3h* | -3.71 | 8.68E^-12^ | 2.87E^-9^ | 13.61 | 1.03 |
|  | *Asb2* | 0.83 | 2.33E^-5^ | 2.76E^-4^ | 5.74 | 10.22 |
|  | *Fbln5* | -0.79 | 2.63E^-5^ | 3.05E^-4^ | 5.08 | 2.93 |
|  | *9030617O03Rik* | 1.20 | 7.60E^-6^ | 1.10E^-4^ | 2.34 | 5.34 |
|  | *Gm20604* | 0.55 | 1.57E^-3^ | 8.08E^-3^ | 16.17 | 23.59 |
|  | *mmu-miR-541-5p* | 0.69 | 2.64E^-8^ | N/A | 2,380.86 | 3,840.36 |
|  | *mmu-miR-341-3p* | -6.19 | 1.60E^-119^ | N/A | 584.58 | 7.02 |
|  | *mmu-miR-431-5p* | 1.53 | 1.13E^-12^ | N/A | 126.56 | 372.35 |
|  | *mmu-miR-540-3p* | 1.03 | 1.90E^-7^ | N/A | 57.47 | 118.14 |
|  | *mmu-miR-376a-5p* | 0.62 | 5.75E^-3^ | N/A | 70.55 | 107.92 |
|  | *mmu-miR-673-5p* | -3.46 | 7.55E^-21^ | N/A | 62.10 | 4.95 |
|  | *mmu-miR-376b-5p* | 0.50 | 1.20E^-2^ | N/A | 67.03 | 94.51 |
|  | *mmu-miR-544-3p* | -1.12 | 1.12E^-2^ | N/A | 24.29 | 10.70 |
| *Mrdq2* | *Elovl2* | 1.84 | 5.97E^-10^ | 6.53E^-8^ | 65.76 | 236.25 |
|  | *Hist1h1c* | 1.13 | 1.67E^-5^ | 2.07E^-4^ | 96.13 | 209.23 |
|  | *Dtnbp1* | 0.59 | 7.03E^-5^ | 6.76E^-4^ | 44.87 | 67.41 |
|  | *Dcdc2a* | 1.65 | 1.59E^-6^ | 2.99E^-5^ | 11.19 | 35.16 |
|  | *Tbc1d7* | 0.65 | 7.06E^-5^ | 6.78E^-4^ | 17.50 | 27.51 |
|  | *Dsp* | 1.77 | 7.16E^-9^ | 3.95E^-7^ | 7.93 | 27.21 |
|  | *Fars2* | 1.11 | 5.09E^-9^ | 3.07E^-7^ | 12.10 | 26.15 |
|  | *Mrs2* | -0.74 | 1.59E^-3^ | 8.18E^-3^ | 21.07 | 12.57 |
|  | *Ripk1* | 0.62 | 5.38E^-5^ | 5.45E^-4^ | 9.38 | 14.39 |
|  | *Nqo2* | -1.08 | 6.61E^-6^ | 9.80E^-5^ | 13.32 | 6.25 |
|  | *Serpinb6a* | -0.60 | 2.52E^-3^ | 1.18E^-2^ | 12.89 | 8.52 |
|  | *Mboat1* | 1.06 | 1.11E^-4^ | 9.86E^-4^ | 5.23 | 10.92 |
|  | *Sirt5* | 0.70 | 6.67E^-5^ | 6.48E^-4^ | 6.60 | 10.72 |
|  | *Hist1h2ac* | 1.32 | 4.22E^-5^ | 4.51E^-4^ | 3.89 | 9.70 |
|  | *Riok* | 0.55 | 1.13E^-5^ | 1.52E^-4^ | 16.94 | 24.74 |
| *Mrdq3* | *Rpgrip1* | -0.57 | 1.02E^-2^ | 3.5E^-2^ | 732.11 | 476.54 |
|  | *Gm26782* | 0.98 | 4.73E^-5^ | 4.94E^-4^ | 9.37 | 18.15 |
|  | *Tep1* | -0.88 | 2.39E^-4^ | 1.81E^-3^ | 9.72 | 18.05 |
| *Mrdq4* | *Ttc39c* | 1.05 | 9.04E^-5^ | 8.29E^-4^ | 45.03 | 93.19 |
| **QTL^a^** | **Gene** | **Log_2_ FC^b^** | ***p*-value** | **FDR^c^** | **BXD24 CPM^d^** | **CAST CPM^d^** |
| *Mrdq4* | *Zfp521* | -0.53 | 9.56E^-6^ | 5.43E^-3^ | 46.97 | 32.49 |
|  | *Ttr* | 1.20 | 1.25E^-6^ | 2.48E^-5^ | 792.92 | 1795.39 |
|  | *Sft2d3* | -0.57 | 6.10E^-5^ | 6.06E^-4^ | 17.36 | 11.69 |
|  | *Gm16344* | 1.29 | 3.27E^-4^ | 2.34E^-3^ | 9.26 | 22.48 |
|  | *Stard4* | -0.72 | 4.22E^-3^ | 1.74E^-2^ | 39.85 | 24.28 |
|  | *Epb41l4aos* | -1.00 | 2.10E^-5^ | 2.53E^-4^ | 6.45 | 3.19 |
|  | *Apc* | -0.51 | 1.20E^-2^ | 3.97E^-2^ | 349.54 | 242.14 |
|  | *Fam53c* | -0.58 | 3.14E^-5^ | 3.54E^-4^ | 112.00 | 74.90 |
|  | *Egr1* | 1.67 | 1.80E^-4^ | 1.44E^-3^ | 18.81 | 59.95 |
|  | *Hbegf* | 0.54 | 1.64E^-4^ | 1.34E^-3^ | 12.36 | 17.98 |
|  | *Gm10544* | 1.96 | 1.96E^-8^ | 8.84E^-7^ | 3.31 | 12.68 |
|  | *Pcdhb3* | -0.83 | 1.07E^-4^ | 9.53E^-4^ | 9.73 | 5.45 |
|  | *Pcdhb6* | 0.60 | 4.41E^-3^ | 1.80E^-2^ | 6.16 | 9.37 |
|  | *Pcdhb8* | -0.95 | 3.18E^-5^ | 3.57E^-4^ | 7.58 | 3.91 |
|  | *Pcdhb9* | -1.18 | 3.64E^-7^ | 9.05E^-6^ | 22.56 | 9.94 |
|  | *Pcdhb10* | -0.76 | 8.18E^-4^ | 4.80E^-3^ | 13.80 | 8.15 |
|  | *Pcdhb11* | -1.12 | 9.85E^-6^ | 1.35E^-4^ | 10.72 | 4.93 |
|  | *Pcdhb14* | -1.32 | 2.22E^-5^ | 2.66E^-4^ | 11.98 | 4.82 |
|  | *Pcdhb15* | -2.12 | 6.80E^-6^ | 9.99E^-5^ | 6.11 | 1.40 |
|  | *Pcdhb16* | -1.78 | 4.14E^-5^ | 4.45E^-4^ | 15.80 | 4.57 |
|  | *Pcdhb17* | -1.17 | 6.27E^-7^ | 1.41E^-5^ | 25.72 | 11.39 |
|  | *Pcdhb18* | -0.71 | 9.55E^-3^ | 3.34E^-2^ | 14.57 | 8.79 |
|  | *Pcdhb19* | -1.72 | 1.52E^-5^ | 1.93E^-4^ | 18.28 | 5.46 |
|  | *Pcdhb20* | -1.11 | 5.05E^-7^ | 1.19E^-5^ | 18.84 | 8.74 |
|  | *Pcdhb21* | -1.67 | 2.82E^-5^ | 3.23E^-4^ | 6.91 | 2.14 |
|  | *Pcdhb22* | -0.65 | 2.35E^-4^ | 1.78E^-3^ | 13.34 | 8.49 |
|  | *0610009O20Rik* | 0.53 | 2.79E^-6^ | 4.80E^-5^ | 55.07 | 79.67 |
|  | *Spry4* | 0.84 | 7.10E^-5^ | 6.81E^-4^ | 13.22 | 23.69 |
|  | *Dpysl3* | -0.88 | 8.91E^-7^ | 1.87E^-5^ | 351.73 | 191.17 |
|  | *Jakmip2* | -0.72 | 1.48E^-4^ | 1.25E^-3^ | 21.89 | 13.30 |
|  | *Atg12* | 0.57 | 1.14E^-5^ | 1.52E^-4^ | 67.86 | 100.45 |
|  | *Snx24* | 0.63 | 4.80E^-3^ | 1.93E^-2^ | 17.31 | 26.72 |
|  | *Aldh7a1* | -0.70 | 6.09E^-7^ | 1.38E^-5^ | 68.53 | 42.30 |
|  | *C330018D20Rik* | 0.83 | 1.02E^-5^ | 1.40E^-4^ | 8.92 | 15.82 |
|  | *Slc27a6* | 1.05 | 1.77E^-4^ | 1.42E^-3^ | 2.50 | 5.26 |
|  | *Ndst1* | -0.67 | 5.26E^-4^ | 3.36E^-3^ | 155.77 | 98.63 |
|  | *Pdgfrb* | -0.54 | 1.00E^-3^ | 5.65E^-3^ | 20.02 | 13.79 |
|  | *Slc26a2* | -0.61 | 1.45E^-3^ | 7.58E^-3^ | 69.14 | 44.44 |
|  | *Rps2-ps10* | -5.87 | 8.07E^-8^ | 2.72E^-6^ | 18.43 | 0.34 |
|  | *Ppargc1b* | -2.80 | 1.74E^-8^ | 8.08E^-7^ | 85.58 | 12.23 |
|  | *Afap1l1* | -1.48 | 8.44E^-8^ | 2.81E^-6^ | 12.62 | 4.54 |
|  | *Onecut2* | -0.84 | 5.84E^-3^ | 2.26E^-2^ | 9.49 | 5.22 |
|  | *Amd-ps3* | 1.70 | 2.46E^-3^ | 1.15E^-2^ | 2.45 | 8.15 |
| **QTL^a^** | **Gene** | **Log_2_ FC^b^** | ***p*-value** | **FDR^c^** | **BXD24 CPM^d^** | **CAST CPM^d^** |
| *Mrdq4* | *Tubb6* | 1.14 | 6.94E^-6^ | 1.02E^-4^ | 3.52 | 7.66 |
|  | *Ldlrad4* | -0.74 | 8.76E^-3^ | 3.13E^-2^ | 14.06 | 8.34 |
|  | *Tcf4* | -0.58 | 2.30E^-3^ | 1.10E^-2^ | 103.15 | 68.66 |
|  | *Poli* | 1.00 | 1.29E^-7^ | 3.92E^-6^ | 11.06 | 22.14 |
|  | *Lipg* | 0.66 | 2.15E^-4^ | 1.66E^-3^ | 4.18 | 6.60 |
|  | *Katnal2* | -0.80 | 1.04E^-3^ | 5.81E^-3^ | 10.83 | 6.21 |
|  | *Slc14a1* | -0.64 | 4.60E^-4^ | 3.03E^-3^ | 12.83 | 8.31 |
|  | *Hsbp1l1* | -0.87 | 2.35E^-5^ | 2.78E^-4^ | 29.02 | 15.95 |
|  | *Gm17383* | -4.10 | 1.00E^-7^ | 3.19E^-6^ | 116.25 | 6.75 |
|  | *Mbp* | -0.96 | 1.61E^-4^ | 1.32E^-3^ | 34.63 | 17.91 |
|  | *Zfp236* | -1.38 | 4.94E^-8^ | 1.87E^-6^ | 94.73 | 36.07 |
|  | *Zfp516* | -0.51 | 6.27E^-4^ | 3.88E^-3^ | 180.72 | 126.73 |
|  | *Cyb5a* | 0.70 | 4.34E^-5^ | 4.62E^-4^ | 23.92 | 38.59 |
|  | *mmu-miR-1258-3p* | 3.84 | 3.63E^-6^ | N/A | 0.00 | 8.60 |
| *Mrdq5* | *Cntnap5a* | -0.78 | 1.51E^-2^ | 4.72E^-2^ | 11.05 | 6.22 |
|  | *Nifk* | -0.51 | 1.53E^-4^ | 1.27E^-3^ | 34.01 | 23.74 |
|  | *Dbi* | 1.00 | 1.32E^-5^ | 1.71E^-4^ | 64.04 | 126.58 |
|  | *C1ql2* | 0.55 | 1.71E^-4^ | 1.39E^-3^ | 73.81 | 107.46 |
|  | *Ccdc93* | -0.55 | 8.06E^-4^ | 4.74E^-3^ | 30.66 | 20.90 |
|  | *Mgat5* | -0.97 | 9.69E^-4^ | 5.49E^-3^ | 58.18 | 30.02 |
|  | *Rpl28-ps1* | -9.51 | 3.68E^-10^ | 4.4E^-8^ | 139.42 | 0.19 |
|  | *Mcm6* | 1.83 | 1.54E^-7^ | 4.52E^-6^ | 1.49 | 5.25 |
|  | *AA986860* | -1.77 | 4.77E^-8^ | 1.81E^-6^ | 8.54 | 2.50 |
|  | *Mfsd4* | 0.60 | 2.46E^-4^ | 1.85E^-3^ | 49.11 | 74.15 |
|  | *Nfasc* | -0.57 | 7.82E^-4^ | 4.62E^-3^ | 371.94 | 247.77 |
|  | *Mdm4* | -0.52 | 8.27E^-3^ | 2.99E^-2^ | 37.53 | 25.46 |
|  | *Gm38394* | -2.15 | 9.97E^-8^ | 3.17E^-6^ | 180.50 | 39.84 |
|  | *Prelp* | -0.86 | 4.94E^-3^ | 1.98E^-2^ | 47.14 | 25.97 |
|  | *Cyb5r1* | 0.60 | 7.34E^-2^ | 4.39E^-3^ | 33.25 | 50.00 |
|  | *Adipor1* | 0.78 | 1.12E^-5^ | 1.51E^-4^ | 495.78 | 848.05 |
|  | *Gm15454* | 8.14 | 8.92E^-12^ | 2.87E^-9^ | 0.09 | 25.28 |
|  | *Kdm5b* | 0.53 | 1.90E^-4^ | 1.51E^-3^ | 203.61 | 294.62 |
|  | *Syt2* | -0.58 | 2.65E^-5^ | 3.06E^-4^ | 36.54 | 24.44 |
|  | *Shisa4* | 1.79 | 1.07E^-8^ | 5.53E^-7^ | 22.49 | 77.51 |
|  | *Pkp1* | -2.09 | 6.72E^-8^ | 2.37E^-6^ | 6.28 | 1.49 |
|  | *Igfn1* | -1.99 | 1.25E^-7^ | 3.81E^-6^ | 70.29 | 17.08 |
|  | *Gm15850* | 1.25 | 2.04E^-4^ | 1.60E^-3^ | 3.43 | 8.07 |
|  | *B3galt2* | -0.60 | 4.42E^-3^ | 1.80E^-2^ | 82.13 | 54.19 |
|  | *Rgs2* | 0.75 | 1.06E^-5^ | 1.44E^-4^ | 35.94 | 59.95 |
|  | *Pdc* | 0.81 | 8.88E^-4^ | 5.12E^-3^ | 1840.88 | 3227.92 |
|  | *Npl* | 0.68 | 1.62E^-5^ | 2.03E^-4^ | 15.97 | 25.39 |
|  | *BC034090* | -0.78 | 9.69E^-6^ | 1.33E^-4^ | 191.21 | 111.44 |
| **QTL^a^** | **Gene** | **Log_2_ FC^b^** | ***p*-value** | **FDR^c^** | **BXD24 CPM^d^** | **CAST CPM^d^** |
| *Mrdq5* | *Gm2000* | 0.63 | 5.20E^-3^ | 2.06E^-2^ | 15.09 | 23.04 |
|  | *BC026585* | 1.61 | 2.03E^-8^ | 9.03E^-7^ | 1.93 | 5.90 |
|  | *Pappa2* | 0.92 | 1.72E^-5^ | 2.12E^-4^ | 11.62 | 21.81 |
|  | *Tnr* | -1.32 | 3.87E^-3^ | 1.63E^-2^ | 38.56 | 15.51 |
|  | *Fmo1* | -5.58 | 6.35E^-11^ | 1.22E^-8^ | 54.76 | 1.14 |
|  | *Slc19a2* | 1.23 | 3.55E^-6^ | 5.81E^-5^ | 12.48 | 29.38 |
|  | *Ccdc181* | 0.57 | 7.35E^-4^ | 4.39E^-3^ | 29.73 | 44.09 |
|  | *Sft2d2* | -0.55 | 4.24E^-4^ | 2.87E^-3^ | 24.57 | 16.71 |
|  | *Pou2f1* | -0.79 | 1.51E^-3^ | 7.86E^-3^ | 100.83 | 57.33 |
|  | *Gm16701* | 1.45 | 1.31E^-7^ | 3.96E^-6^ | 4.61 | 12.58 |
|  | *Rxrg* | 1.25 | 3.04E^-6^ | 5.13E^-5^ | 3.62 | 8.63 |
|  | *Rgs5* | -1.95 | 7.34E^-9^ | 4.03E^-7^ | 75.86 | 19.70 |
|  | *Rgs4* | -0.70 | 1.15E^-5^ | 1.53E^-4^ | 100.92 | 62.14 |
|  | *Hsd17b7* | -0.59 | 4.79E^-4^ | 3.13E^-3^ | 13.45 | 8.93 |
|  | *Uhmk1* | -0.59 | 1.26E^-2^ | 4.13E^-2^ | 371.57 | 247.11 |
|  | *Pcp4l1* | 0.75 | 3.40E^-4^ | 2.41E^-3^ | 46.21 | 77.25 |
|  | *Klhdc9* | 1.87 | 7.99E^-8^ | 2.70E^-6^ | 1.51 | 5.53 |
|  | *F11r* | -1.40 | 7.6E^-5^ | 7.18E^-4^ | 5.97 | 2.25 |
|  | *Alyref2* | -0.93 | 1.79E^-4^ | 1.43E^-3^ | 11.60 | 6.08 |
|  | *Atp1a2* | -2.19 | 1.51E^-8^ | 7.20E^-7^ | 76.98 | 17.04 |
|  | *Kcnj9* | -0.53 | 3.72E^-5^ | 4.06E^-4^ | 95.26 | 65.96 |
|  | *Dusp23* | -0.93 | 1.66E^-4^ | 1.36E^-3^ | 5.93 | 3.11 |
|  | *Kif26b* | -1.44 | 6.28E^-5^ | 6.17E^-4^ | 8.67 | 3.18 |
|  | *Sccpdh* | 0.65 | 3.98E^-4^ | 2.74E^-3^ | 47.43 | 74.38 |
|  | *Psen2* | -0.54 | 5.58E^-4^ | 3.53E^-3^ | 22.28 | 15.35 |
|  | *6330403A02Rik* | -0.76 | 2.29E^-6^ | 4.13E^-5^ | 20.23 | 11.95 |
|  | *Ephx1* | 1.12 | 7.64E^-8^ | 2.60E^-6^ | 13.46 | 29.01 |
|  | *Disp1* | 1.37 | 9.19E^-7^ | 1.93E^-5^ | 6.26 | 16.22 |
|  | *C130074G19Rik* | -0.88 | 4.51E^-4^ | 3.00E^-3^ | 5.75 | 3.12 |
| *Mrdq6* | *Gm13251* | 1.02 | 3.31E^-4^ | 2.36E^-3^ | 3.87 | 7.82 |
|  | *Gm13157* | -7.20 | 1.17E^-9^ | 1.01E^-7^ | 5.82 | 0.04 |
|  | *Zfp933* | -0.72 | 2.47E^-5^ | 2.89E^-4^ | 21.65 | 13.04 |
|  | *Miip* | -1.00 | 6.12E^-6^ | 9.17E^-5^ | 21.35 | 10.66 |
|  | *Fbxo6* | 0.59 | 1.21E^-4^ | 1.06E^-3^ | 14.94 | 22.31 |
|  | *Apitd1* | 3.13 | 1.68E^-9^ | 1.32E^-7^ | 1.97 | 17.12 |
|  | *H6pd* | 0.55 | 1.73E^-4^ | 1.40E^-3^ | 6.26 | 9.16 |
|  | *Park7* | 0.70 | 3.71E^-5^ | 4.05E^-4^ | 104.15 | 167.32 |
| *Mrdq7* | *Ets1* | -1.04 | 3.54E^-4^ | 2.49E^-3^ | 6.03 | 2.94 |
|  | *Gm10698* | -8.10 | 3.09E^-10^ | 3.96E^-8^ | 18.81 | 0.06 |
|  | *Fez1* | 0.51 | 2.29E^-4^ | 1.75E^-3^ | 67.15 | 95.29 |
|  | *Gm10177* | -7.12 | 1.62E^-10^ | 2.29E^-8^ | 23.08 | 0.16 |
| **QTL^a^** | **Gene** | **Log_2_ FC^b^** | ***p*-value** | **FDR^c^** | **BXD24 CPM^d^** | **CAST CPM^d^** |
| *Mrdq7* | *Ccdc15* | -1.04 | 2.53E^-3^ | 1.18E^-2^ | 5.30 | 2.57 |
|  | *Hepacam* | 1.29 | 3.29E^-4^ | 2.36E^-3^ | 4.19 | 10.28 |
|  | *Vwa5a* | -0.74 | 1.66E^-4^ | 1.36E^-3^ | 13.43 | 8.09 |
|  | *Ubash3b* | 0.80 | 5.09E^-7^ | 1.20E^-5^ | 23.77 | 41.48 |
|  | *Arhgef12* | -0.51 | 4.91E^-3^ | 1.96E^-2^ | 251.84 | 175.72 |
|  | *Rnf26* | -0.51 | 2.80E^-5^ | 3.22E^-4^ | 32.95 | 23.10 |
|  | *Mcam* | 0.97 | 1.97E^-6^ | 3.61E^-5^ | 5.86 | 11.46 |
|  | *C030014I23Rik* | -0.90 | 7.58E^-5^ | 7.16E^-4^ | 8.04 | 4.32 |
|  | *Ddx6* | -0.65 | 4.97E^-4^ | 3.22E^-3^ | 271.06 | 172.61 |
|  | *Fxyd6* | -1.03 | 5.62E^-7^ | 1.29E^-5^ | 40.26 | 19.65 |
|  | *Gm5617* | 0.77 | 1.72E^-3^ | 8.70E^-3^ | 5.64 | 9.45 |
|  | *Zbtb16* | -1.11 | 1.90E^-3^ | 9.43E^-3^ | 5.59 | 2.58 |
|  | *Htr3a* | -0.71 | 4.00E^-5^ | 4.31E^-4^ | 17.81 | 10.76 |
|  | *Ttc12* | -0.79 | 1.20E^-5^ | 1.58E^-4^ | 8.50 | 4.93 |
|  | *Bco2* | 4.06 | 8.46E^-10^ | 8.07E^-8^ | 0.64 | 10.81 |
|  | *Dixdc1* | -0.74 | 2.45E^-6^ | 4.35E^-5^ | 155.45 | 93.07 |
|  | *Cryab* | 1.29 | 2.56E^-3^ | 1.19E^-2^ | 78.25 | 186.48 |
|  | *Ppp2r1b* | 0.64 | 2.92E^-4^ | 2.13E^-3^ | 14.31 | 22.22 |
|  | *AI593442* | -0.94 | 3.32E^-6^ | 5.5E^-5^ | 60.92 | 31.80 |
|  | *Dmxl2* | -0.57 | 3.14E^-3^ | 1.39E^-2^ | 358.41 | 240.72 |
|  | *Cib2* | -0.71 | 3.95E^-5^ | 4.27E^-4^ | 13.38 | 8.13 |
|  | *Crabp1* | 0.81 | 1.58E^-4^ | 1.31E^-3^ | 9.81 | 17.02 |
|  | *Chrna3* | 0.71 | 4.93E^-4^ | 3.20E^-3^ | 11.79 | 19.22 |
|  | *Chrnb4* | 1.17 | 1.06E^-5^ | 1.44E^-4^ | 8.36 | 18.72 |
|  | *mmu-miR-6236* | 0.81 | 4.46E^-3^ | N/A | 43.36 | 76.37 |
| *Mrdq8* | *Ppp1r14c* | 0.58 | 4.63E^-4^ | 3.05E^-3^ | 5.77 | 8.62 |
|  | *Plekhg1* | 2.53 | 6.45E^-11^ | 1.22E^-8^ | 15.92 | 91.65 |
|  | *Rmnd1* | -1.07 | 6.55E^-8^ | 2.33E^-6^ | 34.09 | 16.16 |
|  | *Vip* | -0.58 | 9.27E^-3^ | 3.27E^-2^ | 13.49 | 8.97 |
|  | *Epm2a* | 0.58 | 4.12E^-3^ | 1.71E^-2^ | 41.35 | 61.91 |
|  | *Utrn* | -0.91 | 7.35E^-5^ | 6.99E^-4^ | 83.12 | 44.11 |
|  | *Sf3b5* | 0.51 | 8.74E^-3^ | 3.13E^-2^ | 21.05 | 29.60 |
|  | *Fuca2* | 0.55 | 3.55E^-5^ | 3.91E^-4^ | 52.48 | 76.80 |
|  | *Adgrg6* | 1.35 | 1.59E^-6^ | 2.99E^-5^ | 2.00 | 5.08 |
|  | *Ccdc28a* | 0.61 | 5.02E^-5^ | 5.18E^-4^ | 7.29 | 11.19 |
|  | *Nhsl1* | 0.63 | 1.71E^-3^ | 8.65E^-3^ | 6.18 | 9.59 |
|  | *Tnfaip3* | -0.72 | 1.39E^-6^ | 2.70E^-5^ | 363.32 | 221.34 |
|  | *Myb* | -1.05 | 2.10E^-4^ | 1.63E^-3^ | 4.92 | 2.36 |
|  | *Slc2a12* | 1.32 | 8.41E^-8^ | 2.80E^-6^ | 7.41 | 18.69 |
|  | *Rps12* | 1.24 | 6.47E^-5^ | 6.33E^-4^ | 66.14 | 152.85 |
|  | *Ctgf* | 1.61 | 4.67E^-5^ | 4.89E^-4^ | 23.59 | 72.04 |
|  | *Lama2* | 1.49 | 4.35E^-7^ | 1.05E^-5^ | 2.78 | 7.86 |
| **QTL^a^** | **Gene** | **Log_2_ FC^b^** | ***p*-value** | **FDR^c^** | **BXD24 CPM^d^** | **CAST CPM^d^** |
| *Mrdq8* | *Soga3* | -0.60 | 7.27E^-4^ | 4.36E^-3^ | 28.70 | 18.90 |
|  | *Hddc2* | 0.78 | 3.54E^-4^ | 2.49E^-3^ | 4.26 | 7.36 |
|  | *Tpd52l1* | 2.69 | 3.90E^-10^ | 4.62E^-8^ | 0.98 | 6.35 |
|  | *Nt5dc1* | 0.88 | 6.42E^-5^ | 6.29E^-4^ | 5.81 | 10.70 |
|  | *Amd2* | 3.96 | 1.30E^-9^ | 1.08E^-7^ | 3.66 | 56.83 |
|  | *Lama4* | -0.56 | 3.10E^-3^ | 1.38E^-2^ | 5.13 | 3.49 |
|  | *Tube1* | -0.95 | 1.08E^-7^ | 3.37E^-6^ | 29.08 | 15.03 |
|  | *E130307A14Rik* | -1.28 | 3.41E^-5^ | 3.80E^-4^ | 11.78 | 4.86 |
|  | *Rev3l* | -0.61 | 9.62E^-3^ | 3.36E^-2^ | 171.65 | 112.54 |
|  | *AA474331* | 0.89 | 1.01E^-4^ | 9.14E^-4^ | 6.72 | 12.47 |
|  | *Amd1* | -0.61 | 3.57E^-3^ | 1.53E^-2^ | 156.89 | 102.70 |
|  | *Ddo* | 1.49 | 1.33E^-7^ | 4.01E^-6^ | 2.12 | 5.93 |
|  | *Mical1* | 0.71 | 2.20E^-3^ | 1.06E^-2^ | 6.90 | 11.29 |
|  | *Lace1* | -0.90 | 1.74E^-6^ | 3.24E^-5^ | 17.26 | 9.24 |
|  | *1700021F05Rik* | 0.57 | 3.78E^-3^ | 1.60E^-2^ | 11.83 | 17.61 |
|  | *Prdm1* | 0.72 | 2.46E^-6^ | 4.36E^-5^ | 21.56 | 35.54 |
|  | *Msl3l2* | 0.82 | 3.65E^-6^ | 5.93E^-5^ | 6.36 | 11.26 |
|  | *Gja1* | 1.23 | 1.13E^-6^ | 2.26E^-5^ | 35.68 | 83.48 |
| *Mrdq9* | *Pisd-ps1* | 2.29 | 2.88E^-7^ | 7.65E^-6^ | 3.56 | 17.21 |
|  | *Sfi1* | 1.71 | 2.83E^-7^ | 7.57E^-6^ | 56.79 | 185.74 |
|  | *8430429K09Rik* | -3.81 | 3.67E^-9^ | 2.39E^-7^ | 5.11 | 0.36 |
|  | *Smtn* | -1.49 | 9.26E^-6^ | 1.29E^-4^ | 8.36 | 3.00 |
|  | *Slc35e4* | 1.07 | 1.98E^-7^ | 5.56E^-6^ | 28.18 | 59.09 |
|  | *Tcn2* | 1.26 | 6.83E^-8^ | 2.4E^-6^ | 39.98 | 95.51 |
|  | *Mtfp1* | -0.54 | 6.70E^-5^ | 6.51E^-4^ | 48.56 | 33.38 |
|  | *Gm11960* | 2.96 | 2.97E^-7^ | 7.82E^-6^ | 0.74 | 5.80 |
|  | *Gm11961* | 0.84 | 3.03E^-5^ | 3.43E^-4^ | 4.10 | 7.34 |
|  | *Thoc5* | 1.81 | 1.33E^-10^ | 1.97E^-8^ | 96.90 | 339.83 |
|  | *Gas2l1* | -0.58 | 1.20E^-3^ | 6.52E^-3^ | 18.38 | 12.26 |
|  | *Rhbdd3* | 0.82 | 1.89E^-4^ | 1.50E^-3^ | 13.88 | 24.48 |
|  | *mmu-miR-3061-3p* | -1.70 | 2.48E^-5^ | N/A | 35.22 | 10.289 |
|  | *mmu-miR-216b-5p* | -0.92 | 3.70E^-2^ | N/A | 51.72 | 26.49 |
|  | *mmu-miR-216a-5p* | -1.26 | 6.57E^-11^ | N/A | 89.74 | 36.77 |
|  | *mmu-miR-217-5p* | -0.83 | 1.44E^-12^ | N/A | 237.01 | 132.18 |
| *Mrdq10* | *Cep112it* | 0.94 | 1.25E^-3^ | 6.76E^-3^ | 16.44 | 30.53 |
|  | *Axin2* | -0.54 | 4.79E^-3^ | 1.93E^-2^ | 41.01 | 28.12 |
|  | *Slc16a6* | -0.95 | 2.18E^-5^ | 2.62E^-4^ | 161.65 | 83.61 |
|  | *Abca8b* | -0.51 | 4.12E^-3^ | 1.71E^-2^ | 32.75 | 22.98 |
|  | *Abca8a* | -0.65 | 2.51E^-4^ | 1.89E^-3^ | 215.20 | 137.37 |
|  | *Abca6* | -2.49 | 7.29E^-7^ | 1.58E^-5^ | 8.48 | 1.49 |
|  | *Fam104a* | 0.68 | 1.18E^-4^ | 1.04E^-3^ | 43.93 | 70.00 |
| **QTL^a^** | **Gene** | **Log_2_ FC^b^** | ***p*-value** | **FDR^c^** | **BXD24 CPM^d^** | **CAST CPM^d^** |
| *Mrdq10* | *D11Wsu47e* | 0.66 | 3.20E^-4^ | 2.31E^-3^ | 14.39 | 22.70 |
|  | *Kif19a* | 1.00 | 2.51E^-6^ | 4.43E^-5^ | 4.85 | 9.71 |
|  | *Slc9a3r1* | 0.62 | 1.26E^-5^ | 1.65E^-4^ | 49.02 | 75.12 |
|  | *Fads6* | -0.59 | 3.68E^-5^ | 4.03E^-4^ | 22.61 | 15.02 |
|  | *Otop3* | -0.81 | 3.85E^-4^ | 2.66E^-3^ | 12.42 | 7.07 |
|  | *Ict1* | 0.52 | 3.65E^-4^ | 2.56E^-3^ | 35.08 | 49.97 |
|  | *Armc7* | 0.56 | 4.18E^-5^ | 4.48E^-4^ | 7.12 | 10.51 |
|  | *Tsen54* | 0.68 | 1.21E^-2^ | 4.01E^-2^ | 7.32 | 11.78 |
|  | *Galk1* | 0.59 | 1.36E^-2^ | 4.36E^-2^ | 10.08 | 15.08 |
|  | *Trim65* | 1.19 | 7.32E^-8^ | 2.54E^-6^ | 4.59 | 10.55 |
|  | *Evpl* | 0.57 | 1.38E^-3^ | 7.30E^-3^ | 3.94 | 5.82 |
|  | *Foxj1* | 0.87 | 3.29E^-3^ | 1.44E^-2^ | 4.40 | 8.03 |
|  | *Prpsap1* | 0.63 | 2.08E^-4^ | 1.62E^-3^ | 44.49 | 68.72 |
|  | *St6galnac2* | 1.10 | 1.40E^-5^ | 1.80E^-4^ | 64.37 | 137.71 |
|  | *Sept9* | 0.55 | 1.63E^-5^ | 2.03E^-4^ | 20.78 | 30.43 |
|  | *Sgsh* | -2.04 | 1.79E^-11^ | 4.62E^-9^ | 62.38 | 15.15 |
|  | *Slc26a11* | -1.22 | 2.39E^-5^ | 2.81E^-4^ | 15.89 | 6.84 |
|  | *Rnf213* | -1.49 | 3.81E^-4^ | 2.64E^-3^ | 14.34 | 5.56 |
|  | *Baiap2* | -0.59 | 3.11E^-3^ | 1.38E^-2^ | 23.65 | 15.69 |
|  | *Aatk* | -0.51 | 1.06E^-4^ | 9.51E^-4^ | 138.79 | 97.11 |
|  | *Faap100* | 0.54 | 6.36E^-3^ | 2.43E^-2^ | 30.16 | 43.91 |
|  | *Tspan10* | 1.24 | 4.34E^-7^ | 1.05E^-5^ | 26.88 | 62.65 |
|  | *Oxld1* | 0.77 | 1.72E^-4^ | 1.39E^-3^ | 5.03 | 8.55 |
|  | *Arl16* | 1.15 | 1.79E^-7^ | 5.12E^-6^ | 12.34 | 27.31 |
|  | *Slc16a3* | -0.51 | 2.42E^-3^ | 1.14E^-2^ | 75.32 | 52.52 |
|  | *Hexdc* | -0.77 | 1.79E^-4^ | 1.43E^-3^ | 92.08 | 54.17 |
|  | *Fn3krp* | -0.82 | 2.21E^-7^ | 6.14E^-6^ | 33.84 | 19.18 |
|  | *B3gntl1* | 1.09 | 4.25E^-7^ | 1.03E^-5^ | 5.79 | 12.31 |
| *Mrdq11* | *Rps6-ps4* | -9.36 | 4.42E^-10^ | 5.15E^-8^ | 302.83 | 0.47 |
|  | *Gm13394* | -8.88 | 1.48E^-7^ | 4.40E^-6^ | 3193.43 | 9.96 |
|  | *A730017L22Rik* | -5.69 | 1.99E^-13^ | 2.83E^-10^ | 10.00 | 0.19 |
|  | *5330413P13Rik* | -5.18 | 2.76E^-10^ | 3.57E^-8^ | 6.64 | 0.18 |
|  | *Mrpl23-ps1* | -4.84 | 3.04E^-6^ | 5.14E^-5^ | 17.72 | 0.61 |
|  | *Spc25* | -2.59 | 1.41E^-10^ | 2.00E^-8^ | 138.60 | 22.96 |
|  | *Upp2* | -2.00 | 1.19E^-9^ | 1.02E^-7^ | 5.68 | 1.42 |
|  | *Cerkl* | -1.91 | 5.25E^-9^ | 3.10E^-7^ | 92.56 | 24.49 |
|  | *Col5a1* | -1.83 | 3.40E^-6^ | 5.61E^-5^ | 12.17 | 3.32 |
|  | *Proser2* | -1.82 | 1.16E^-7^ | 3.56E^-6^ | 11.87 | 3.37 |
|  | *Gm9821* | -1.81 | 1.35E^-5^ | 1.75E^-4^ | 9.32 | 2.64 |
|  | *Cd93* | -1.74 | 1.53E^-5^ | 1.94E^-4^ | 8.54 | 2.63 |
|  | *Pla2r1* | -1.67 | 2.29E^-5^ | 2.73E^-4^ | 110.36 | 34.74 |
|  | *Tgm2* | -1.67 | 3.12E^-9^ | 2.16E^-7^ | 12.43 | 3.93 |
| **QTL^a^** | **Gene** | **Log_2_ FC^b^** | ***p*-value** | **FDR^c^** | **BXD24 CPM^d^** | **CAST CPM^d^** |
| *Mrdq11* | *Gm13583* | -1.64 | 1.44E^-6^ | 2.77E^-5^ | 5.53 | 1.78 |
|  | *Metap1d* | -1.63 | 1.65E^-7^ | 4.80E^-6^ | 47.98 | 15.50 |
|  | *Egfl7* | -1.48 | 3.06E^-7^ | 7.95E^-6^ | 8.53 | 3.05 |
|  | *Stom* | -1.42 | 1.01E^-10^ | 1.63E^-8^ | 98.06 | 36.60 |
|  | *Ryr3* | -1.35 | 3.32E^-6^ | 5.50E^-5^ | 30.67 | 11.71 |
|  | *Itga4* | -1.35 | 4.08E^-6^ | 6.50E^-5^ | 136.95 | 52.37 |
|  | *Hmcn2* | -1.24 | 1.07E^-4^ | 9.56E^-4^ | 10.23 | 4.37 |
|  | *Cd59a* | -1.19 | 2.54E^-7^ | 6.91E^-6^ | 25.39 | 11.13 |
|  | *Ccdc173* | -1.17 | 8.25E^-7^ | 1.75E^-5^ | 13.66 | 6.05 |
|  | *Rexo4* | -1.11 | 2.72E^-3^ | 1.25E^-2^ | 27.60 | 13.51 |
|  | *Csrnp3* | -1.11 | 2.46E^-5^ | 2.87E^-4^ | 62.09 | 28.52 |
|  | *Fbn1* | -1.09 | 1.39E^-5^ | 1.79E^-4^ | 39.24 | 18.57 |
|  | *Necab3* | -1.06 | 1.97E^-7^ | 5.55E^-6^ | 9.67 | 4.65 |
|  | *Itga6* | -1.03 | 1.77E^-5^ | 2.17E^-4^ | 11.77 | 5.81 |
|  | *Gm14204* | -1.01 | 6.38E^-8^ | 2.29E^-6^ | 28.64 | 14.17 |
|  | *Thbs1* | -0.96 | 4.90E^-5^ | 5.08E^-4^ | 111.52 | 57.48 |
|  | *Snhg17* | -0.95 | 9.01E^-5^ | 8.27E^-4^ | 8.65 | 4.46 |
|  | *Adamtsl2* | -0.94 | 3.55E^-6^ | 5.81E^-5^ | 7.80 | 4.07 |
|  | *Cdc25b* | -0.89 | 7.89E^-6^ | 1.14E^-4^ | 7.51 | 4.04 |
|  | *4930405A21Rik* | -0.89 | 2.56E^-4^ | 1.92E^-3^ | 7.55 | 4.10 |
|  | *Fam171b* | -0.87 | 3.97E^-6^ | 6.32E^-5^ | 334.77 | 183.01 |
|  | *Soga1* | -0.83 | 1.18E^-2^ | 3.94E^-2^ | 34.96 | 19.49 |
|  | *Lrp1b* | -0.82 | 1.26E^-2^ | 4.13E^-2^ | 17.33 | 9.70 |
|  | *Cd44* | -0.79 | 4.16E^-4^ | 2.82E^-3^ | 29.04 | 16.81 |
|  | *Myl9* | -0.76 | 4.39E^-4^ | 2.94E^-3^ | 7.95 | 4.68 |
|  | *Rif1* | -0.75 | 2.55E^-3^ | 1.19E^-2^ | 87.06 | 51.76 |
|  | *Eng* | -0.75 | 1.88E^-4^ | 1.50E^-3^ | 10.61 | 6.32 |
|  | *Nebl* | -0.74 | 4.38E^-4^ | 2.93E^-3^ | 75.97 | 45.07 |
|  | *Ehd4* | -0.72 | 3.99E^-7^ | 9.77E^-6^ | 48.61 | 29.52 |
|  | *Sfmbt2* | -0.72 | 1.57E^-3^ | 8.11E^-3^ | 6.91 | 4.16 |
|  | *A430105I19Rik* | -0.72 | 5.79E^-4^ | 3.64E^-3^ | 7.96 | 4.83 |
|  | *Thnsl1* | -0.71 | 3.60E^-3^ | 1.55E^-2^ | 17.60 | 10.73 |
|  | *Gm996* | -0.71 | 4.36E^-5^ | 4.63E^-4^ | 40.40 | 24.66 |
|  | *Chgb* | -0.71 | 1.50E^-3^ | 7.79E^-3^ | 503.36 | 307.84 |
|  | *Snhg11* | -0.69 | 2.35E^-3^ | 1.11E^-2^ | 412.98 | 257.86 |
|  | *Nr6a1* | -0.69 | 2.22E^-5^ | 2.66E^-4^ | 21.83 | 13.52 |
|  | *Cutal* | -0.67 | 3.41E^-5^ | 3.80E^-4^ | 9.07 | 5.72 |
|  | *Abtb2* | -0.67 | 5.37E^-4^ | 3.41E^-3^ | 7.46 | 4.70 |
|  | *Pdk1* | -0.66 | 2.10E^-4^ | 1.63E^-3^ | 59.53 | 37.61 |
|  | *Sptan1* | -0.66 | 6.66E^-5^ | 6.48E^-4^ | 964.53 | 611.98 |
|  | *Phpt1* | -0.63 | 4.57E^-5^ | 4.81E^-4^ | 53.34 | 34.19 |
|  | *Slc1a2* | -0.63 | 3.93E^-6^ | 6.27E^-5^ | 403.37 | 259.64 |
|  | *Klhl23* | -0.63 | 1.12E^-4^ | 9.93E^-4^ | 36.45 | 23.49 |
| **QTL^a^** | **Gene** | **Log_2_ FC^b^** | ***p*-value** | **FDR^c^** | **BXD24 CPM^d^** | **CAST CPM^d^** |
| *Mrdq11* | *Phf21a* | -0.63 | 3.35E^-4^ | 2.38E^-3^ | 56.94 | 36.67 |
|  | *Rc3h2* | -0.63 | 1.39E^-3^ | 7.32E^-3^ | 73.76 | 47.38 |
|  | *Rasgrp1* | -0.62 | 1.39E^-4^ | 1.19E^-3^ | 36.72 | 23.90 |
|  | *Kcnt1* | -0.62 | 1.15E^-3^ | 6.34E^-3^ | 23.08 | 14.89 |
|  | *Ttn* | -0.61 | 1.12E^-3^ | 6.17E^-3^ | 9.79 | 6.41 |
|  | *Accs* | -0.61 | 3.17E^-4^ | 2.29E^-3^ | 5.01 | 3.29 |
|  | *Pdyn* | -0.60 | 1.61E^-2^ | 4.95E^-2^ | 10.65 | 7.00 |
|  | *Madd* | -0.60 | 9.77E^-5^ | 8.88E^-4^ | 573.35 | 377.45 |
|  | *Kcnj3* | -0.58 | 2.40E^-3^ | 1.13E^-2^ | 17.12 | 11.36 |
|  | *Kcnh7* | -0.56 | 8.23E^-3^ | 2.98E^-2^ | 23.63 | 15.60 |
|  | *Edem2* | -0.55 | 1.47E^-3^ | 7.67E^-3^ | 46.47 | 31.79 |
|  | *Scrn3* | -0.54 | 1.20E^-4^ | 1.05E^-3^ | 38.50 | 26.47 |
|  | *Pfkfb3* | -0.53 | 6.19E^-5^ | 6.11E^-4^ | 42.33 | 29.31 |
|  | *Sardh* | -0.53 | 3.88E^-3^ | 1.63E^-2^ | 8.00 | 5.61 |
|  | *Hspa12b* | -0.52 | 1.87E^-3^ | 9.30E^-3^ | 5.05 | 3.51 |
|  | *Kif3b* | -0.52 | 5.82E^-5^ | 5.82E^-4^ | 158.20 | 110.69 |
|  | *Osbpl6* | -0.51 | 6.04E^-4^ | 3.76E^-3^ | 59.67 | 41.59 |
|  | *Nfe2l2* | 0.51 | 4.94E^-4^ | 3.21E^-3^ | 25.69 | 36.59 |
|  | *Mrps26* | 0.51 | 3.88E^-4^ | 2.68E^-3^ | 30.34 | 43.04 |
|  | *Gpr155* | 0.52 | 2.24E^-3^ | 1.07E^-2^ | 25.96 | 37.05 |
|  | *Gm13736* | 0.53 | 4.82E^-3^ | 1.93E^-2^ | 29.94 | 42.93 |
|  | *Ccbl1* | 0.55 | 1.28E^-4^ | 1.11E^-3^ | 21.49 | 31.49 |
|  | *Rbbp9* | 0.56 | 4.16E^-5^ | 4.46E^-4^ | 26.08 | 38.51 |
|  | *Pnpla7* | 0.57 | 1.48E^-4^ | 1.24E^-3^ | 12.39 | 18.33 |
|  | *Gorasp2* | 0.57 | 4.68E^-3^ | 1.89E^-2^ | 131.79 | 196.10 |
|  | *Tprn* | 0.57 | 8.03E^-5^ | 7.49E^-4^ | 9.35 | 13.91 |
|  | *Bmp2* | 0.59 | 4.78E^-5^ | 4.98E^-4^ | 5.95 | 8.92 |
|  | *Ntng2* | 0.59 | 2.00E^-5^ | 2.42E^-4^ | 77.20 | 116.06 |
|  | *A230005M16Rik* | 0.60 | 4.12E^-3^ | 1.71E^-2^ | 4.25 | 6.40 |
|  | *Serf2* | 0.60 | 3.67E^-4^ | 2.56E^-3^ | 171.55 | 259.24 |
|  | *Eid1* | 0.60 | 1.53E^-3^ | 7.92E^-3^ | 122.14 | 185.10 |
|  | *Eif6* | 0.60 | 5.04E^-3^ | 2.01E^-2^ | 31.22 | 47.19 |
|  | *Stard9* | 0.61 | 8.60E^-3^ | 3.09E^-2^ | 13.83 | 19.71 |
|  | *Trp53i11* | 0.61 | 6.97E^-5^ | 6.72E^-4^ | 15.87 | 24.21 |
|  | *Pak6* | 0.61 | 4.79E^-4^ | 3.13E^-3^ | 3.47 | 5.30 |
|  | *Rprm* | 0.61 | 2.18E^-4^ | 1.68E^-3^ | 4.91 | 7.45 |
|  | *C1qtnf4* | 0.62 | 8.47E^-4^ | 4.93E^-3^ | 14.73 | 22.26 |
|  | *Scand1* | 0.63 | 2.37E^-3^ | 1.12E^-2^ | 16.31 | 24.82 |
|  | *Dtd1* | 0.63 | 4.25E^-5^ | 4.53E^-4^ | 18.81 | 29.04 |
|  | *Cox4i2* | 0.64 | 1.00E^-3^ | 5.65E^-3^ | 17.39 | 26.74 |
|  | *Galk2* | 0.64 | 7.81E^-5^ | 7.31E^-4^ | 19.75 | 30.60 |
|  | *Haus2* | 0.64 | 2.75E^-3^ | 1.26E^-2^ | 13.54 | 21.15 |
| **QTL^a^** | **Gene** | **Log_2_ FC^b^** | ***p*-value** | **FDR^c^** | **BXD24 CPM^d^** | **CAST CPM^d^** |
| *Mrdq11* | *Fibcd1* | 0.65 | 3.43E^-5^ | 3.81E^-4^ | 16.02 | 25.09 |
|  | *Snrpb* | 0.65 | 5.37E^-3^ | 2.12E^-2^ | 104.94 | 164.23 |
|  | *Fap* | 0.66 | 7.68E^-5^ | 7.22E^-4^ | 17.51 | 27.60 |
|  | *Gca* | 0.66 | 1.66E^-4^ | 1.36E^-3^ | 15.34 | 24.11 |
|  | *Timm10* | 0.67 | 1.24E^-2^ | 4.07E^-2^ | 6.42 | 10.16 |
|  | *Rnf208* | 0.68 | 2.16E^-3^ | 1.05E^-2^ | 28.87 | 46.25 |
|  | *Wdr34* | 0.71 | 1.94E^-3^ | 9.58E^-3^ | 23.88 | 38.94 |
|  | *Nelfb* | 0.71 | 2.72E^-5^ | 3.15E^-4^ | 42.67 | 69.87 |
|  | *Hnmt* | 0.72 | 4.14E^-4^ | 2.81E^-3^ | 8.64 | 14.17 |
|  | *Ptgds* | 0.73 | 1.61E^-5^ | 2.02E^-4^ | 650.23 | 1067.06 |
|  | *Pla2g4e* | 0.78 | 4.94E^-6^ | 7.67E^-5^ | 9.35 | 15.99 |
|  | *Fbln7* | 0.79 | 6.60E^-5^ | 6.43E^-4^ | 3.52 | 6.07 |
|  | *Trub2* | 0.79 | 2.34E^-6^ | 4.19E^-5^ | 27.92 | 48.36 |
|  | *Sord* | 0.80 | 7.72E^-7^ | 1.66E^-5^ | 13.21 | 22.93 |
|  | *Id1* | 0.80 | 4.66E^-4^ | 3.06E^-3^ | 5.61 | 9.67 |
|  | *Msrb2* | 0.82 | 1.38E^-4^ | 1.18E^-3^ | 5.16 | 9.08 |
|  | *Cd82* | 0.83 | 1.09E^-6^ | 2.20E^-5^ | 19.00 | 33.68 |
|  | *Bub1b* | 0.83 | 6.53E^-4^ | 4.01E^-3^ | 5.01 | 8.94 |
|  | *Lmo2* | 0.84 | 1.93E^-6^ | 3.55E^-5^ | 12.91 | 23.01 |
|  | *Casc4* | 0.86 | 1.81E^-4^ | 1.45E^-3^ | 39.82 | 71.62 |
|  | *Spef1* | 0.87 | 2.22E^-4^ | 1.70E^-3^ | 11.49 | 20.96 |
|  | *2500004C02Rik* | 0.90 | 9.62E^-6^ | 1.32E^-4^ | 5.86 | 10.94 |
|  | *Lrrc57* | 0.93 | 4.19E^-7^ | 1.02E^-5^ | 21.99 | 41.86 |
|  | *Dync1i2* | 0.97 | 1.11E^-8^ | 5.70E^-7^ | 102.73 | 201.77 |
|  | *Cst3* | 0.98 | 1.13E^-5^ | 1.52E^-4^ | 1132.06 | 2210.12 |
|  | *Nek6* | 1.03 | 3.10E^-5^ | 3.50E^-4^ | 17.89 | 36.59 |
|  | *Chchd5* | 1.03 | 2.71E^-7^ | 7.29E^-6^ | 13.11 | 26.71 |
|  | *Elp4* | 1.04 | 6.32E^-6^ | 9.43E^-5^ | 26.16 | 53.75 |
|  | *D330023K18Rik* | 1.07 | 5.52E^-6^ | 8.45E^-5^ | 2.86 | 6.01 |
|  | *Cd302* | 1.09 | 6.99E^-5^ | 6.72E^-4^ | 2.62 | 5.55 |
|  | *Gsn* | 1.38 | 2.64E^-7^ | 7.14E^-6^ | 4.61 | 11.99 |
|  | *A530058N18Rik* | 1.48 | 1.84E^-7^ | 5.23E^-6^ | 15.12 | 41.61 |
|  | *Entpd2* | 1.86 | 5.20E^-9^ | 3.08E^-7^ | 1.70 | 6.13 |
|  | *Bfsp1* | 2.24 | 1.79E^-3^ | 8.99E^-3^ | 5.29 | 23.01 |
|  | *Stamos* | 2.42 | 7.77E^-9^ | 4.19E^-7^ | 0.97 | 5.21 |
|  | *Gm14150* | 2.78 | 3.55E^-6^ | 5.81E^-5^ | 16.84 | 113.85 |
|  | *Slc27a2* | 2.92 | 3.62E^-10^ | 4.37E^-8^ | 1.16 | 8.80 |
|  | *Hdc* | 3.70 | 4.88E^-11^ | 1.07E^-8^ | 0.62 | 8.09 |
|  | *Gm14494* | 3.86 | 3.03E^-6^ | 5.13E^-5^ | 1.85 | 25.86 |
|  | *A930006I01Rik* | 5.34 | 3.25E^-13^ | 4.15E^-10^ | 0.51 | 20.79 |
|  | *Gm13339* | 10.06 | 4.87E^-7^ | 1.16E^-5^ | 0.16 | 175.97 |
|  | *Gm13340* | 10.10 | 8.37E^-9^ | 4.44E^-7^ | 4.69 | 4579.38 |
|  | *Gm13341* | 10.16 | 2.06E^-7^ | 5.75E^-6^ | 2.43 | 2490.06 |
| **QTL^a^** | **Gene** | **Log_2_ FC^b^** | ***p*-value** | **FDR^c^** | **BXD24 CPM^d^** | **CAST CPM^d^** |
| *Mrdq11* | *Gm13772* | 10.46 | 6.25E^-6^ | 9.34E^-5^ | 0.00 | 11.02 |
|  | *mmu-miR-467c-5p* | -4.06 | 5.19E^-7^ | N/A | 9.63 | 0.00 |
|  | *mmu-miR-467a-5p* | -1.48 | 1.01E^-2^ | N/A | 17.16 | 5.69 |
|  | *mmu-miR-10b-5p* | -2.77 | 3.83E^-7^ | N/A | 89.80 | 10.49 |
|  | *mmu-miR-670-3p* | -1.55 | 5.24E^-3^ | N/A | 21.45 | 6.92 |

^a^ QTL, quantitative trait locus

^b^ FC, fold change > 0.5 or < -0.5

^c^ FDR, false discovery rate

^d^ CPM, counts per million
